# Hypergraph Cortical Cytoarchitectonic Parcellation with Multimodal Canine Brain Atlas

**DOI:** 10.1101/2024.05.09.593154

**Authors:** Shiyao Zhai, Chenyao Jiang, Zhenglin Chen, Dongmei Yu, Ijaz Gul, Xi Yuan, Hengrui Song, Yangliu, Ziheng Zhang, Tao Xu, Hu Xu, Jiusheng Wan, Aiguo Mao, Jie Li, Yuxing Han, Peiwu Qin

**Affiliations:** Center of Precision Medicine and Healthcare, Tsinghua-Berkeley Shenzhen Institute, Shenzhen, Guangdong Province, 518055, China; Institute of Biopharmaceutical and Health Engineering, Tsinghua Shenzhen International Graduate School, Shenzhen, Guangdong Province, 518055, China; Division of Information Science and Technology, Tsinghua Shenzhen International Graduate School, Shenzhen, Guangdong Province, 518055, China; Kunming Police Dog Base of the Ministry of Public Security, Kunming, Yunnan Province, 650204, China; MOE Key Lab of Industrial Biocatalysis, Department of Chemical Engineering, Tsinghua University, Beijing, 100084, China; School of Mechanical, Electrical & Information Engineering, Shandong University, Weihai, Shandong, 264209, China

**Author notes:** Corresponding Author Peiwu Qin,., Yuxing Han, Jie Li. Marking the co-first authors of this article (Shiyao Zhai, Chenyao Jiang).

## Abstract

Brain atlases are vital tools in exploring the brain structure-function relationship. The burgeoning cross-species atlases have significantly accelerated our understanding of human brain development, evolution, function, and diseases. However, the existing coarse-grained macroscopic canine brain atlases greatly constrain their utility as an animal model for neurocognition research. Finer-grained brain atlas and partitions are crucial for decoding brain spatial heterogeneity and topology at different scales. Therefore, we conduct macroscopic and microscopic brain imaging to construct an interactive online dataset of multimodal canine brain atlas. Additionally, we develop a pioneering method for cortical cytoarchitectonic partitioning based on hypergraph learning. By integrating high-dimensional cytoarchitectonic features and spatial connections between cortical columns, the method leads to fine-grained partitioning patterns. This innovative approach aims to decode the biological heterogeneity of cortical microstructures, contributing to the structural annotation of canine atlas as well as public human brain atlases. The study not only offers valuable resources but also presents a novel zonation approach to investigate the cellular organization pattern and topology of the cortex.

## Introduction

Brain atlases serve as navigational maps to explore the spatial distribution and local microscopic structure of diverse regions and elucidate the intricate relationship between brain structure and function. In addition, the brain atlas guides diverse fields including clinical diagnosis, therapeutic intervention, artificial intelligence, education, and psychology. Since Brodmann’s cortical atlas and manual partition emerged in 1909 ^1^, brain atlases have been updated continually as technological progress ^2^. In terms of atlas content, cross-scale, multiple-spices and multimodal database provide data support and interpretation for brain parcellation. The multi-scale integration of multimodal information occurs at the level of cell, microcircuits, columnar zone, cortex, to brain, where multiscale segmentation is the automatic identification of these anatomical features, critical for establishing connections between brain regions and their functions ^3^. Probabilistic maps of cellular architecture indicate correlations between microstructures and cognitive functions ^4^. The platform of brain atlases has evolved from physical hand-drawn image to digital atlases with interactivity and functionality, offering visualization possibilities for multimodal, cross-scale atlas ^5^ ^6^.

Multimodal brain atlases have been available for multiple species during the last decades^7^ ^8^ ^9^. However, progress in canine brain atlases seems stagnant, although which shares common diseases, medical treatments, and living environments with humans as a promising animal model to study aging, neurocognition, and neurological diseases ^10^. Existing canine brain atlases are limited to single imaging modalities and macroscopic scales ^11^ ^12^ ^13^, becoming a bottleneck for in-depth neuroscience and clinical study. The low resolution of canine brain atlases and coarse granularity of parcellation restricts the projection between function and structure, identification of intraregional neural circuits, and pathological comparison. Thus, effective microstructural partitioning of the canine cerebral cortex is unavailable. Refined cortical parcellations establish a connection between structure and function ^14^, and cytoarchitectonics plays an indispensable role in brain parcellation. Specific neurons with different anatomical and physiological properties formulate the distinct neural network visualized through the multimodal brain atlas. Understanding brain function is enhanced through the description of cellular organization patterns and cortical parcellation based on cytoarchitectonic heterogeneity ^15^.

Cytoarchitecture encompasses neuron morphology, relative spatial orientation, cellular composition, and organization pattern, serving as a fundamental principle for brain microstructural zonation ^4^. Cytoarchitecture serves as an interface and reference for integrating various sources of the brain data including connection patterns, myelin density, enzymatic activity, and gene expression ^16^. The cytoarchitectonic differences across various regions reveal the different distribution of specific types of neurons, closely associated with distinct gene expression and electrophysiological traits. Moreover, cellular morphology and extension patterns represent different neuronal connection pathways at specific brain regions, indicating a cortical function differentiation across brain areas. The cerebral cortex can be divided into columnar regions via cytoarchitectonic lamination patterns ^17^. Physiological and imaging studies have shown different neuronal responses between two columnar areas. Microscopic analysis of brain tissue sections remains the "gold standard" for cytoarchitectonic parcellation ^4^. However, the explosion of high-throughput and super-resolution imaging presents a challenge for visual inspections. An observer-independent technique for pinpointing cytoarchitectonic boundaries, the Gray Level Index (GLI) method, has been used in human cortical partitioning, offering a quantification of cytoarchitecture with exceeding granularity ^4^ ^18^. However, the abstraction of intraregional cytoarchitecture inevitably overlooks the structural nuances and interregional connections, reaching a bottleneck for GLI in the finer-grained parcellation of cortical microstructure.

To overcome the scarcity of the multimodal canine atlas and reliable parcellation method, we create a multimodal, cross-scale, brain atlas encompassing MRI and multiple staining modalities (image of gigapixel with pixel resolution of 0.5 μm). The size of sample slide is 76 x 52 mm, corresponding to 152 k x 104 k or ∼1.58 x 10^10^ pixels in total. If half slide is occupied with brain slice, we have ∼7 x 10^9^ pixels per whole slide imaging. We develop an alternative cortical partitioning method with hypergraph learning that aids in delineating cytoarchitectonics complementary to GLI partition (Fig.1). A hypergraph is constructed, with each vertex representing a single cortical columnar unit. The feature vector of vertex incorporates 49 distinct cytoarchitectural features computed from the neuron architecture at different cortex layers (Table S1). We validate the generalization of hypergraph zonation using publicly available human brain atlases and our newly created canine brain atlas. The partitions, integrating myeloarchitectonics, neuroanatomy, and neuroarchitecture at macroscopic and microscopic scales, are demonstrated. Finally, we integrate the original multimodal data with the zonation maps into an openly accessible, interactive web-based database.

**Figure 1.**
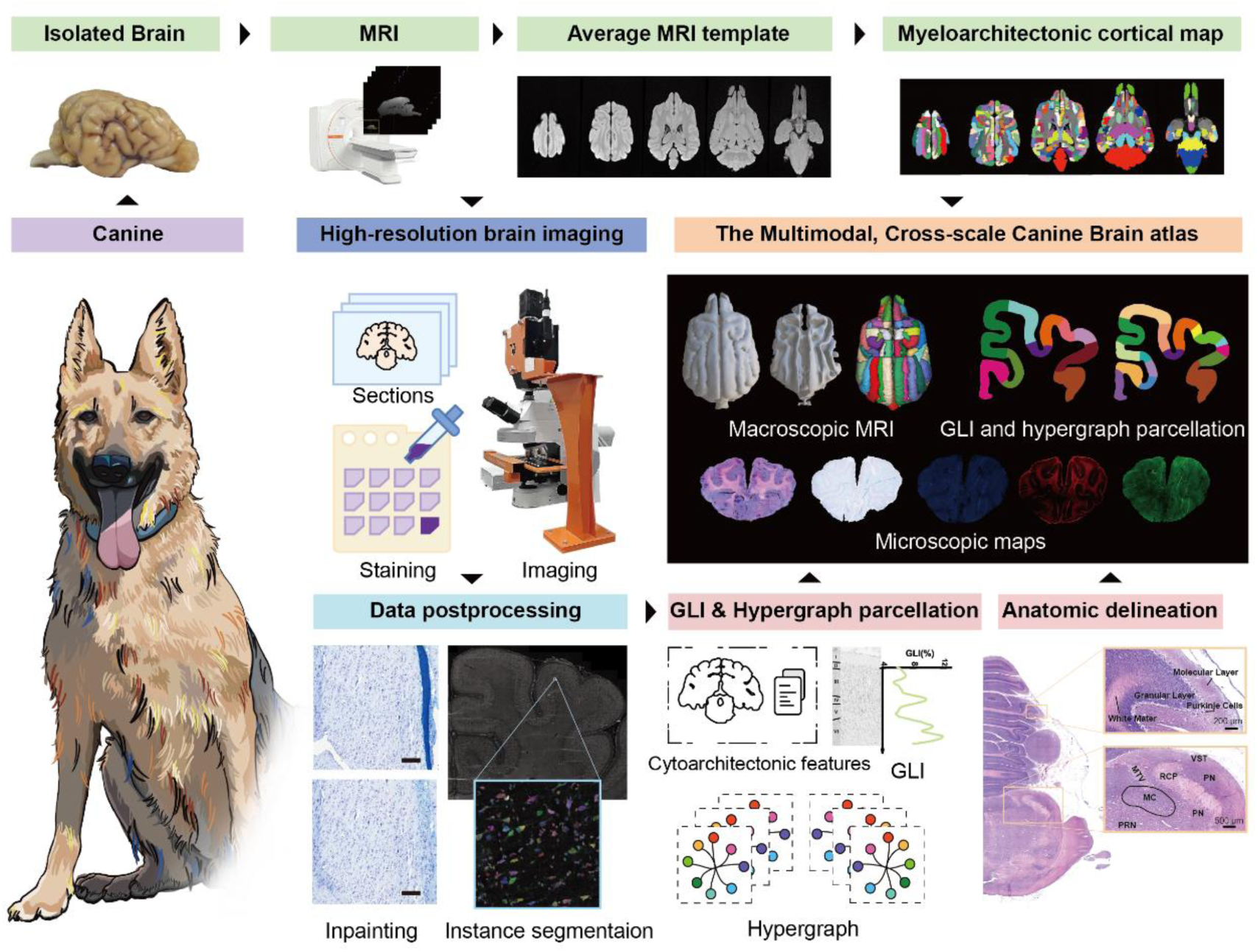
Workflow for the construction of the multimodal canine brain atlas. The brains are imaged by the T1W MRI, and aligned to generate an average brain template in MNI space after noise reduction. The voxel size is 0.5 mm^3^. We can manually delineate gray and white matter distribution and myeloarchitectonic cortical zonation. Sectioned tissue specimens undergo a series of treatments, including pre-processing, sectioning, staining, and imaging, to acquire coronal brain slice images. The images undergo post-processing steps including the detection and repairment of artifacts and the cellular instance segmentation. Instance segmentation facilitates the extraction and computation of cytoarchitecture for all cells in the image. Cellular features serve as inputs for the GLI and the hypergraph learning for the cytoarchitectonic cortical parcellation. The interactive canine brain atlas is created by integrating macroscopic MRI, high-resolution microstructural brain images with anatomical annotation, and the results of cytoarchitectonic parcellation.

## Results

### Canine brain’s MRI Template and its Morphology

We depict the morphology of the Kunming canine brain acquired with smartphone in Fig. 2A, and Table S2 contains the name list of all the sulci, gyrus, and subcortical regions. We create multiple templates including the average brain template (Fig. 2B), the gray matter (GM) and white matter (WM) templates at 0.5 * 0.5 * 0.5 mm/voxel resolution (Fig. 2C), and the cortical and subcortical parcellation template (Fig. 2D). Through the myeloarchitectonic and the anatomical delineation ^13^, the cortex and the subcortex in MRI are parcellated into 212 and 12 regions, respectively. The complete parcellations and zone names with labels are illustrated in Fig. S1, and their statistical volumes are provided in the Table S3. Fig. 2E shows the three-dimensional (3D) reconstruction of different brain maps.

**Figure 2.**
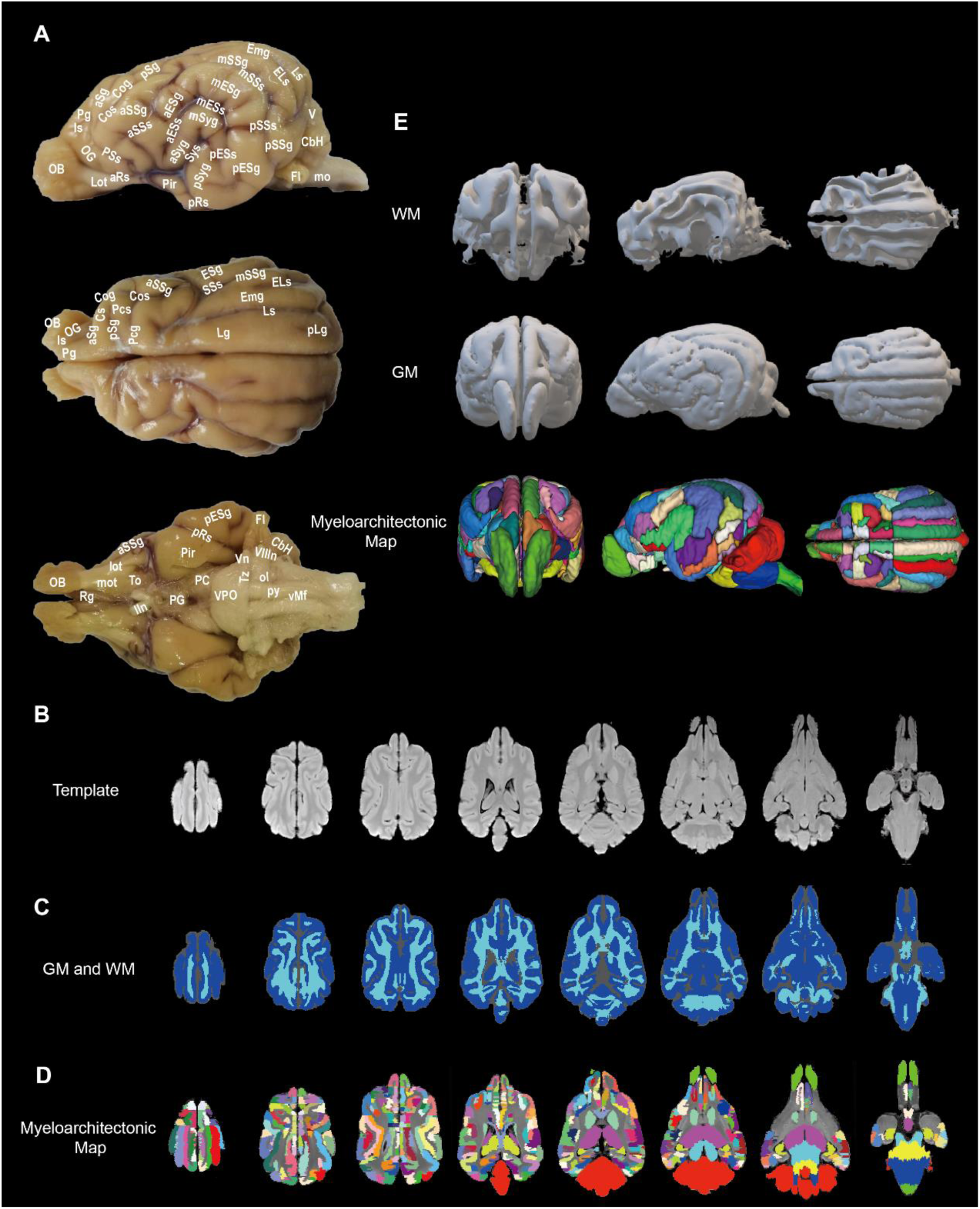
The morphology and MRI template of Kunming canine Brain. (A) shows the sulci, the gyri, and the subcortex structure through the lateral, vertical, and bottom view. The full names of these abbreviations are provided in the Table S2. (B) The MRI template averaged from four *ex vivo* samples imaged with T1W sequence at 0.5 * 0.5 * 0.5 mm/voxel resolution. (C) Axial slices show the template of white and gray matter segmentation (WM, light blue; GM, blue). (D) Cortex and subcortex parcellation based on the T1W template. (E) The 3D models of segmented GM, WM, and myeloarchitectonic maps with zone labels.

### Brain Atlas with Different Histological Staining

We cryosection the brain into 20-μm-thick coronal slices in groups spaced at 500 μm in a rostral-to-caudal direction. Each group contains ten consecutive slices and a total of 800 slices are selected for Hematoxylin and Eosin (H&E) staining (Fig. 3A). Then, we pick 200 coronal sections for the Klüver-Barrera (KB) staining. All Nuclei and neuronal nissl bodies appear blue-purple, while myelin fibers appear light blue and exhibit a darker shade as fiber density increases (Fig. 3B). Subsequently, we select 80 slices for triple immunofluorescence staining with Hoechst, anti-PSD-95, and anti-GFAP simultaneously. Immunofluorescence imaging of whole coronal slices has five focal planes per 3D volume and three-colors with a resolution of 500 nm/pixel (Fig. 3C). The scaffolding protein on the postsynaptic membrane, postsynaptic density protein 95 (PSD-95), primarily exists in glutamatergic neurons where it is translated and transported within the neural cytosol to participate the formation of excitatory synapses. Thus, Hoechst-stained nuclei appear blue (Fig. 3D) and the cytosol and dendrites of glutamatergic neurons exhibit a green color due to PSD-95 staining (Fig.3E) in confocal images. Moreover, glial fibrillary acidic protein (GFAP) staining with red probes to make astrocytes as red color in merged image (Fig. 3F). The H&E stained images with cellular resolution facilitate the delineations of microscopic neuroanatomy for individual slices shown in Supplementary Figures S2-16. We can visualize certain brain regions with the naked eye such as the molecular layer, granular layer, and white matter of cerebellum in the H&E staining and the distribution of certain cells such as Purkinje cells easily (Fig. 3J). The cellular morphology, density, and orientation of neural fiber in H&E facilitate the identification of white matter tracts and brain nuclei in the pons (Fig. 3K). The boundary of the rostral cerebellar peduncle (RCP) is clearly distinguished from the parabrachial nuclei (PN). We can trace the orientation of ventral spinocerebellar tract (VST) in pink color and find the aggregation of bigger neurons in the micturition center. Astrocytes show a heterogeneous distribution between gray and white matter in GFAP-stained images (Fig. 3L). The brain cortex has 6 separate layers stacked vertically and the distribution of astrocytes in different layers is seldom reported. Although it is impossible to identify the cortex layers based on astrocytes, the astrocytes adjacently connected to the pia mater help determine the layer I boundary, and the astrocytes become progressively denser from the layer V to VI. The density of astrocytes in the white matter is higher, supporting oligodendrocytes and myelination. In addition, GFAP staining helps confirm the anatomical delineation of the subcortical gray matter and white matter, such as the corpus callosum, caudate nucleus, thalamic reticulate nucleus, internal capsule, and claustrum (Fig. 3M). KB staining shows the cortical laminar cytoarchitecture (Fig. 3N), where layer I has sparse cell density with scattering distribution of horizontal cells of Cajal-Retzius and astrocytes. Although layer II and layer III have continuous transition with vague boundary, we can visualize the external granular layer (layer II) consisting of dense small granular cells. Compared to layer II, layer III is identifiable due to larger and deeply stained pyramidal cells. The layer IV is narrow and consists mostly of the granular cells and a smaller fraction of the pyramidal cells. The pyramidal cells of layer V are smaller or similar than those from layer III in the ROIs of the visual cortex, while the pyramidal cells of layer V are larger in motor cortex. Layer VI has a smooth transition to the white matter, which contains blue-purple nuclei and light blue myelin fibers (Fig. 3N). The cortical band of comu ammonis (CA) has four regions according to bandwidth, cell size, and cell density (Fig. 3O). CA1 contains smaller pyramidal cells, while CA2 is a narrow and dense band with larger pyramidal cells. CA3 is a broad and loose band of larger pyramidal neurons, while CA4 forms the loosely-structured end zone (Fig. 3O). CA is enclosed by the narrow and dark band of neural cells from the dentate gyrus. The subiculum is the transitional region between the CA1 and the adjacent entorhinal cortex, which contains the larger and relatively loosely organized pyramidal. The stratum oriens of CA1 terminates into the subiculum.

**Figure 3.**
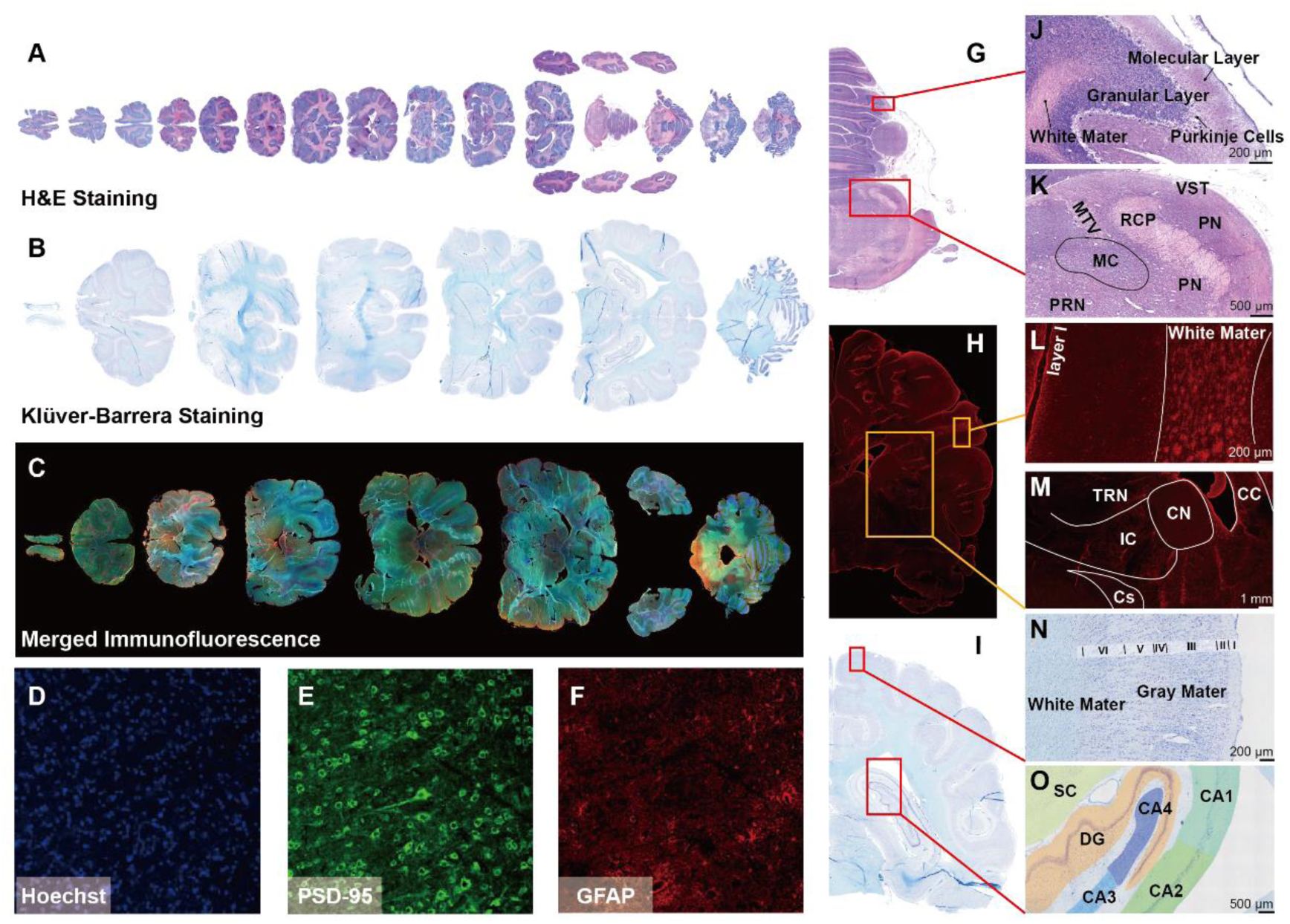
Brain atlas of high-resolution slices with various staining methods. Examples of brain coronal histological images in the rostral to caudal direction with H&E staining (A), KB staining (B), and immunofluorescence (C). (D), (E) and (F) show immunofluorescent images under different illuminating channels, with nuclei in blue, PSD-95 in green, and GFAP in red. (G) is an example of a whole-hemisphere section of the cerebellum and pons with H&E staining, and (J) and (K) are manual zone parcellation in square ROIs (J) shows cerebellar cortex and white matter fiber tracts, and (K) shows subcortical structures. (H) is an image of GFAP staining. (L) and (M) are anatomical annotations for the cortex and subcortex. (I) is a section with KB staining and (N) shows the laminar delineation of cortical layers I – VI. (O) labels hippocampus subregions in different colors. Abbreviations VST – ventral spinocerebellar tract; PN – parabrachial nuclei; RCP – rostral cerebellar peduncle; MTV – mesencephalic tract of V; MC – micturition center; PRN – pontine reticular nucleus; CC – corpus callosum; CN – caudate nucleus; TRN – thalamic reticulate nucleus; IC – internal capsule; Cs – claustrum. CA, comu ammonis; DG, dentate gyrus; SC, subiculum.

### Cell Instance Segmentation and Inpainting of Histological Images

During the cryosectioning and the staining, artifacts, such as distortion, folding, tearing, and inhomogeneous staining are unavoidable (Fig. 4A-C). The panoramic images and expanded insets with different staining indicate the artifacts and outcomes of digital repair: faded spots, balanced uneven stains, patched broken areas, and replaced wrinkles, highlighting the feasibility and generalizability of the repair algorithm. Furthermore, we employ the Cellpose for cell instance segmentation in various digital sections ^19^. We conduct grayscale pre-processing for all sections. Fig4. M-O showcases the nuclei instance segmentation for the whole and zoomed area of H&E sections. Fig4. P-R shows the instance segmentation of the nucleus and neuronal cytosol within the ROI for the KB-stained section, enabling the extraction of cytoarchitectonic features including instance masks, semantic masks, outlines, coordinates, perimeters, and area for subsequent analyses. For immunofluorescence sections, we independently segment nuclei or cells under different channels. The instance segmentation of cell nuclei in the 408 nm channel is shown in Fig. 4S-U.

**Figure 4.**
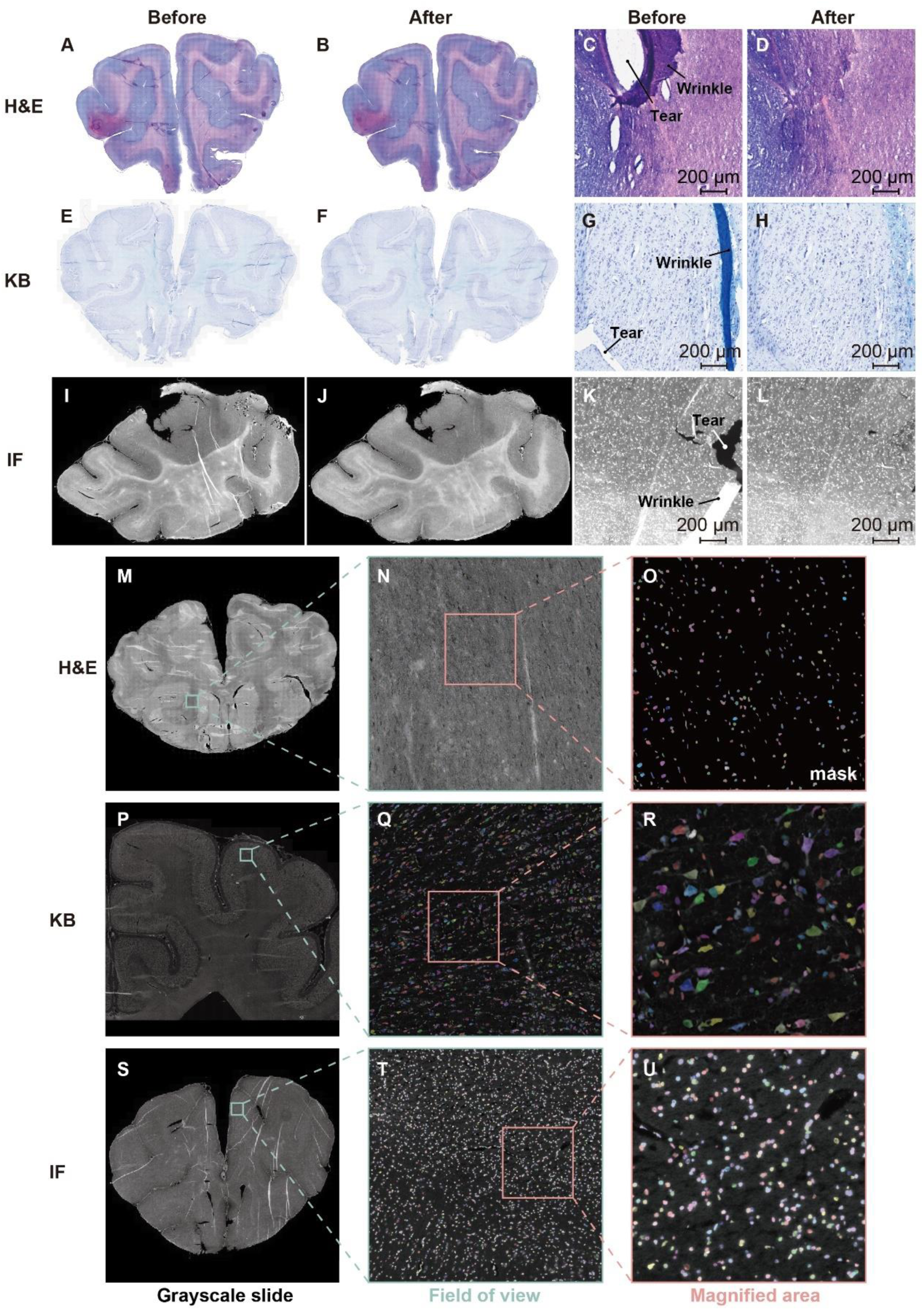
Cell instance segmentation and Inpainting for digital sections. (A-D) shows the images of a HE-stained coronal section before and after inpainting. (A) (B) (E) (F) (I) (J) are the panorama and (C) (D) (G) (H) (K) (L) are the expanded insets of H&E, KB staining, and immunofluorescence before and after inpainting repairment.

### Parcellation of Canine Visual Cortex via GLI

The brain parcellation is a vital target of brain atlas research, which helps to decode the brain heterogeneity with various zoning standards and connect the brain structures and functions. Cytoarchitecture is a fundamental principle for brain microstructural parcellation. The KB-stained serial coronal sections are employed for the cytoarchitectonic parcellation. The visual cortex plays a crucial role in processing visual information in both canines and humans, including facial recognition and emotional cues related to social behavior and relationships ^20^ ^21^. In addition, studies have shown the direct connection between the olfactory bulb and occipital lobe, functionally the visual cortex in canines, revealing a new link between canine smell and vision^22^. We pick the canine visual cortex of the left hemisphere to calculate the streamlines that traverse the laminar cortex and the space between streamlines is called a column or block (Fig. 5A and 5B). An ordinal number of the streamlines is counted incrementally from the suprasylvian sulcus towards the longitudinal fissure. The GLI profiles include mean GLI value, the center of gravity in the x-direction, standard deviation, skewness, kurtosis, and analogous parameters of the profiles’ first derivatives. The GLI profiles characterize the cytoarchitectonics facilitating the identification of zone boundaries with the largest cytoarchitectonic statistical differences, for example, the border 220 with the blocksize of 24 on both sides as region a and b in visual cortex (Fig. 5C-F). We compare the normalized GLI profile for a and b region, where the peaks and valleys of the two curved are not aligned with an overall lower GLI for region a. Maximal Mahalanobis Distance (MD) was detected as sliding window moved to profile position of 220, indicating the border between the region a and b (Fig. 5D). The locations of the maximum MD (Fig. 5E) and the MD frequency peaks (Fig. 5F) provide statistical evidence for boundary discovery. Eventually, the GLI identifies nine boundaries and ten subregions in the visual cortex (Fig. 5G). The average GLI profiles representing the laminar distribution of cytoarchitectonic (blocksize is 10) in these ten subregions are shown in Fig. S17. Three dysgranular regions (streamlines 170-220, 220-260, 260-334 respectively) discovered by GLI MD test can be visually inspected to evaluate the cytoarchitectonic difference in Fig. 5H. Comparing with region 170-220, layer IV is absent in region 220-260 and the transition between layer III and layer V is less pronounced. In addition, layer V of region 220-260 exhibits a higher density of larger pyramidal cells, and the boundary between layer VI and the white matter is much sharper. Compared with region 220-260, region 260-334 has a distinct demarcation between layer II and layer III and Layer III has higher cell density, increasing with depth. In addition, layer V of region 260-334 has smaller pyramidal cells, which are oriented to layer I with a gradual transition into layer VI (Fig. 5H). Region 334-478 and region 478-546 are agranular in Fig. 5I. In comparison to region 334-478, region 478-546 lacks layer IV, layer II has less granule cells and sparse pyramidal cells from layer III to layer VI (Fig. 5I). Comparing with region 478-546, region 546-630 is dysgranular and layer II and layer III have an increased cell density. Layer IV and larger pyramidal cells of layer V are easily recognized due to dark staining. Moreover, layer VI naturally transits towards the white matter without sharp boundary (Fig. 5I).

**Figure 5.**
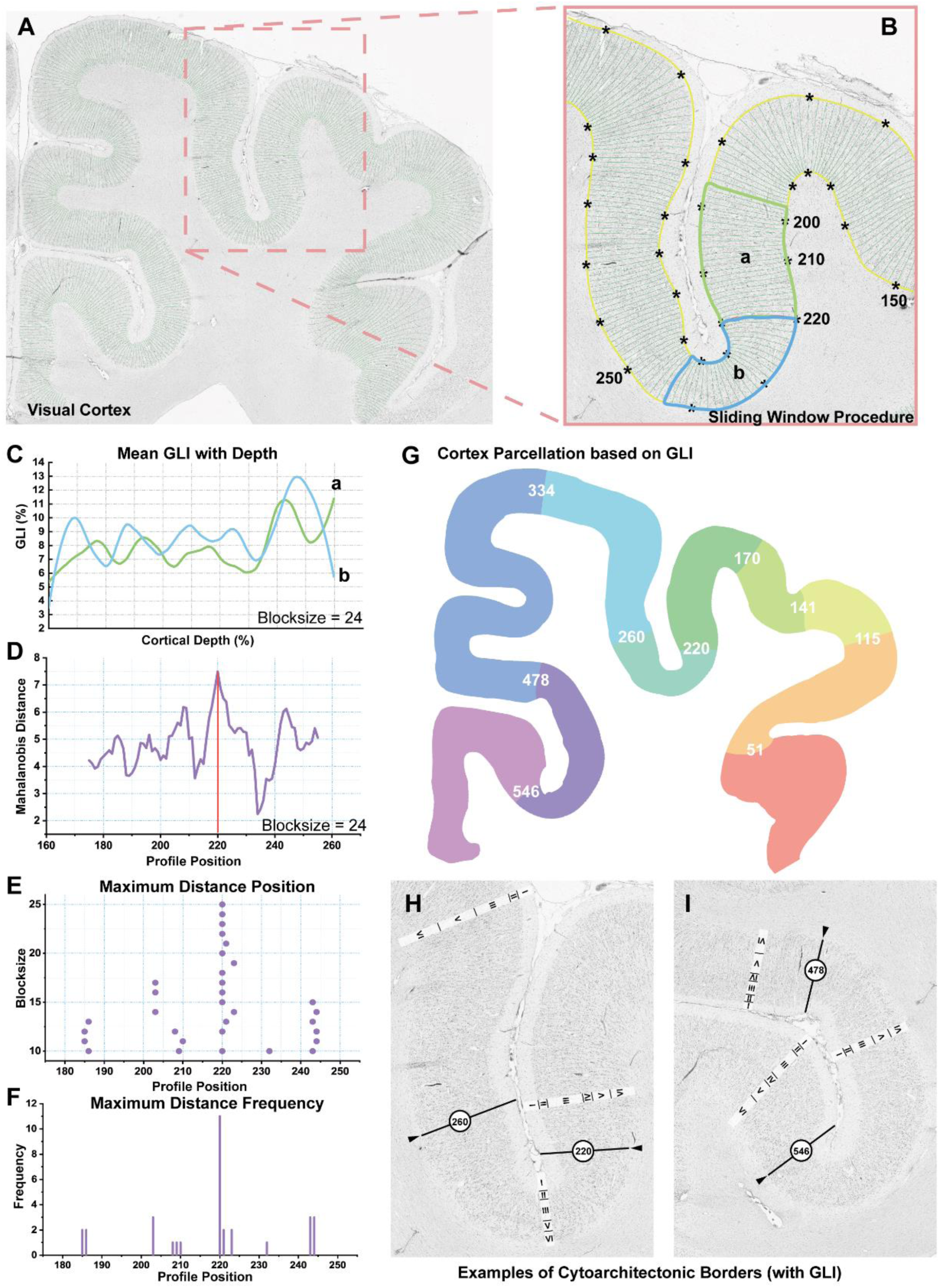
The cytoarchitectonic parcellation of the canine visual cortex with GLI. (A) The left hemispherical visual cortex of a coronal brain section with KB staining. The image is grayscaled and the green lines represent the streamlines starting to form the outer contour line (between layer I and II) to the inner contour line (between layer VI and the white matter). (B) The expanded inset of (A). The yellow boundaries represent the outer and the inner lines. We define the serial numbers of streamlines increase from the suprasylvian sulcus towards the longitudinal fissure. Every tenth-line interval is labeled with an asterisk. Within a sliding window with two equidistant blocksize, mean GLI profiles are computed for each side along streamlines, representing the cytoarchitectonic composition of the corresponding cortical section. The blue and green boxes are examples that represent two equally spaced blocks when the block size is 24 and the sliding window slides to No.220 streamline. (C) The mean GLI profiles of blocks **a** and **b** in (B), showing the variations in cytoarchitectonic composition with cortical depth. (D) Significant changes in the profile structure are measured by using the MD reaching maximum at profile position 220 with blocksize 24. (E) Across the region from profile position 175 to 255, the significant maxima position (purple dots) depends on the block size, where most dots are aligned at profile position 220. (F) Corresponding frequency of significant maxima at different profile positions across block sizes 10–25. The highest frequency occurred at profile position 220, as a putative cytoarchitectonic border. (G) The borders and subregions of the visual cortex are found by GLI. (H) and (I) show examples of cytoarchitectonic borders and the laminar cytoarchitecture of subregions parcellated by GLI. The solid black lines depict the position of the MD maximum and the boundary of visual cortex.

### Hypergraph Cytoarchitectonic Parcellation for Visual Cortex

Although GLI can parcellate cortex via the quantitative cytoarchitecture, it ignores intercolumnar connections and over-generalization of cytoarchitecture. We develop a Hypergraph Cytoarchitectonic Parcellation Method (HCPM) for the fine-grained zonation of cerebral cortex (Fig. 6A). Hypergraph models allow for high fidelity representation of data that may contain multi-way relationships^23^. Hypergraph convolution enables efficient information propagation between vertices by fully exploiting high-order relationships and local structures within the data ^24^. The hypergraph aggregates features from neighboring vertex through hyperedge convolution. The vertices are the cortical columnar units demarcated by the cortical inner and outer boundaries and streamlines from GLI. We calculate the cell morphological features (listed in Table 1) by utilizing masks of cell instance segmentation to determine the boundary of individual cells. The cell features from each columnar unit on KB-staining images are aggregated to form vertex feature vector. The cytoarchitectonic features of the vertex consist of 10 GLI profile features, cell density, and 38 cell morphological features. We define hyperedge to contain *k* neighboring columnar units due to the connectivity between cortical columns. For example, the hyperedge I contains the 1^st^ to *k*^th^ columns, and the hyperedge II will comprise the 2^nd^ to *(k+1)* ^th^ columns. We define this process of creating hyperedges on cortical columnar regions as the *k*-size sliding hyperedge. We generate 21 hypergraphs with different sliding window size for hyperedge construction as *k* varies from 10 to 30. Hypergraph convolution is then utilized as the encoder and fully connected network as the decode for deep embedding clustering (Fig. 6A). We obtain 21 distinct clustering outcomes because each hypergraph gives one clustering output. A streamline is regarded as a potential parcellation boundary if the neighboring five regions on either side belong to different clusters or zones after hypergraph learning. We summarize the frequency of each streamline as a potential boundary for all 21 clustering results (Fig. 6B). Streamlines with frequency peak above five are defined as weak boundaries. If at least three consecutive slices show weak boundaries at the similar positions, we define them as robust boundaries to delineate cortical subregions. The HCPM finds 16 borders and 17 subregions in canine visual cortex (Fig. 6C), discovering additional eight borders compared with GLI. Richer cytoarchitectonic features and intercolumnar connections lead to fine-grained parcellations. Borders at streamlines of 115, 167, 222, 252, 476, and 543 are identified boundaries by both GLI and HCPM (Fig. 6D). Consistent boundaries demonstrate the validity of the HCPM for cytoarchitectonic delineation. Boundaries based on different cytoarchitectures are different. We can see the variations in boundary location for 49 cytoarchitectonic features across the entire visual cortex with the heat map (Fig. S18). Fig. 6E shows the mean and standard deviation of two cytoarchitectures: the direction of the longer axis on Feret diameters and the cellular contour area. The visualization of pseudocolored cytoarchitecture demonstrates the continuity and heterogeneity of feature distribution in the cortex. Region 476-543, as a single region in GLI, has been subdivided into three subregions: region 476-492, region 492-516, and region 516-543 in HCPM (Fig. 6F). Region 476-492 has greater density of large pyramidal cells in layer V in contrast to region 492-516. Additionally, region 476-492 displays a sharper boundary between layer V and layer VI. Compared to region 492-516, region 516-543 has a more pronounced thickness in layer I, while layer II demonstrates a reduced occupancy of granule cells. Meanwhile, layer III and layer V in region 516-543 appear thinner, and cells in layer VI orientate parallel to cortical surface.

**Figure 6.**
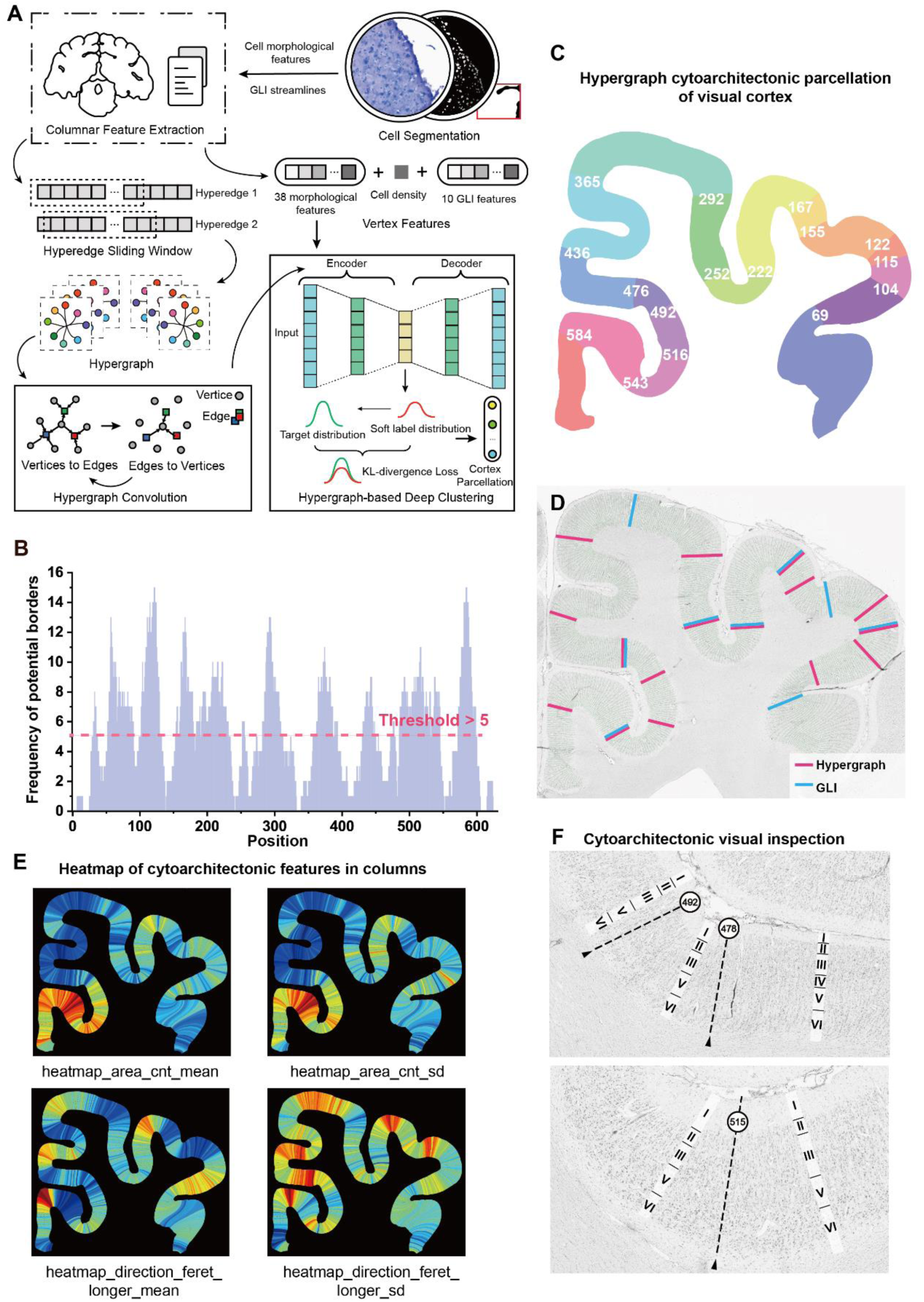
Hypergraph cytoarchitectonic parcellation and cytoarchitecture of canine visual cortex. (A) Illustration of models and pipeline of the hypergraph cytoarchitectonic parcellation method (HCPM) for cortex. (B) The frequency of different streamlined positions becomes potential boundaries across all clustering results. Positions with counts greater than five are considered weak boundaries. (C) The borders and subregions of the visual cortex parcellated by the HCPM. (E) Comparison of the borders produced by the GLI and HCPM in the same region. (F) The heatmap of four cytoarchitectonic features in the visual cortex: the mean of contour area, the standard deviation of contour area, the mean of longer ferret diameter, and the standard deviation of longer ferret diameter. (G) The HCPM parcellates one single region 476-543 of GLI into three subregions: region 476-492, region 492-516, and region 516-543. This panel shows the cytoarchitectonic borders of the HCPM and the laminar cytoarchitecture of subregions.

**Table 1.**
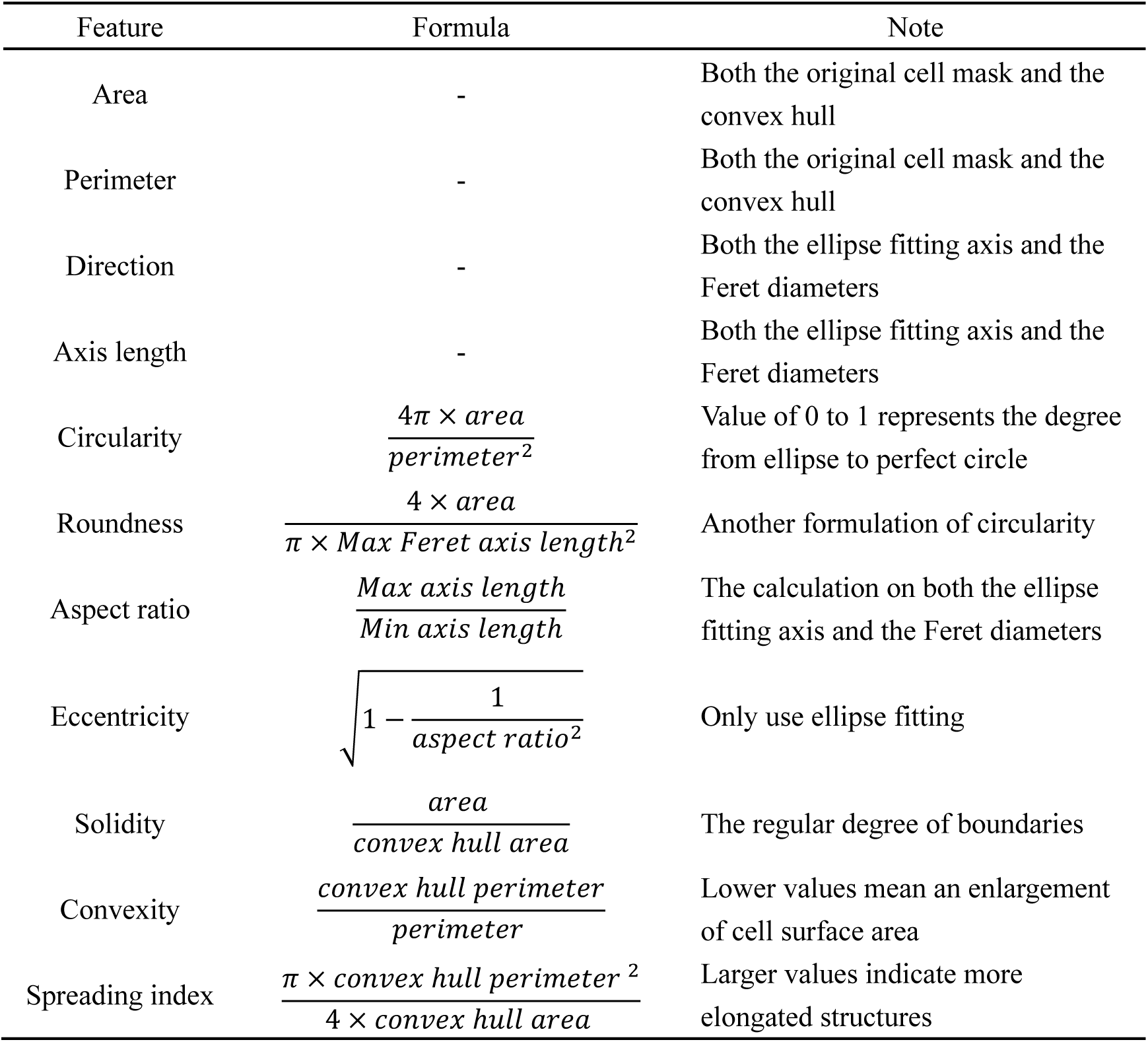
Formula for calculating cell morphological features.

| Feature | Formula | Note |
| --- | --- | --- |
| Area | - | Both the original cell mask and the convex hull |
| Perimeter | - | Both the original cell mask and the convex hull |
| Direction | - | Both the ellipse fitting axis and the Feret diameters |
| Axis length | - | Both the ellipse fitting axis and the Feret diameters |
| Circularity | $\frac{4\pi \times area}{perimeter^2}$ | Value of 0 to 1 represents the degree from ellipse to perfect circle |
| Roundness | $\frac{4 \times area}{\pi \times Max\ Feret\ axis\ length^2}$ | Another formulation of circularity |
| Aspect ratio | $\frac{Max\ axis\ length}{Min\ axis\ length}$ | The calculation on both the ellipse fitting axis and the Feret diameters |
| Eccentricity | $\sqrt{1 - \frac{1}{aspect\ ratio^2}}$ | Only use ellipse fitting |
| Solidity | $\frac{area}{convex\ hull\ area}$ | The regular degree of boundaries |
| Convexity | $\frac{convex\ hull\ perimeter}{perimeter}$ | Lower values mean an enlargement of cell surface area |
| Spreading index | $\frac{\pi \times convex\ hull\ perimeter^2}{4 \times convex\ hull\ area}$ | Larger values indicate more elongated structures |

### The GLI and HCPM for Human and Canine Motor Cortex Parcellation

We apply the GLI and HCPM to parcellate the premotor cortex of a Nissl-stai ned coronal slices of the right hemisphere from a 34-year-old human brain downloaded from a public database, the *Atlas of The Developing Human Brain* (https://www.brainspan.org/static/atlas). We select three consecutive Nissl-stained coronal slices and manually segment the premotor cortex (area 6) (female, 34 y ears old, https://www.brainspan.org/ish/experiment/show?id=100149965, tissue ind ex = 1321, 1325, 1329) (Fig. 7A). The GLI identifies nine borders and ten subregions while the HCPM finds 16 borders and 17 subregions (Fig. 7B, C). There is a strong consensus between these two methods regarding the localization of border 761 and 860 as well as a weak consensus for boundaries of 418 and 1022. Interestingly, these two boundaries align closely with the boundaries identified in the Modified Brodmann Atlas, which divides area 6 into the laterodorsal, the lateroventral, and the medial subdivision via visual inspection of cytoarchitecture (Fig. 7D). To study the relationship between the cytoarchitectonic and the functional regions of cortex, we manually align the functional mode 1024 atlas of the *Human Brain Project* (https://www.humanbrainproject.eu/en/science-development/focus-areas/brain-atlases/) to our specific slice. In area 6, the functional atlas consists of 13 borders and 14 subregions, ten of which match the boundaries of the HCPM, indicating the feasibility of the HCPM for associ ating cytoarchitectures and cortical functions (Fig. 7E).

**Figure 7.**
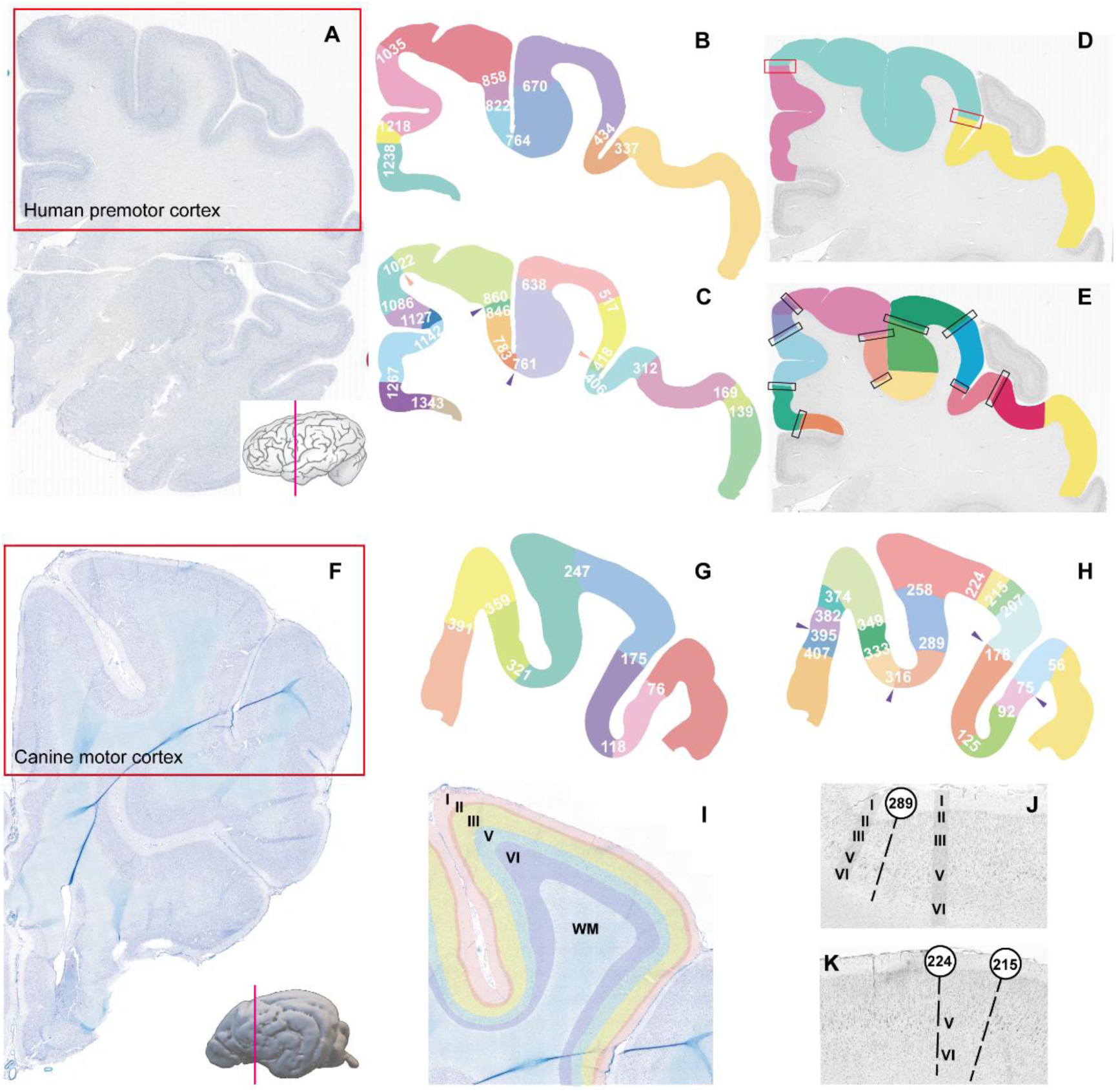
Cytoarchitectonic parcellation of the human and canine motor cortex. (A) A Nissl-stained coronal section of the right hemisphere of the human brain. The magenta line on the brain model marks the spatial location of this slice. The premotor cortex (area 6) is marked as ROI with the red rectangle. (B) and (C) are the parcellations of area 6 based on the GLI and HCPM, respectively, with distinctive colors marking different subregions and streamline numbers labeling borders. Both methods identify the same boundaries marked with purple arrows; whereas weak boundaries are marked with orange arrows. (D) Modified Brodmann cortical map of area 6 in this section. The rectangles mark the borders pointed out by the GLI and HCPM. (E) The area 6 parcellation with the human brain functional modes 1024 atlas. Black rectangles point to the borders that match the HCPM results. (F) A KB-stained coronal section of the right hemisphere of the canine brain. The magenta line marks the spatial location of the slice, and the red rectangle labels the canine motor cortex (M1). (G) and (H) are the M1 parcellations with the GLI and HCPM, and the purple arrows point to borders that are recognized by both methods. (I) An example of laminar delineation of the M1 with five layers. WM, white matter. (J) and (K) show the cytoarchitecture of the HCPM parcellating subregions.

We select and manually delineate the canine KB-stained serial sections of motor cortex (Fig. 7F). The GLI divides this cortex region into seven borders and eight subregions, with four boundaries recognized by the HCPM, which identifies 17 boundaries and 18 subregions (Fig. 7G, H). The laminar pattern of the canine motor cortex comprises five layers via visual inspection of cytoarchitecture (Fig. 7I). Layer III gradually transits into layer V, lacking the internal granular layer that traditionally corresponds to layer IV. Layer V have the largest and most intensely stained pyramidal neurons, which displays a similar cytoarchitecture of the motor cortex with other mammals (Fig. 7I). We pick border 289 and validate parcellations of the motor cortex by the HCPM through visual inspection of cytoarchitecture. Compared with region 258–289, layer II of region 289–316 is wider, while layer III shrinks; layer V has denser pyramidal neurons orthogonal to cortical surface; layer VI rapidly narrow down showing a sharper boundary with white matter. In Fig. 7K, regions 215-224 manifest cytoarchitectonic disparities in layer V comparing to adjacent regions. Specifically, layer V of region 215-224 shows a decreased abundance of large and deeply colored pyramidal neurons. Additionally, both GLI and the HCPM also provide consensus boundaries with the myeloarchitectonic parcellations in the canine visual and motor cortical slices according to the MRI atlas. Consensus boundaries between cytoarchitecture and myeloarchitecture seem to be identified in the transition area between the sulcus and gyrus (Fig. S19) ^13^.

### The Morphology of Canine Choroid Plexus from Histological Images

The canine brain contains four ventricles and the choroid plexus (CP) distributes in all ventricles. We manually segment all ventricles and reconstruct their surface model using the canine MRI template. Each cerebral hemisphere has a lateral ventricle (LV). The third ventricle (TV) is in the diencephalon, while the fourth ventricle (FV) is situated between the cerebellum and the brainstem. The TV and FV are interconnected by the cerebral aqueduct (Fig. 8A and 8B). The CP is an epithelial-endothelial complex, comprising a highly vascularized stroma with connective tissue and special glial cells known as epithelium (Fig. 8C). The CP serves as a barrier separating the blood and the cerebrospinal fluid (CSF) within ventricles. In the central region of LV, the CP is a thin undulating veil and ridge-like prominences of the epithelium encapsulating the underlying capillaries. Between the capillaries and epithelium, there is a loose connective stroma consisting of fiber bundles and cells. Capillaries are arranged in a sheet-like manner parallel to both the longitudinal axis of the plexus and each other (Fig 8D-8H). Fig. 8E demonstrates the continuity between the CP located in the LV and the TV. The CP within the TV is thicker and has a richer continuous stroma compared to that of the LV. The vascular orientations in CPs are parallel, where CPs are connected with epithelium bundles with varying thickness and cell organization pattern. Compared to the central part of CP in LV, the CP in temporal horn has a morphology of polygonal shape and occupies a greater ventricular space (Fig. 8G, 8H). Additionally, the CP of LV attaches to the hippocampus and the optic tract, within which the CP epithelium transitions into the ependyma at the tip of these brain parenchyma. The CP is thin and narrow in the atrium of the LV, showing a stretchy morphology at both ends and a compact middle segment (Fig. 8I). The CP structure changes at its lateral boundary, where the capillaries show various degrees of tortuosity with extended loops toward the ventricle. The glomus, at the confluence of the narrow central tunnel and temporal horn of the ventricle, displays a higher degree of tortuosity. The CP in the FV is highly lobulated and displays greater structural complexity compared to its counterpart in the LV (Fig. 8J-L). The loops between CP and capillary have smooth inner capillary and rough CP cross section with various protrusion toward the ventricular lumen, depending on the section plane. The capillaries are more intimately associated with the overlying epithelium in CP of FV due to more enveloped epithelium and thinner interstitial connective stroma (Fig. 8J-L). The CP in the TV shows morphological features that exist in both the CP of the LV and the FV, consistent with previous reports (Fig. 8M-O). Although the CP of TV has a smaller physical size, it exhibits the highest stoma thickness and ventricle occupancy.

**Figure 8.**
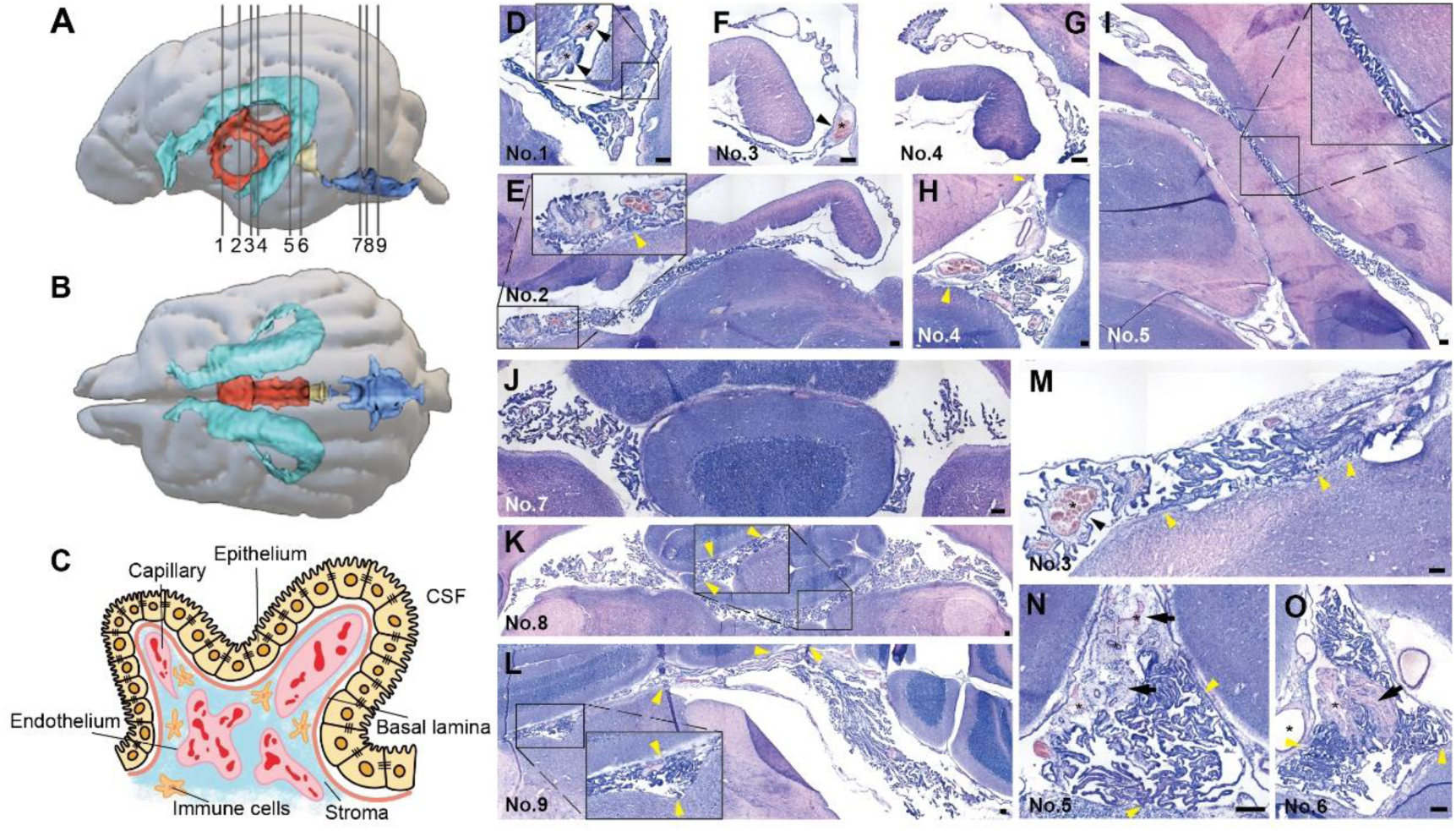
The morphology of the choroid plexus (CP) on the H&E stained sections. (A) and (B) are surface models of the canine brain (gray) and the ventricles from lateral (A) and dorsal (B) directions. The lines with serial numbers in (A) represent the positions of selected coronal sections. The lateral ventricles in lake blue, the third ventricle in orange, the cerebral aqueduct in yellow, and the fourth ventricle in blue. (C) is a schematic illustration of the CP anatomy. (D)-(I) are transverse sections of the CP in the right lateral ventricle. (E) is the CP from the third ventricle, (D)-(G) show the CP in the central narrow channel and (H) indicates the CP in the temporal horn of the lateral ventricle. (J)-(L) show the CP from the fourth ventricle, and the CP from the third ventricle is shown in (M)-(O). The numbers in the lower left corner of all slices correspond to the slice numbers in (A). Epitheliums (black arrowheads) are separated from underlying capillaries (*) by relatively large amounts of connective stroma (black arrows), and the attachment between the CP and brain parenchyma is pointed out by yellow arrowheads. (Scale bar 200 μm)

### The Morphology of Canine Leptomeninges

The meninges play a vital role in protecting and supporting the central nervous system. To our best knowledge, systematic comparisons of microscopic morphology and histology in different areas of the canine meninges are scarce ^25^ ^26^. We can visualize the morphology of the fixed leptomeninges from canine serial coronal brain sections with H&E staining. In vivo, meninges consist of three layers: the dura mater, arachnoid, and pia mater (Fig. 9A). The arachnoid and pia mater, together known as the leptomeninges, are separated by the CSF-filled subarachnoid space and interconnected by arachnoid trabeculae. The pia mater imprints the contour of the brain and spinal cord, which is highly vascularized and the innermost layer. The surface of pia mater contains the network of blood vessels, encasing blood vessels deep into the brain with perivascular spaces, which show pseudolymphatic function (Fig. 9A). To study the heterogeneity of meninges covering diverse brain regions such as the cerebral gyrus, sulcus, subcortical region, basal ganglia, ventricles, and cerebellum, we select Regions of Interest (ROIs) from target slices along the rostral-caudal axis (Fig. 9B). The dura mater is removed during dissection in our isolated brain and the leptomeninges is the remaining meninges.

**Figure 9.**
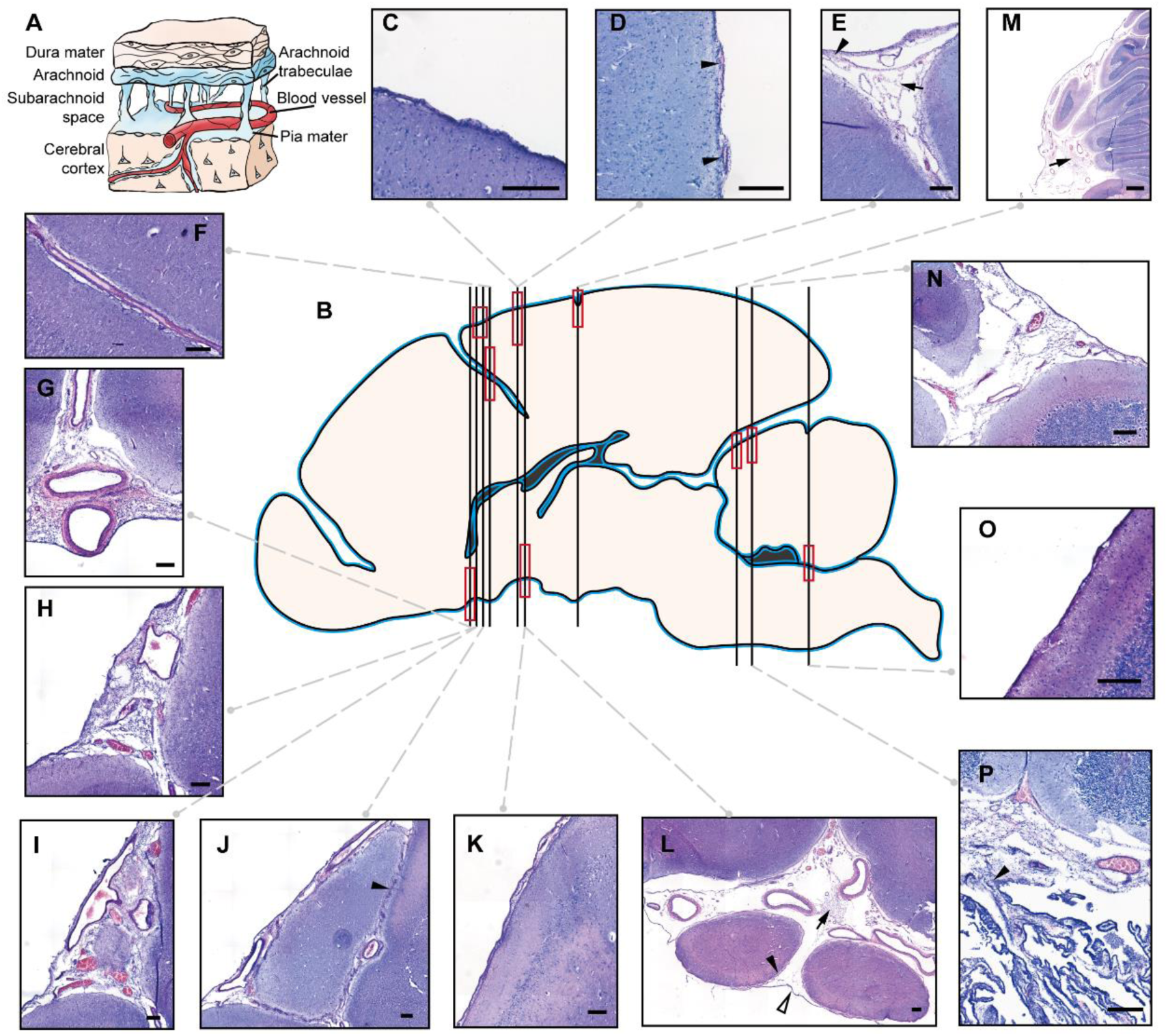
The morphology of the canine leptomeninges on the H&E staining slices. (A) Illustration of the anatomy of the meninges. (B) A schematic shows the canine brain sagittal section. The blue line represents the distribution of the leptomeninges. The black lines show the positions of the selected coronal slices, and the red boxes are the spatial locations of the leptomeningeal ROIs. (C)–(P) ROIs encompass various morphological characteristics of meninges from the H&E stained sections. (C) and (D) are pia mater covering the cerebral gyri, demonstrating different morphology of encapsulating versus exposed blood vessels, respectively. Blood vessels are marked with black arrowheads. (E) The morphology of the leptomeninges in the transition region between the sulcus and gyrus. (F) The morphology of the pia mater and vessels deep into the sulcus. (G) The morphology of the leptomeninges covering the medial olfactory tracts of two hemispheres. (H)-(J) are ROIs showing the morphological variations of the leptomeninges in nearby slices. The black arrowhead points out the position where the pia maters of both sides are so close together that it is difficult to distinguish the layers. (K) shows the pia mater on the pyriform cortex, and (L) shows the leptomeninges covering the optic tracts and the olfactory stria. The arachnoid trabeculae, black arrow; the pia mater, black arrowhead; the arachnoid, hollow arrowhead. (M)-(N) show morphology of the leptomeninges covering the cerebellar vermis. (O) shows the pia mater from the cerebellar gyrus and (P) indicates the continuity of the pia mater and the choroid plexus in the fourth ventricle (black arrowhead). All scale bars are 200 μm except the 1 mm scale bar of (M).

**Figure 10.**
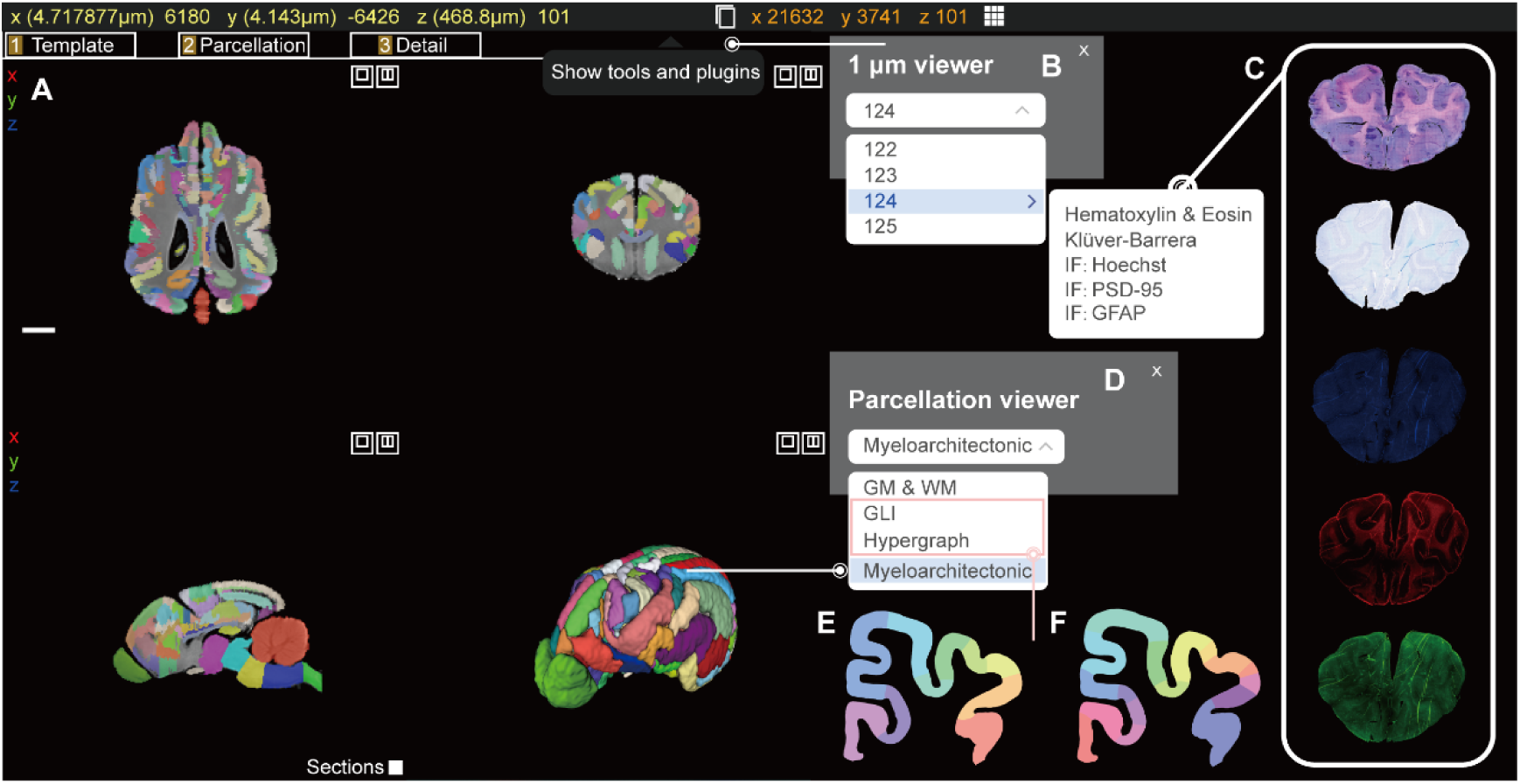
The interactive interface and functionality demonstration of our canine brain atlas. (A) 3D reconstructed surface models and three views (horizontal, coronal, and sagittal) of the MRI template and myeloarchitectonic parcellation. (B) Users can choose the "1-micron display tool" option from the interactive interface’s "Tools and Plug-ins" icon, which has a drop-down menu to select the corresponding number of the MRI coronal plane and the staining image. (C) Images of three staining techniques are presented: H&E, KB, and IF. The IF has three-channels including Hoechst, the PSD-95, and the GFAP with the resolution of 1 μm/pixel. Selecting the "Parcellation" can show the outcomes of various partitioning techniques, including the gray and white matter segmentation, the GLI cytoarchitectonic cortical parcellation, the HCPM parcellation, and the myeloarchitectonic maps. (E) and (F) are cortical parcellation examples of the GLI and HCPM, respectively, which are part of the visual cortex.

The leptomeninges that cover the cerebral gyrus is easily recognized as the pia mater due to the appearance of a single-layer thin structure on the slice (Fig. 9C). The pia mater wraps around the blood vessels, and its morphology undulates with them (Fig. 9D). The meninges at the transition between the gyrus and sulcus has network structure and thicker than that of the gyrus, which expand and stratify into layers of the arachnoid and pia mater. The arachnoid trabeculae facilitate the formation of subarachnoid space (Fig. 9E). In the sulcus, the penetrated blood vessels are separated from the brain parenchyma by the pia mater. The pia mater and the blood vessels display intermittent cavity but robust association, characterized by a noticeable gap (Fig. 9F). The leptomeninges is thicker in the ventral brain, specifically in basal ganglia and the intersection of two hemispheres. The arachnoid serves as the external boundary due to the absence of dura mater. Arachnoid trabeculae surround blood vessels while connecting the arachnoid and the pia mater (Fig. 9G). The morphology and thickness of the meninges depends on the angle of the sectioning, demonstrating variations of the meninges from the gyrus surface with denser and clearer arachnoid trabeculae (Fig. 9H-J). Fig. 9J shows the connecting meninges between adjacent cerebral gyri, where they encase the blood vessels penetrating deeply into sulcus with a vague border between two meninges layers (Fig. 9J). The leptomeninges from the ventral pyriform cortex is thicker than that from the dorsal gyrus (Fig. 9H). The pia maters wrapping two optic tracts and the olfactory stria are independent covered entirety by an arachnoid layer, and the space among them is filled with arachnoid trabeculae and ventral vessels (Fig. 9L). The folia and fissures of cerebellar vermis are denser than the cerebral gyri and sulcus, where folium is wrapped by pia mater and the vermis is enveloped by the arachnoid (Fig. 9M). The subarachnoid space between the cerebral vermis and the pons makes the leptomeninges thicker than that of other vermis areas due to the abundant vessels and arachnoid trabeculae. Arachnoid trabeculae in the transitional zone from vermis to pons are encapsulated by an extensive network of blood vessels, occupying a large area, where vascularization increases the leptomeninges thickness in this region (Fig. 9M, N). The leptomeninges from the cerebral and the cerebellar gyri have morphological similarities, and the latter one seems to detach more easily from the cerebellar dura mater leading to continuous boundary contours (Fig. 9M-O). Fig. 9P demonstrates the continuity from the CP to the pia mater within the FV.

### Visualization of the Canine Brain Atlas via the Interactive Web Server

The advent of public digital platforms represents a pivotal trend in modern brain atlases, which transcends the limitations imposed by traditional physical maps or eBooks in terms of atlas resolution and accessibility, enabling the interactive visualization of brain images from macro to cellular scale. Moreover, web server serves as a universal framework to integrate multimodal brain images and data across species, amplifying the functionality of brain atlases and value in research and clinical applications. To our knowledge, the interactive, multimodal, cross-scale canine brain atlas is scarce. Thus, we develop a multimodal canine brain atlas accessed through online database with a user-friendly interactive interface. The average MRI template with 0.5 mm voxel resolution serves as a reference for spatial alignment and 3D constructed surface model. Users can zoom, rotate, and shift across three views to access or visualize desired region. Stained sections aligned to various MRI coronal planes are available for visualization including three staining protocols employed: H&E, KB, and IF, where IF has the Hoechst, PSD-95, and GFAP channels.

Interactive plug-ins with drop-down menus facilitate the filtering of images by allowing users to select specific sections and staining protocol. The interactive visualization is available for histological images, and we initially upload compressed histological images at a resolution of 1 μm/pixel to resolve internet loading challenges. We provide a link for downloading the original 0.5 μm/pixel resolution image. Various partitioning strategies can be interactively displayed based on user preference. Our interactive atlas supports the visualization of 3D surface models and three-view representations depicting the structural parcellation of myeloarchitectonic and the segmentation of gray and white matter. The GLI and HCPM partitions are temporarily restricted to the cortices where validation has been conducted. Efforts are undergoing to encompass parcellations of the entire canine cortex.

## Discussion

Interspecies brain atlas has enriched our knowledge scope, contributing to a deeper understanding of brain function, evolution, development, and disease. Despite the evolutionary divergence between dogs and humans, canines demonstrate high congruence with humans in brain structure, gene expression and proteomics ^27^. Furthermore, as a species domesticated and evolved alongside with humans, dogs exhibit adeptness in unique cognitive behaviors within human societies than non-human primates ^28^ ^29^. Canine’s physical size, similarity to human neurological diseases, and experimental conditions render them greater clinical translational value. Therefore, Canine emerges as a potential model for comprehending human brain structures and functionalities, especially in studies of cognition and aging. However, the existing canine brain atlases with a coarse-grained resolution present a bottleneck for in-depth and cross-species research. Public resources for histological canine brain atlases exist in physical or eBook formats ^30^. Available canine atlases rely on a singular imaging modality and exhibit limitations in spatial resolution. The coarse segmentation employed hampers the precision of brain structure-function mapping and impedes their potential as a universal framework for effectively integrating multimodal information.

To overcome these limitations, we have developed a multimodal and cross-scale canine brain atlas. At the macroscopic level, we describe the brain morphology and cortical folding patterns of the canine brain. We generate an MRI template and manually delineate and reconstruct cortical myeloarchitectonic partitions with a spatial resolution of 500 μm. On the microscopic scale, we collect high-resolution images of canine brain sections with multiple staining, including H&E, KB, and IF to visualize cell nuclei, neurons, nerve fibers, astrocytes, and PSD-95, achieving a resolution of 0.5 μm. The integration of multimodal and multiscale brain images significantly improves the localization and delineation of brain structures. This fine-grained brain atlas is crucial for studies or surgeries involving deep brain stimulation for canine. Our canine brain atlas is accessible on an interactive online platform, facilitating the integration and display of multimodal and multiscale brain atlas as well as partition generated from various methodologies. Similar to the Allen Brain Atlas and the Human Brain Project ^4,31^, our interactive atlas not only improves the readability and accessibility of the canine brain but also serves as a framework for cross-species brain information.

Our high-resolution microscopic canine brain images provide a foundation for detailed cortical segmentation, particularly in cytoarchitecture. Cytoarchitecture is determined by gene expression, leading to distinct patterns across brain regions ^32^ ^33^. During neurodevelopment, various cortical structures form through processes such as neuronal migration, differentiation, and synapse formation. These processes establish specific structural patterns, defining the cytoarchitecture and functional characteristics of the adult brain. Cytoarchitecture, reflecting the heterogeneity of cellular physical arrangement, offers a unique perspective for understanding neural circuit functions and information processing within different brain regions. The spatial arrangement of cortical layer and subcortical nuclei demonstrates high heterogeneity, which is important for explaining brain function and individual variability. Cytoarchitecture is widely used in brain structural divisions as the basis for integrating and compressing multimodal biological information. The segmentation of the brain into spatial units is essential for decoding its inherent heterogeneity. Additionally, the widespread similarity in cytoarchitectural laminar patterns across mammalian cortices makes cytoarchitecture-based structural partitioning possible for cross-species applicability ^16^.

For the canine cortex parcellation, we use the observer-independent GLI and self-developed HCPM to do cytoarchitectonic segmentation and verify the partitioning results by visual inspection. The GLI is primarily employed and validated in the cytoarchitectonic parcellation of the human cortex ^2,18,34^. We attempt cross-species application of GLI in canine cortical zonation, achieving fine-grained segmentation. However, GLI simplifies cytoarchitecture into ten features, inevitably neglecting many cortical microstructures. Considering the continuity of neuronal arrangement within the cortex, the structural relationships between "columnar units" should be included. Our approach focuses on cytoarchitecture feature aggregation at the single-neuron level, compressing massive structural information into "columnar units" for cortical zonation, and each unit is represented by 49 cytoarchitectonic features. The HCPM integrates microscopically cytoarchitectonic features and relationships among cortical columns, yielding valuable cortical decoding and leading to finer-grained structural zonation in both canine and human brain. The variation of cytoarchitectonic features across cortical bands offers biological interpretability of boundaries through multiple structural features. Visual inspection identifies transitions in neuronal arrangement patterns at the newly generated boundaries, verifying the HCPM parcellation results. The large overlap between human brain functional MRI partitions and the HCPM parcellation indicates the potential of the HCPM in mapping the relationship between brain microstructure and function.

However, further validation of the biological mechanisms behind HPCM cortical divisions is needed in the future. Researchers have shown differential gene expression among GLI cytoarchitectonic subregions of the human brain cortex, the correlation between gene expression or electrophysiological signals with finer-grained zones remains to be explored. To study the correlation between functional and structural differences across subregions is important to ascribe known function to discrete zone and find cognitive functions of novel zones or network. Relating the fine-grained structural partitioning of the cortex to response or behaviors under different tasks will provide essential clinical and research value. We build the complete pipeline to segment cells, calculate cellular features, aggregate cellular features into columnar unit feature vector, and parcellate columnar units into zones, which is verified in multiple cortex regions. We will complete and release the fine-grained division and reconstruction of the entire canine cortex, resulting in a 3D cortical atlas of the canine brain. Considering inter-subject variability, we will incorporate cytoarchitectonic information from more samples and create a probabilistic atlas in the future. As our canine atlas and the HCPM rely on uniaxial brain imaging, to include other axial imaging to complement and balance errors introduced by the uniaxial approach is necessary. Caution is advised when directly applying brain atlas to different cranial species, especially brachycephalic dogs. Brachycephalic dogs exhibit cranial shortening, leading to a ventral tilt of the brain’s long axis and a ventral displacement of the olfactory lobe.

In summary, we have developed a new canine brain atlas and a novel approach for fine-grained cytoarchitectonic cortical parcellation based on hypergraphs. The hypergraph learning integrates richer cytoarchitectural features, yielding detailed cortical divisions. HCPM not only demonstrates cross-species versatility but also unveils the potential for fine-grained structural and functional mapping. The canine brain atlas, presented in an interactive platform, integrates multimodal macroscopic and microscopic brain imaging and provides canine brain partition maps using GLI and HPCM. Our atlas with high-resolution structural imaging and HCPM zonation surpasses the existing coarse-grained canine brain atlases, facilitating the precise localization and identification of structural features. The fine-grained brain atlas will stimulate research on canine brain development and aging as well as disease detection and therapy. This study advances our understanding of brain structure and function, providing insights into human brain evolution, development, function, and disease.

## Method

### Animals and Anatomy

We have ensured full compliance with all pertinent ethical regulations governing animal study. All animal experiments are approved by the Animal Care and Use Committee of Kunming Police Dog Base, China. We select four healthy male Kunming dogs at ∼3-5 years old (Table S4). All dogs undergo euthanasia before being dissected. To guarantee the procedure with minimal damage to the brain, we use an air drill and a pendulum saw to circumcise the dog skulls (Fig. S20). Meanwhile, it is necessary to open the nasal sieve plate to create enough space for the dissection of the olfactory bulbs. Isolated brains are promptly immersed in a solution of 10% neutral buffered formalin for overnight fixation, followed by preservation in the newly prepared fixative until further processing.

### MRI Acquisition

MRI of the four adequately fixed brains is acquired on a Siemens Tim Trio 3T scanner (Erlangen, Germany), using a human knee coil for signal acquisition. High-resolution T1-weighted anatomical images are recorded with parameters of TR = 2300 ms, TE = 3 ms, inversion time = 1000 ms, flip angle = 9^◦^, and acquisition voxel size = 0.5 * 0.5 * 0.5 mm^3^.

### MRI Template Creation and Parcellation

Fig. S21 depicts the pipeline of the MRI data processing. Firstly, the images of the four subjects undergo reorientation to establish the x-axis as the right-left orientation, the y-axis as the caudal-rostral orientation, and the z-axis as the ventral-dorsal orientation. Secondly, the origin of the images is manually adjusted to rostral commissure using SMP12. Thirdly, we apply N4 bias field correction to each image to reduce intensity non-uniformity. This correction is performed using SimpleITK, a Python library specifically designed for this purpose. Fourthly, considering the individual differences, the MRI volumes are registered to one population canine brain template before template construction, exploiting *Symmetric Normalization* in ANTs. Finally, we use the *MultivariateTemplateConstruction* command in ANTs to generate the average MRI template of canine brains.

The Automated Segmentation Tool of the Oxford Centre for Functional Magnetic Resonance Imaging of the Brain (FAST) is a widely employed method for brain tissue segmentation. Typically, FAST segments and classifies a three-dimensional brain into three categories: GM, WM, and CSF. Since our data is derived from ex vivo MRI scans, we classify the brain templates into GM, WM, and background. To ensure coherence with the MRI template, we manually adjust the tissue segmentation maps. The 3D surface reconstruction and smoothing of GM and WM templates are generated by using AFNI’s IsoSurface command. Moreover, according to the cortical myeloarchitectonic atlas and the brain neuroanatomy, we manually delineate the MRI template and reconstruct a 3D surface model by using the ITKSNAP.

### 3D Printed Chunking Mold of Canine Brain

We selected serial Computed Tomography (CT) images of the Kunming dog brain that are provided by the Kunming Police Dog Base. The consecutive CT slices have 1.0 mm thickness and the matrix size of 512*512. We manually segment the brain region from the dog’s skull and binarize each image. We use *Materialise’s Interactive Medical Image Control System* and the *Blender* to reconstruct the CT images into a three-dimensional model and smooth out sharp points. With Boolean operations in the software, we construct a brain chunking model capable of accommodating the entire dog brain. This model incorporates 22 knife grooves, each measuring 1.5 mm in width and spaced at 5 mm intervals. The grooves are equally distributed along the axis (from the rostral to the caudal), supporting the tissue chunking in the coronal plane. The mold is printed with polylactic acid through the HORI Z600 printer, which utilizes the fused deposition molding technique (Fig. S22).

### Section, Staining and High-resolution Imaging

We extract the canine brains from formalin and rinse them under tap water for two hours to remove the residual fixative on the sample surface. To facilitate subsequent dehydration and enable easy cryosectioning, we use the canine brain chunking mold to divide the samples into eight pieces along the coronal plane. These chunked samples are dehydrated in sucrose solution with a concentration gradient of 10%, 20%, and 30%, followed by blotting off surface water post-dehydration. We submerge the samples in the Optimal Cutting Temperature (OCT) compound and incubate them overnight to eliminate trapped air bubbles. Subsequently, we embed the samples in liquid nitrogen and store them in the ultra-low temperature refrigerator until sectioning. The brain blocks are crysectioned into groups, with each group containing ten consecutive coronal sections, each with a thickness of 20 μm, spaced at intervals of 500 μm (Fig S23). We select 900 sections and stain them with the H&E, following standardized staining procedures. We select 300 sections for the KB staining (cresyl violet for labeling nuclei and neuronal cell bodies, as well as Luxol fast blue (LFB) for labeling myelin). Additionally, 80 sections are designated for the immunofluorescence staining. All sections are mounted onto glass slides, desiccated for 24 hours at 4 ℃, and washed to remove OCT compound before staining. In the KB staining, sections are initially stained in a 0.1% LFB solution overnight and then rinsed with 95% ethanol and distilled water. We then differentiate the sections with the 0.05% lithium carbonate solution and the 70% ethanol until the gray matter appear white under microscopy. Sections are subsequently stained with 1% cresyl violet solution for 10 minutes at 37 ℃, washed with distilled water, and subjected to differentiation using 70% and 95% alcohol. Subsequently, sections are dehydrated in 95% and 100% ethanol, cleared in xylene, and finally coverslipped with permount™ mounting medium.

For immunofluorescence staining, each section is incubated with a trypsin antigen retrieval kit to repair antigens at 37 ℃ for 30 minutes, following removal of the OCT compound. We then wash away the excess trypsin with 0.1 M phosphate-buffered saline (PBS) solution, and incubate the sections for 3 hours at room temperature with a blocking buffer (containing 0.1 M PBS, 0.5% Trition X-100, 10% normal goat serum (NGS)). Incubation with primary antibodies (anti-PSD95 antibody (abcam18258, rabbit), 1:300; anti-GFAP antibody (abcam4674, chicken), 1:300) in a blocking buffer takes place for 14 hours at 4 ℃ with gentle agitation. Then a series of 10-min rinses in 0.1 M PBS solution is performed, and the sections are incubated in the secondary antibody solution, which contains a 1:300 dilution of goat anti-rabbit IgG H&L (Alexa Fluor 488) (ab150077) and goat anti-chicken IgY H&L (Alexa Fluor 647) (ab150171) within a blocking buffer. This incubation step is carried out for 2 hours at room temperature in darkness. We then wash the excess secondary antibodies with 0.1 M PBS and incubate sections with 1:100 Hoechst solution in a blocking buffer for 15 minutes. We rinse the section again and coverslip them with the fluoromount aqueous mounting medium (sigma, F4680). All panoramic digital images of sections are acquired with the Tissue FAXS plus Q+ system, which is equipped with the ZEISS imager z2 microscope. The resolution of the original image is 0.5 μm/pixel in plane and the field of view is 2048 * 2048, and original data are *.tiff* images. The excitation wavelengths of fluorescent imaging are 405 nm, 488 nm, and 647 nm.

### Artifact Detection and Inpainting of Histological Images

The process of artifact detection and correction is shown in Fig. S24. To initiate, a reference section for histogram matching and registration is manually selected for each group based on the criterion of exhibiting minimal artifacts. Since the sections within the same group are arranged consecutively and have a uniform thickness of 20 μm, they are display a comparable visual appearance. All other sections within the same group undergo intensity normalization and section-to-section registration to align with the reference section. Considering the requisite precision for detection and the computational burden associated with analyzing images at the megapixel level, we perform fold/tear detection on the image at a 20% resolution. To ensure the validity of the detected masks, a section mask is initially generated using a uniform threshold. We employ two main categories of detection strategies: the intensity-based and the comparison-based strategies.

The intensity-based strategies vary depending on the specific staining methods. For immunofluorescence and KB staining, fold detection employs an adaptive thresholding method, while the protocol of fold detection in the H&E staining follows the methodology outlined in the reference ^35^. Tear detection is achieved by using a threshold that is equivalent to generating the section mask because the pixel value within the tear range matches the background value. In the comparison-based strategy, a mean section is computed for each group of sections, followed by assessment of each section by comparing it to the mean section within its respective group. Specifically, for a pixel at position (x, y) in a certain section, the mean and variance of a block (e.g. 25*25) surrounding the pixel in the mean section are counted. If the absolute value of the discrepancy between the mean value within the block and the intensity of the pixel (x, y) surpasses two standard deviations, the pixel is designated as a fold or tear. However, this approach does not differentiate between folds and tears in the detected region. Finally, the identified locations of folding and tearing are delineated employing a binarized mask, which is resized to match the dimensions of the original segment. Portion of the mask that encompasses an area smaller than 0.0005% of the overall mask area is disregarded to optimize correction time. To enhance the handling of the edge portion of the fold and tear, we apply an edge expansion technique using a 10*10 kernel on the mask. Additionally, we eliminate any small disconnected areas present in the final mask. To address the artifact correction, we employ a similar method to that of BigBrain. We remove the identified artifact from each section and replace it with the corresponding portion from the nearest neighbor section. If the neighboring portion is also flagged as an artifact, subsequent closest neighbors are referenced until all slices within the group have been selected. The process entails an initial intensive detection phase, followed by an artifact correction step and two iterations of comparison-based detection and correction.

### Nucleus and Cell Instance Segmentation

We argue that fine-grained segmentation will provide more informative capacity for downstream tasks due to the data derived from biomedical experiments. Therefore, we decide to implement nuclear and cellular instance segmentation and calculate meaningful features in slices. Against other prevalent imaging data, our data inherently possess a magnitude of ten thousand high-resolution that are demanding and time-consuming to label even one holistic slice, thereby having to resort to pre-trained models for persuasive segmentation performance. However, the majority of current algorithms could not accommodate the discrepancy of nuclear/cellular diameters between training data and ours, giving rise to catastrophic results. After extensive empirical trials, we proceed to segment nuclei and cells with a canonical method called Cellpose^19^, which is seamlessly adapted to our objective due to its generalist characteristic and sensitivity toward nuclear/cellular diameters. The details of the neural network of Cellpose are shown in Fig. S25. Despite its salient capacity, cautious and rigorous adaptions of the preprocessing are required to handle the high-resolution data. Specifically, we adopt a fourfold autonomous pipeline called Split-Segment-Extract-Stitch (SSES) to tackle the challenge. **Split**: The high-resolution data is arguably handled with the divide-and-rule policy. We split one holistic slice into homogeneous patches with the size of 2048*2048 to satisfy the appetite of Cellpose. Zero padding is implemented on the right and bottom sides for potential scale matching and is eliminated afterward. **Segment**: Preprocessed with grayscale and polar inversion, the patches are fed into the chosen algorithm to produce instance segmentation harvests. **Extract**: Given the logits of segmentation, a crafted feature extraction paradigm is designed to compute practical features toward instance masks, semantic masks, outlines, coordinates, perimeters, and area. **Stitch**: Once all computation is completed, the sub-results of the patches are logically stitched to constitute an assembly homologous to the original high-resolution data. Finally, this pipeline endows the project with autonomously self-adaptive nuclear/cellular instance segmentation and practical feature extraction on multimodal slices with immunofluorescence, H&E, and KB staining.

### Cortical Parcellation Based on Gray Level Index

The rectangular ROIs are defined in the histological section and digital images with an in-plane resolution of 0.5 μm. We employ the Cellpose method to segment the nucleus and neuron cell bodies from the KB staining slices and provide a binary mask in the original resolution. GLI images are computed as each pixel quantifying the volume fraction of cell bodies within a measuring field of 16*16 pixels in the original digitized ROI. The GLI images encode the cell density of ROI as 8-bit gray value and then the corresponding GLI image is rescaled to the same size of ROI. We define an outer contour line (between layer I and II) and an inner contour line (between layer VI and the white matter), and the curvilinear traverses between both counter lines are calculated using a physical model based on electric field lines. Extracting GLI value profiles along these traverses capturing changes that reflect the regional cytoarchitecture. The profiles could be operationalized with ten features: mean GLI value, the center of gravity in the x-direction, standard deviation, skewness, kurtosis, and analogous parameters of the profiles’ first derivatives. To compare profiles that have different lengths, each profile is re-sampled with linear interpolation to a standard length corresponding to a cortical thickness of 100% (0% = the outer counter line; 100% = the inner counter line). The ten-element feature vector combines the features of the GLI profiles, which are used to calculate the Mahalanobis distance (MD) between two adjacent regions of profiles. The MD serves as an indicator of the cytoarchitectonic difference between profiles. A specific number of profiles (block size from 10 to 24) are consolidated into a singular block. The sliding window procedure is used to compute the MD for blocks surrounding each profile position in all block sizes across the whole cortical ribbon. A significant maximum of MD for different block sizes (block size from 10 to 24) indicates a change in the shape of the profiles and serves as a criterion for delineating an area border. A border is accepted if a significant maximum (Hotelling’s t-test, Bonferroni corrected, p<0.01) is consistently detected at the same position across several block sizes and is also present in at least three consecutive histological sections. Finally, each border is controlled by visual inspection of the histological images.

### Hypergraph Stacked Autoencoder

Autoencoder is a neural network trained in an unsupervised manner, and it is highly prevalent for clustering tasks ^36^. Hypergraph offers several notable benefits, particularly in the realm of mining nonlinear high-order information ^37^. The hypergraph stacked autoencoder (HSAE) is an architecture to acquire efficient feature representations by considering high-order information, building upon the basis of autoencoders. A stacked autoencoder enhances its efficacy in feature extraction by increasing the depth of the hidden layer. However, with the increasing depth of the hypergraph neural network, there arises a phenomenon known as over-smoothing or over-squashing. To mitigate this, the HSAE has been configured with a depth of two.

In each layer of the HSAE, the encoder function is defined as *Z* = *f*(*X*) and the decoder function as *X*′ = *g*(*Z*). The decoder is a fully connected neural network, while the encoder is a hypergraph convolution neural network. The loss function is the reconstruction loss:

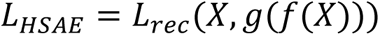

### Hypergraph-based Deep Embedding Clustering Method

The clustering model used in this study is based on deep embedding clustering, aimed at partitioning the columnar units into distinct clusters ^38^. Traditional clustering methods, such as k-means, perform clustering in the original feature space *X* and aim to optimize the clustering centers. In contrast, deep embedding clustering accomplishes this task in a smaller latent feature space *Z*, derived from *X*, to reduce the dimensionality of the feature and avoid the curse of dimensionality. It not only learns the clustering centers but also the nonlinear transformation *f*: *X* → *Z*, which is a HSAE in this work.

Soft labels *Q* represent the similarity between points and clustering centers by student’s t-distribution:

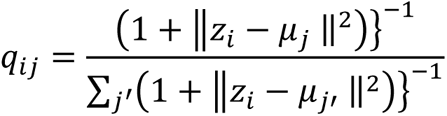

A target distribution P is got from Q by:

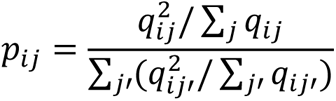

The clustering loss is the Kullback-Leibler divergence:

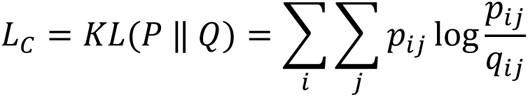

The HSAE is pre-trained before clustering. The initial clustering centers are set by the traditional k-means clustering method in latent feature space Z, which is obtained by a pre-trained HSAE.

Above all, the objective function of hypergraph-based deep embedding clustering is:

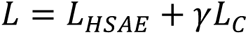

### The Static Website and Interactive Atlas

We employ JavaScript, HTML, and CSS to construct the static website, which contains basic information regarding this study as well as several data links. We offer an obvious entry point to access the canine brain atlas within this page. We select NGINX, a highly efficient and lightweight server well-regarded in the professional community for hosting the website. The presentation of the brain atlas facilitated through Neuroglancer, a powerful tool developed by Google, enabling an interactive 3D experience. The Neuroglancer-scripts project is employed to pre-process the brain atlas data for the Neuroglancer. Integration of the VUE front-end architecture seamlessly enhances interactive brain mapping capabilities. The procedural sequence is as follows:

1. Environment configuration: Initiate by setting up the Node.js environment and deploying both the Neuroglancer project and the Neuroglancer-scripts tool.
2. Data composition: The Neuroglancer-scripts project tool is used to convert the MRI in nii-format into pre-processed data, serving as the source data for the Neuroglancer interaction. The website showcases three datasets converted by Neuroglancer-scripts: the MRI template, coronal digital sections, and parcellation maps.
3. Preprocessing of multimodal datasets: We preprocess all multimodal brain images by using the Neuroglancer scripts. This involves transforming the MRI template into the pyramid pattern and organizing preprocessed digital sections into a tree folder structure suitable to effective data presentation. To preprocess 3D parcellation maps, conversion into surface grid files using the SPM in MATLAB is essential.
4. Composition of the interactive interface: We use the VUE framework to build the dynamic front-end, and responsive data binding is implemented via APIs to enable the interactivities. Interactive brain maps are presented on the front-end interface and align with URL parameters. Modification of the URL address segment occurs upon switching slice display.
5. Synergistic exhibition of datasets: The focal point of the exhibition is the MRI template, featuring three distinct panels displaying coronal, horizontal, and sagittal views, alongside a panel presenting the 3D surface model. Partitioned maps are registered to the MRI template within the Neuroglancer tool using a transform matrix accounting for rotation and translation. Histological images selected from the plug-in components are visually positioned within the MRI template.
6. Display of digital sections: To expedite loading and optimize display, the system renders only the user-selected section while preventing simultaneous loading of other sections. Compressed images with a resolution of 1 μm/pixel are offered instead of the original resolution to enhance performance.

## Acknowledgments

We thank the support from the National Natural Sci-ence Foundation of China 32350410397;Science,Technology,Innovation Commission of Shenzhen Municipality,JCYJ20220530143014032,JCYJ20230807113017035,KCXFZ20211020163813019, WDZC20200820173710001,Shenzhen Medical Research Funds,D2301002;Depart ment of Chemical Engineering-iBHE special cooperation joint fund project, DC E-iBHE-2022-3; Tsinghua Shenzhen Interna-tional Graduate School Cross-disciplinary Research and Innovation Fund Research Plan, JC2022009; and Bureau of Planning, Land and Resources of Shenzhen Municipality (2022) 207.

## Declaration of Competing Interest

The authors declare that they have no known competing financial interests or personal relationships that could have appeared to influence the work reported in this paper.

## Notes

### Competing Interest Statement

The authors have declared no competing interest.

