## Supplementary Figures for "Hypergraph Cortical Cytoarchitectonic Parcellation with Multimodal Canine Brain Atlas"

*Corresponding Author*

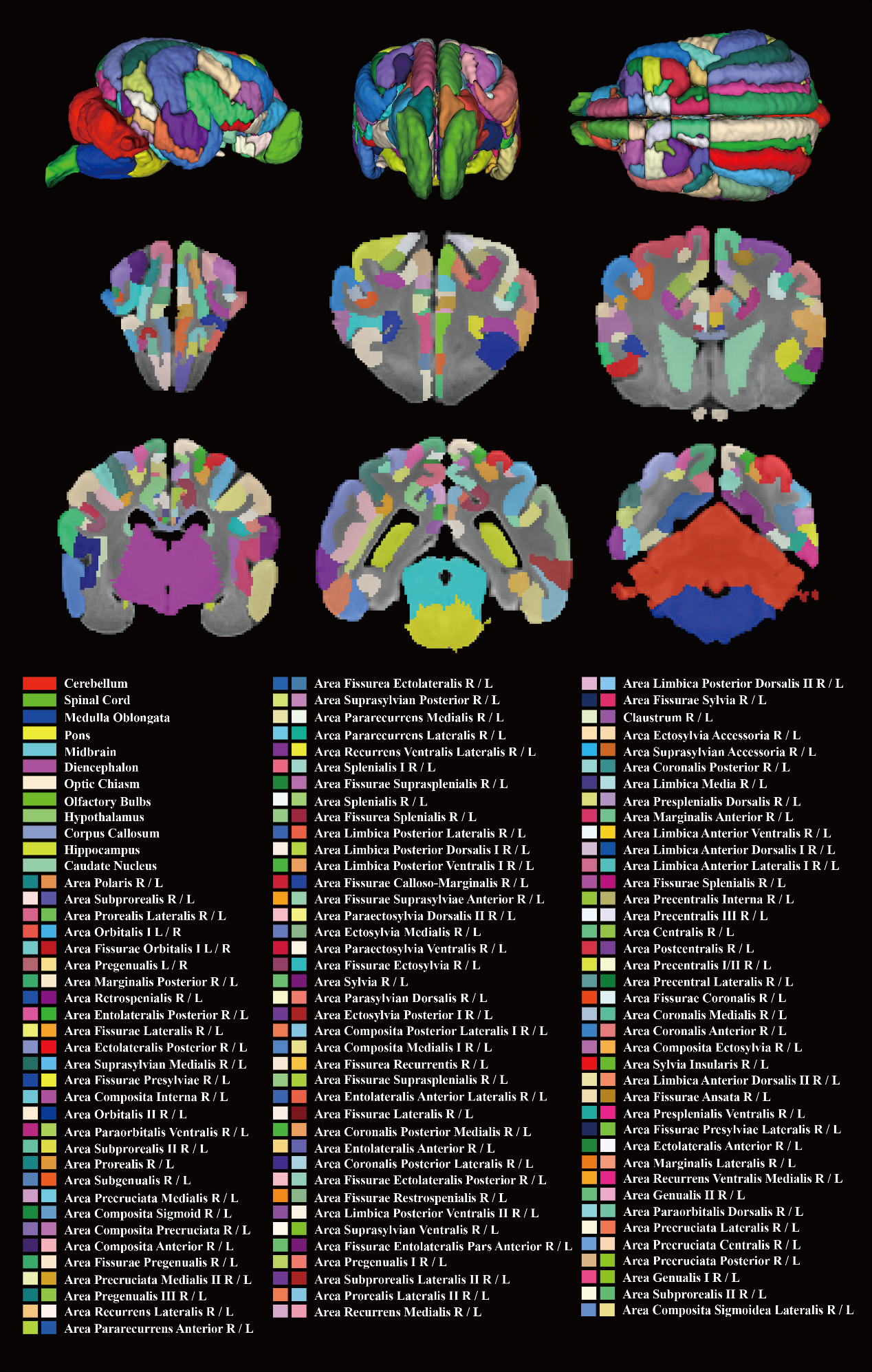

**Figure S1. Meyloarchitectonic cortical parcellation and anatomic delineation of the Kunming canine MRI.** The reconstructed 3D model within the parcellations is shown in three views, including the right lateral, rostral, and dorsal. We label all subregions with different colors and note the corresponding structural names. This map is offered as one mode of the interactive atlas.

Figure S2-16 show the H&E stained brain slices along the rostral-to-caudal axis and delineations of microscopic neuroanatomy and myeloarchitecture for individual slices.

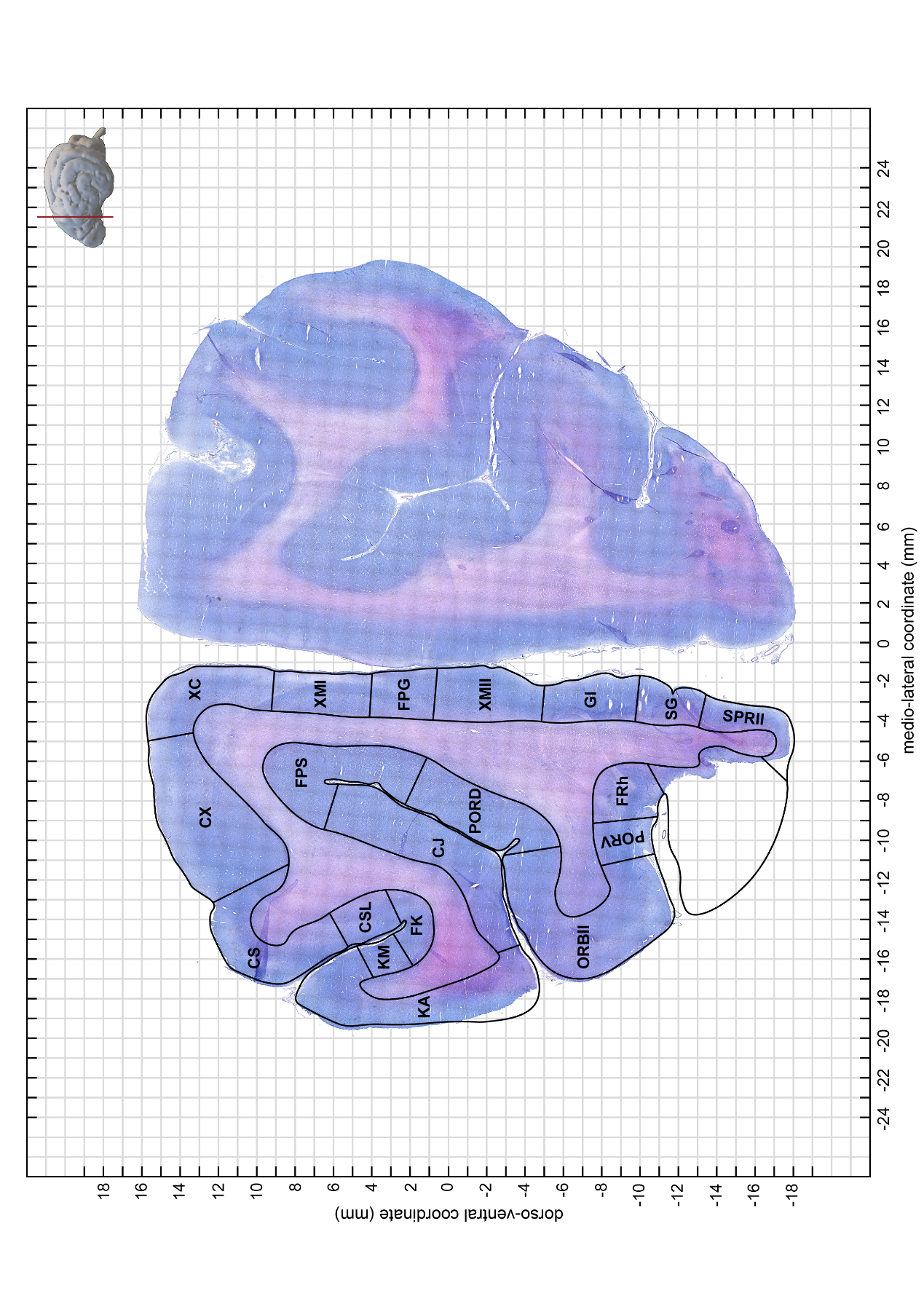

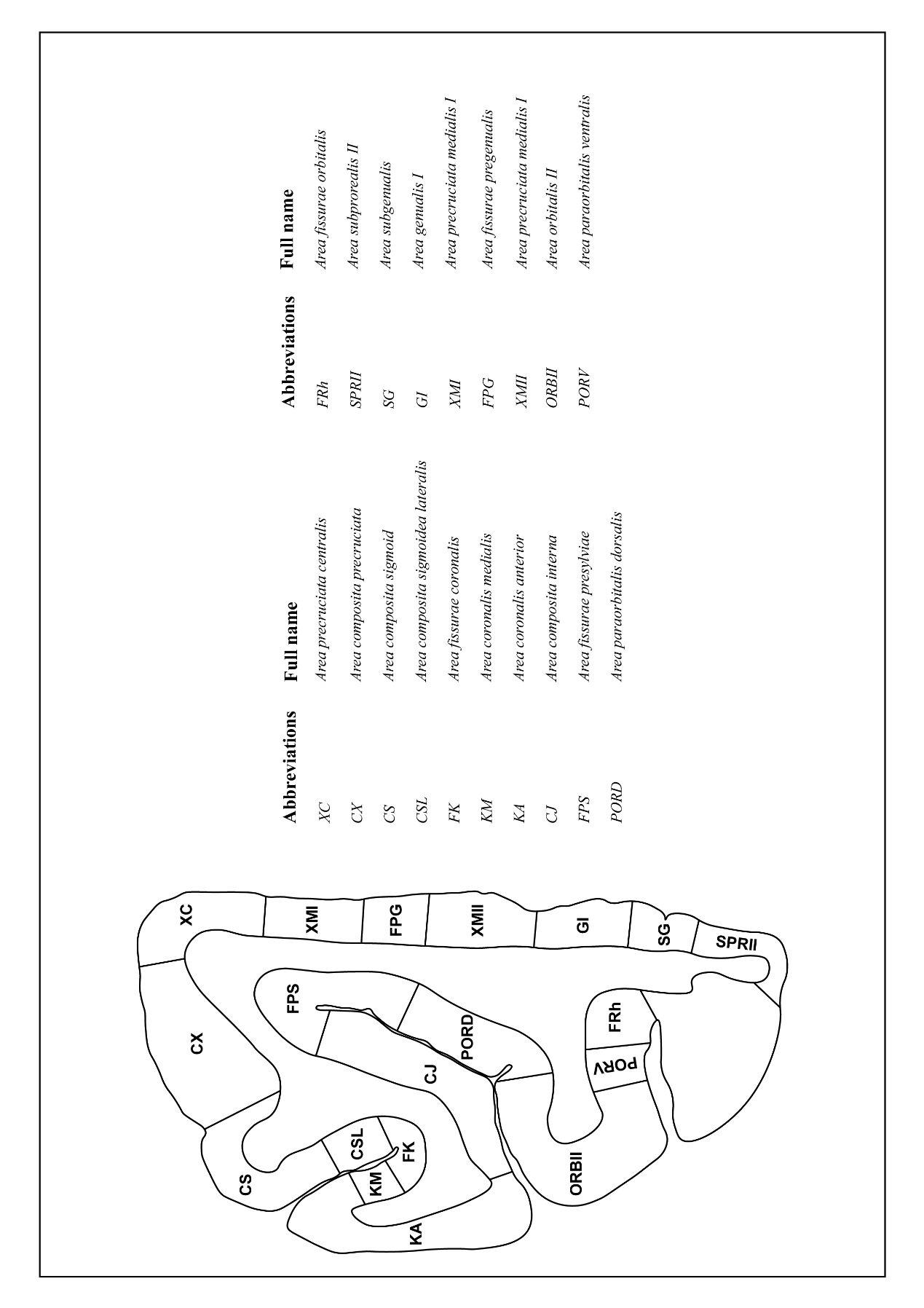

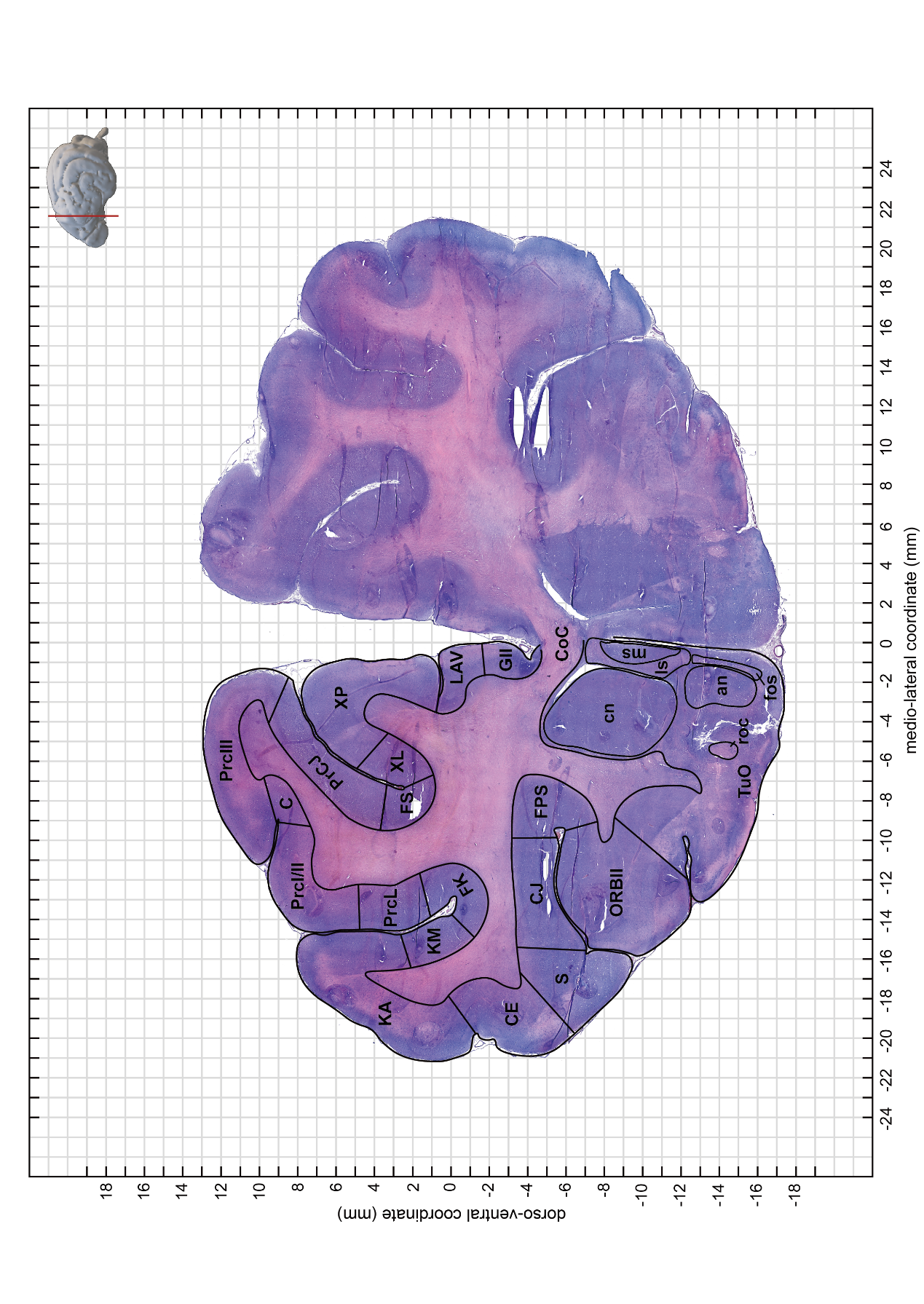

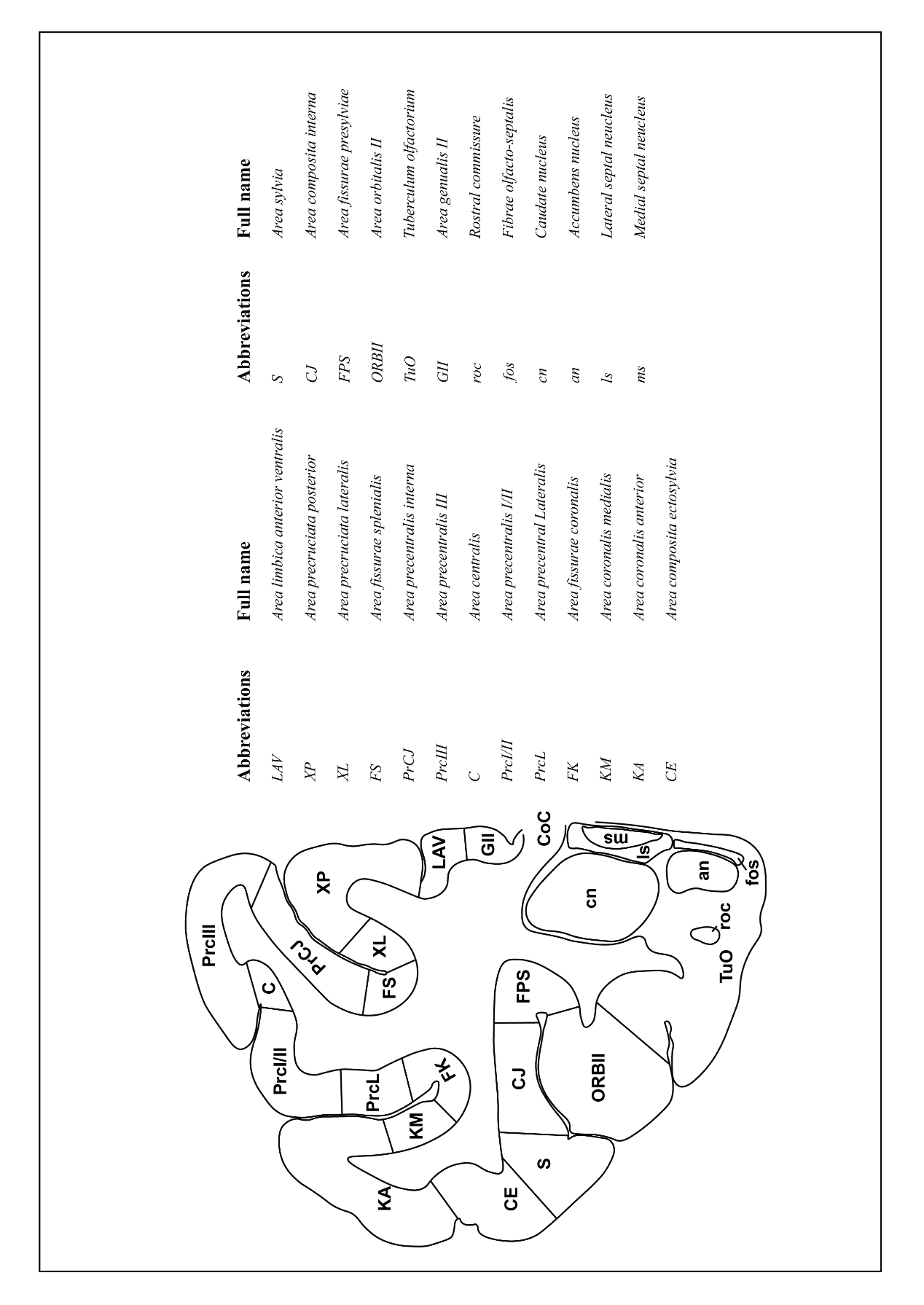

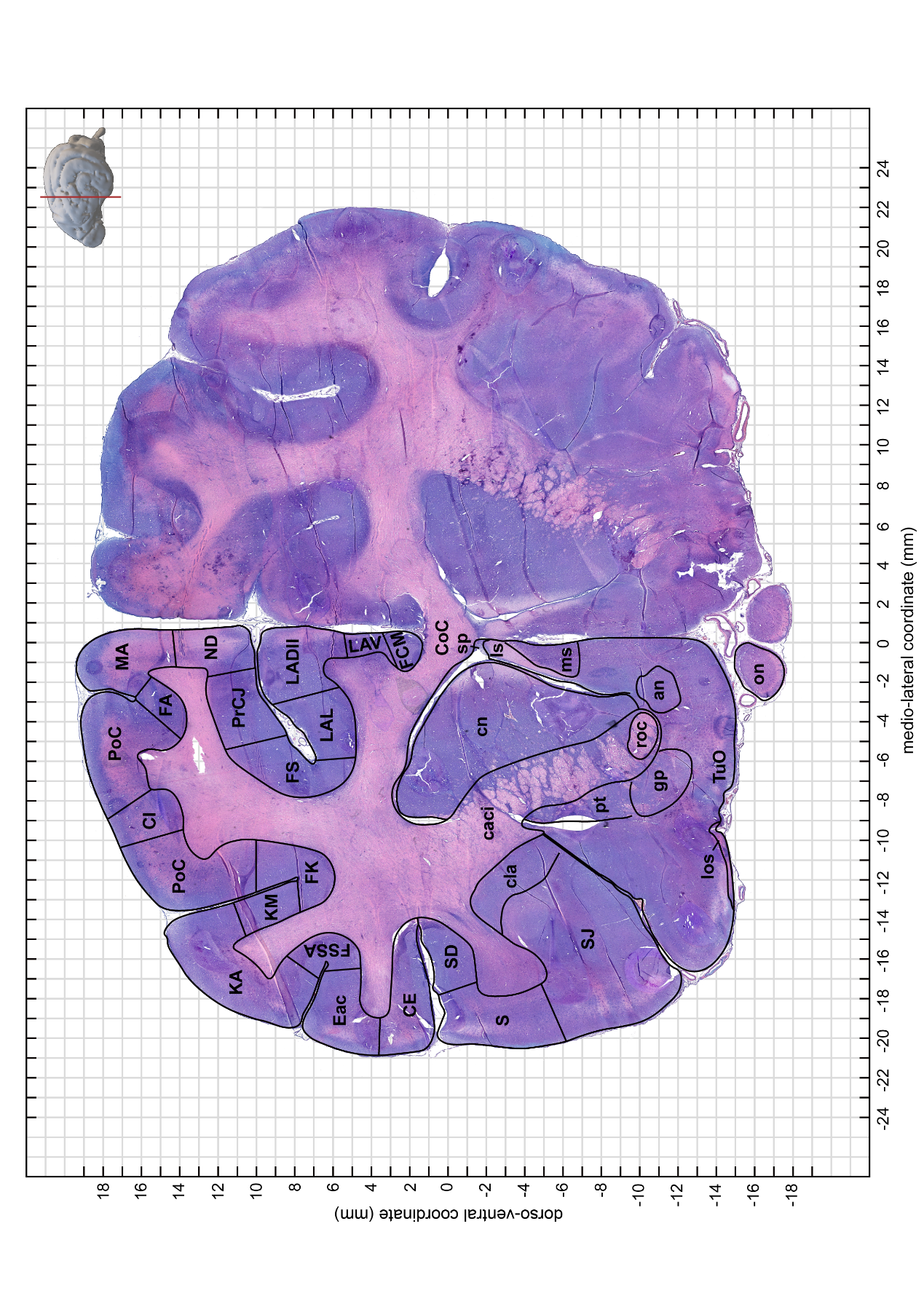

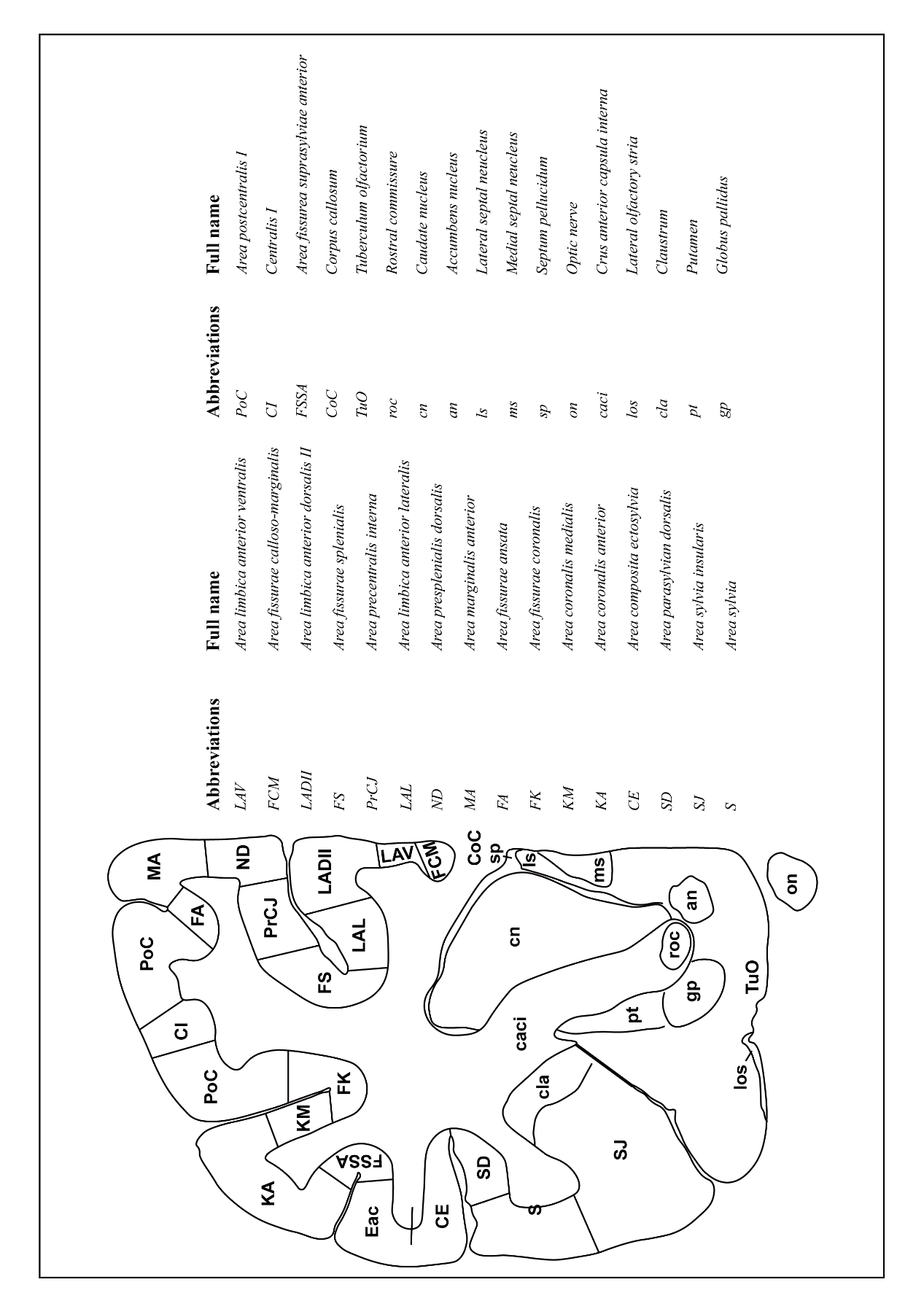

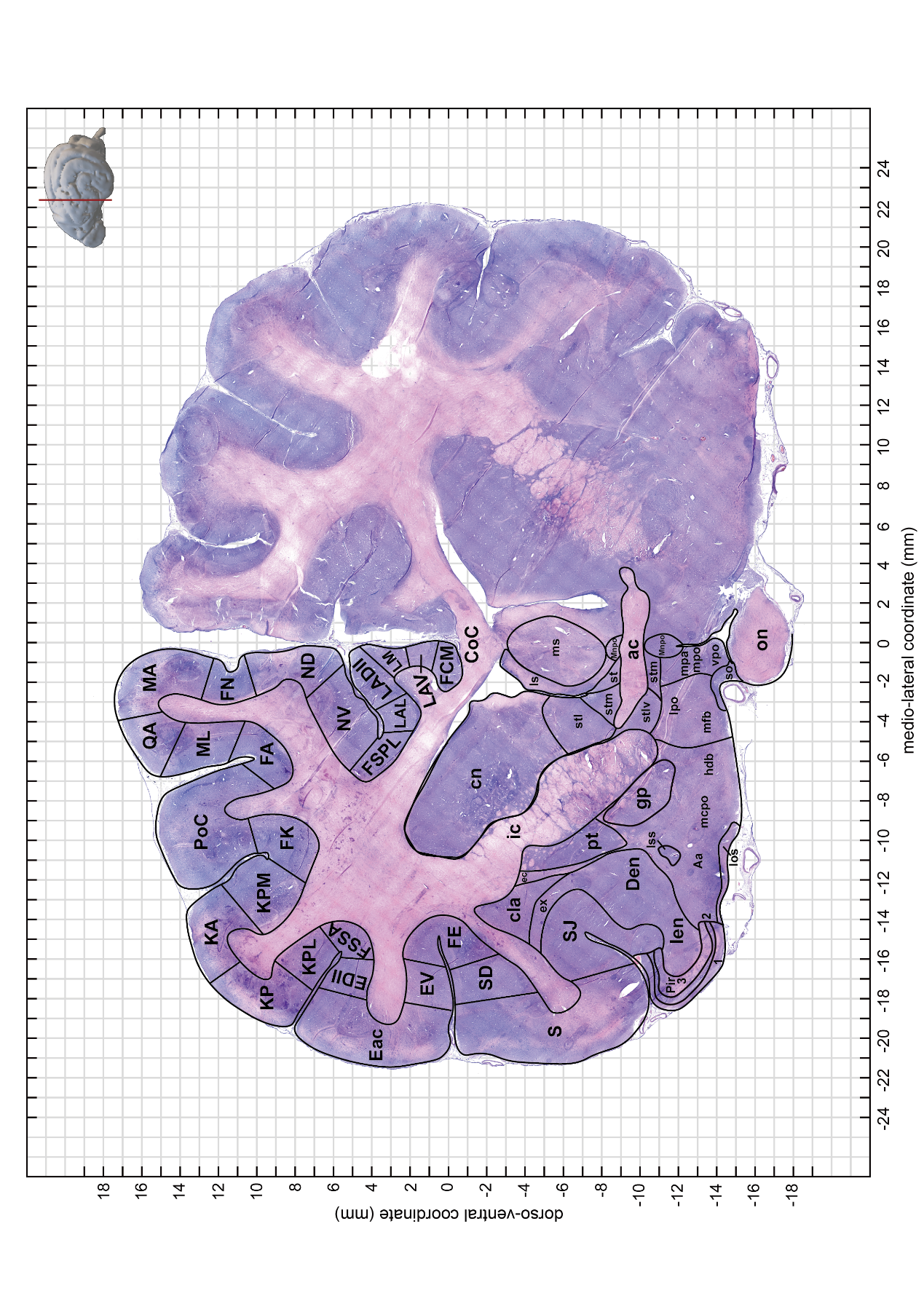

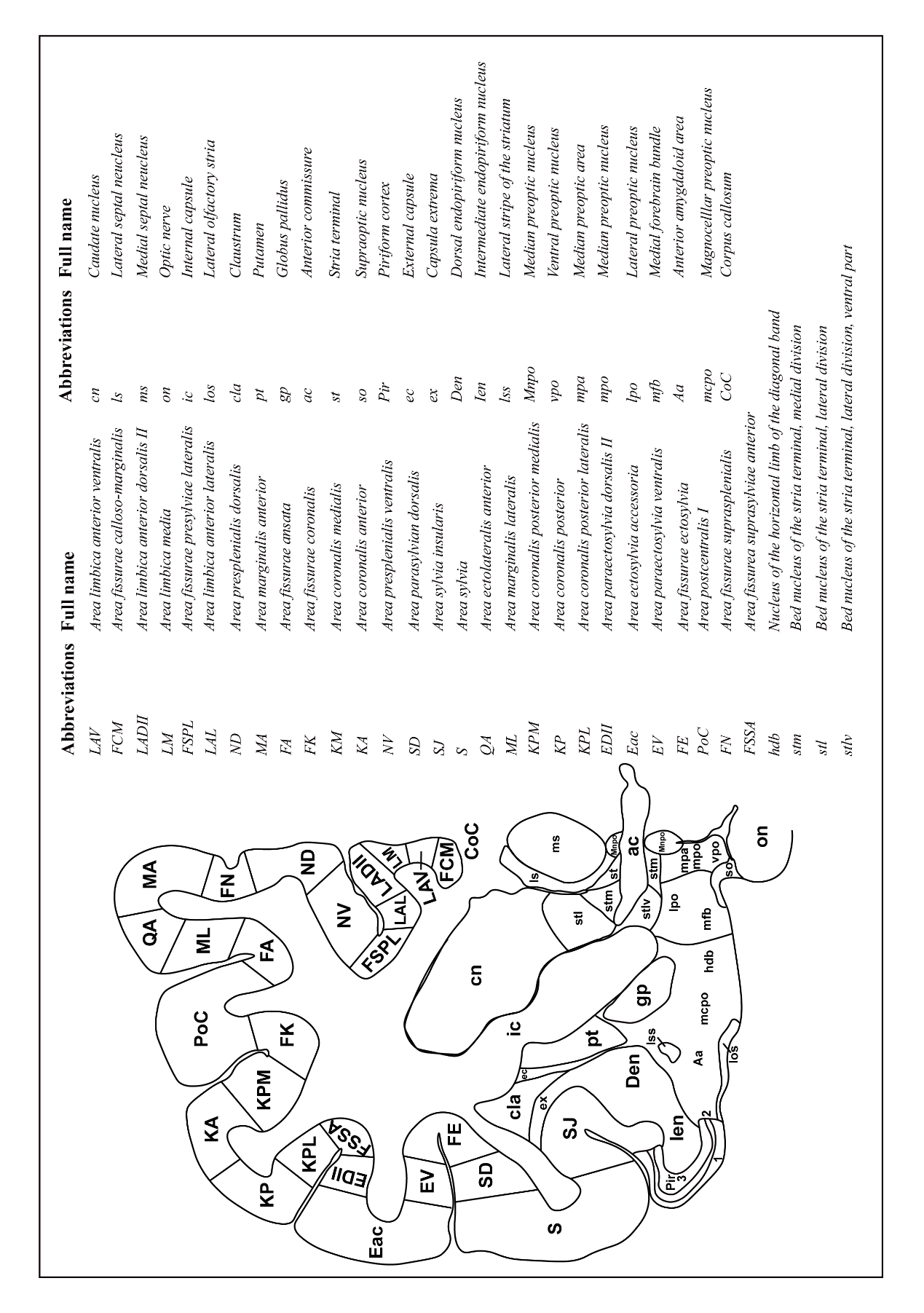

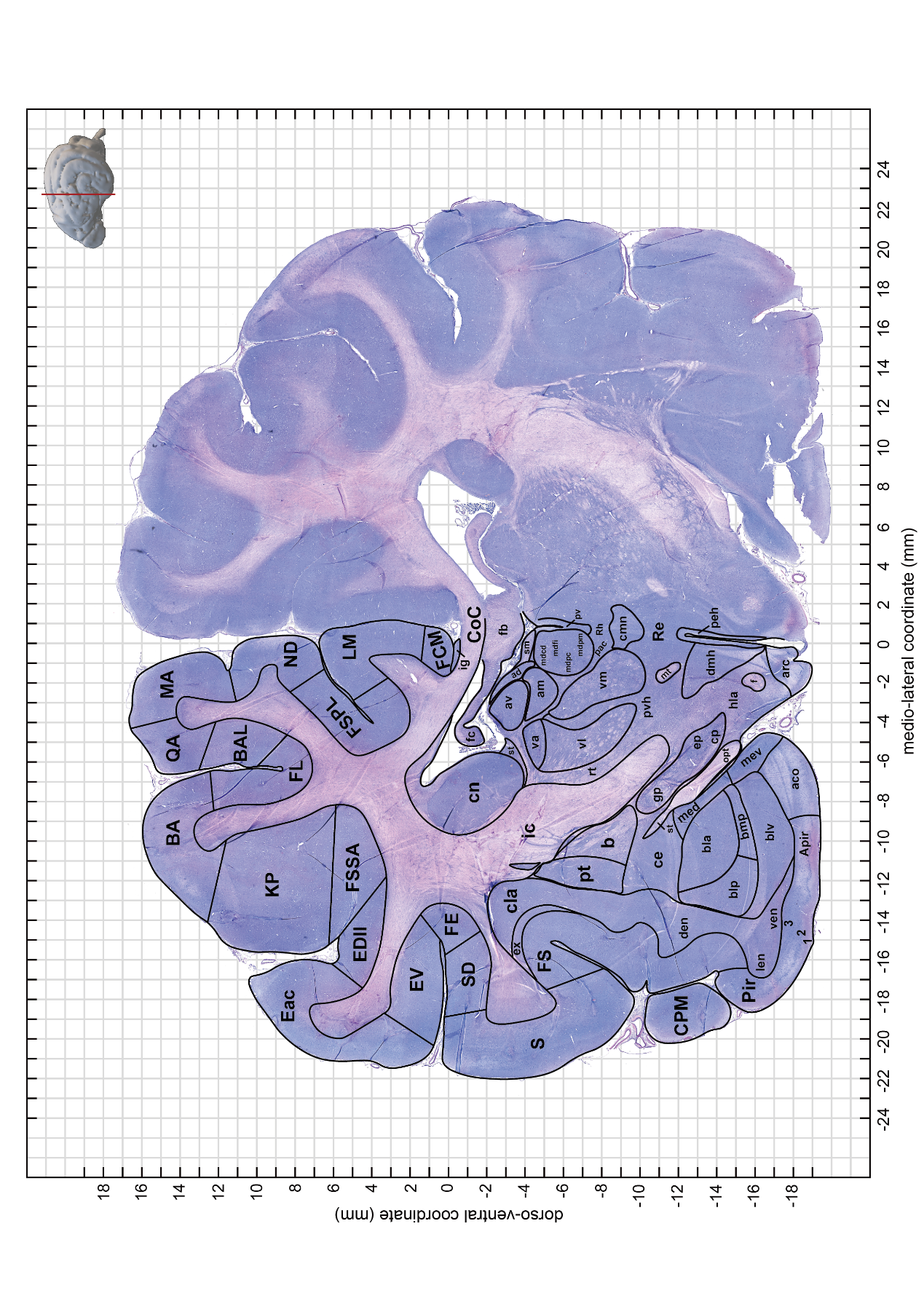

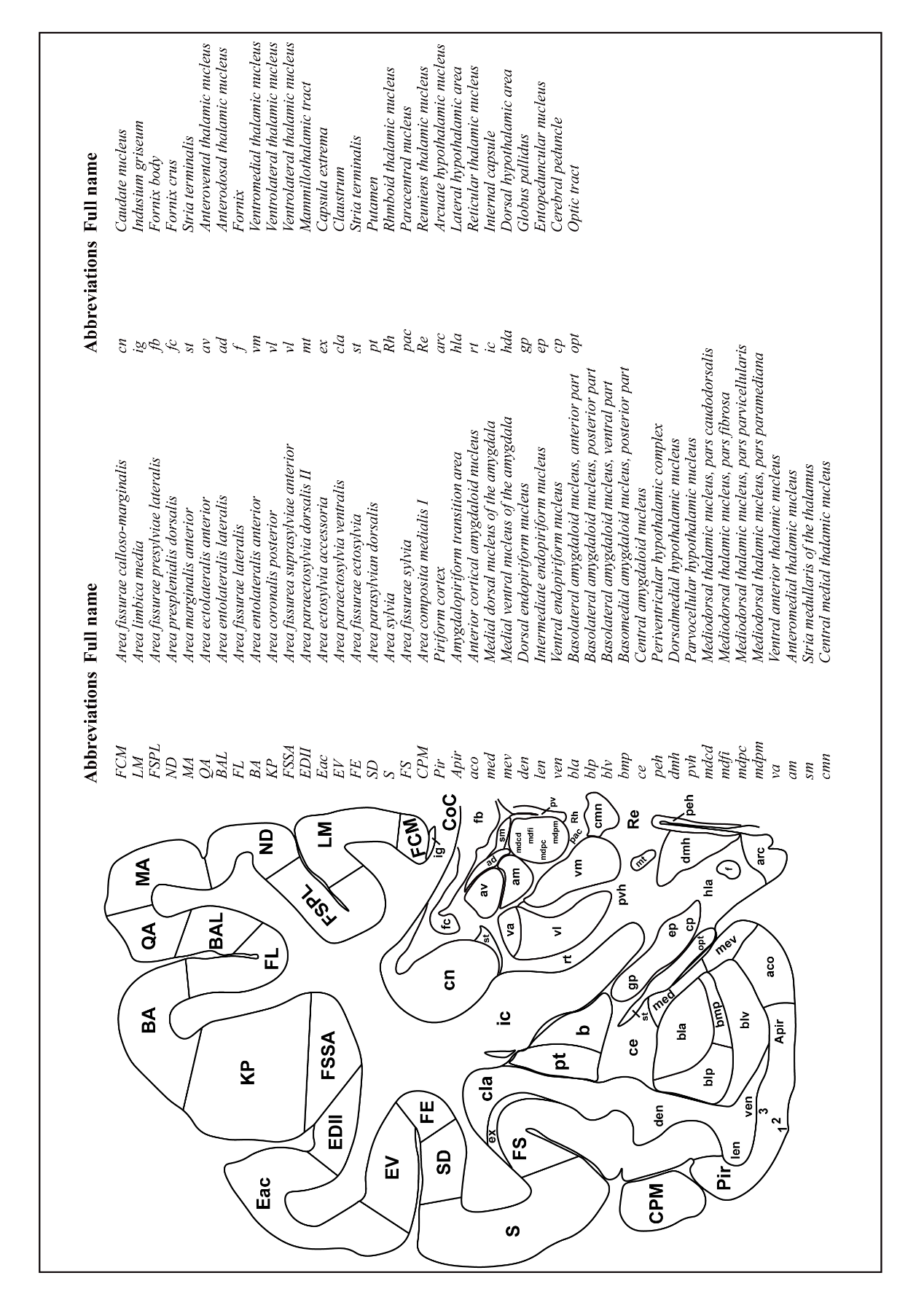

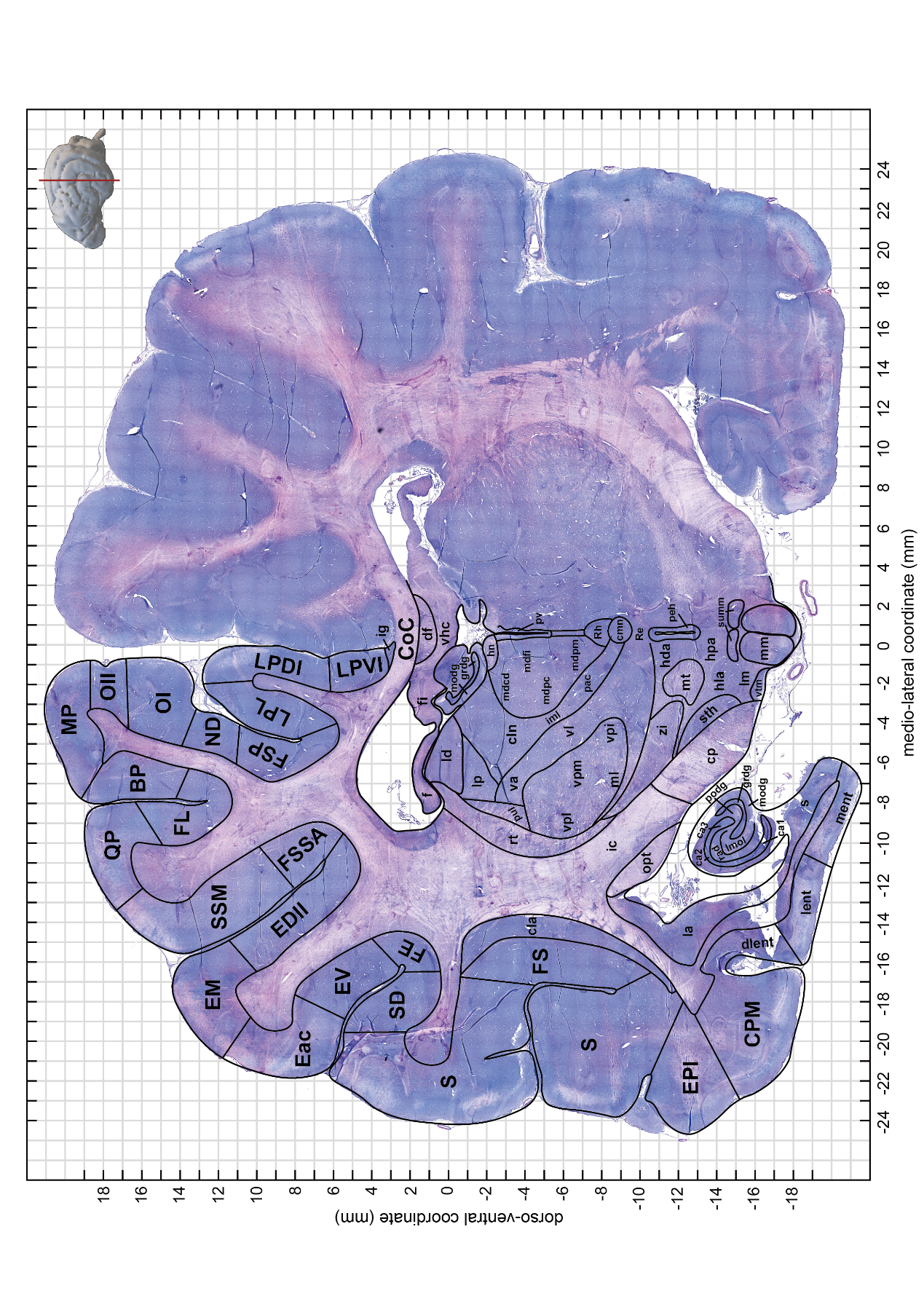

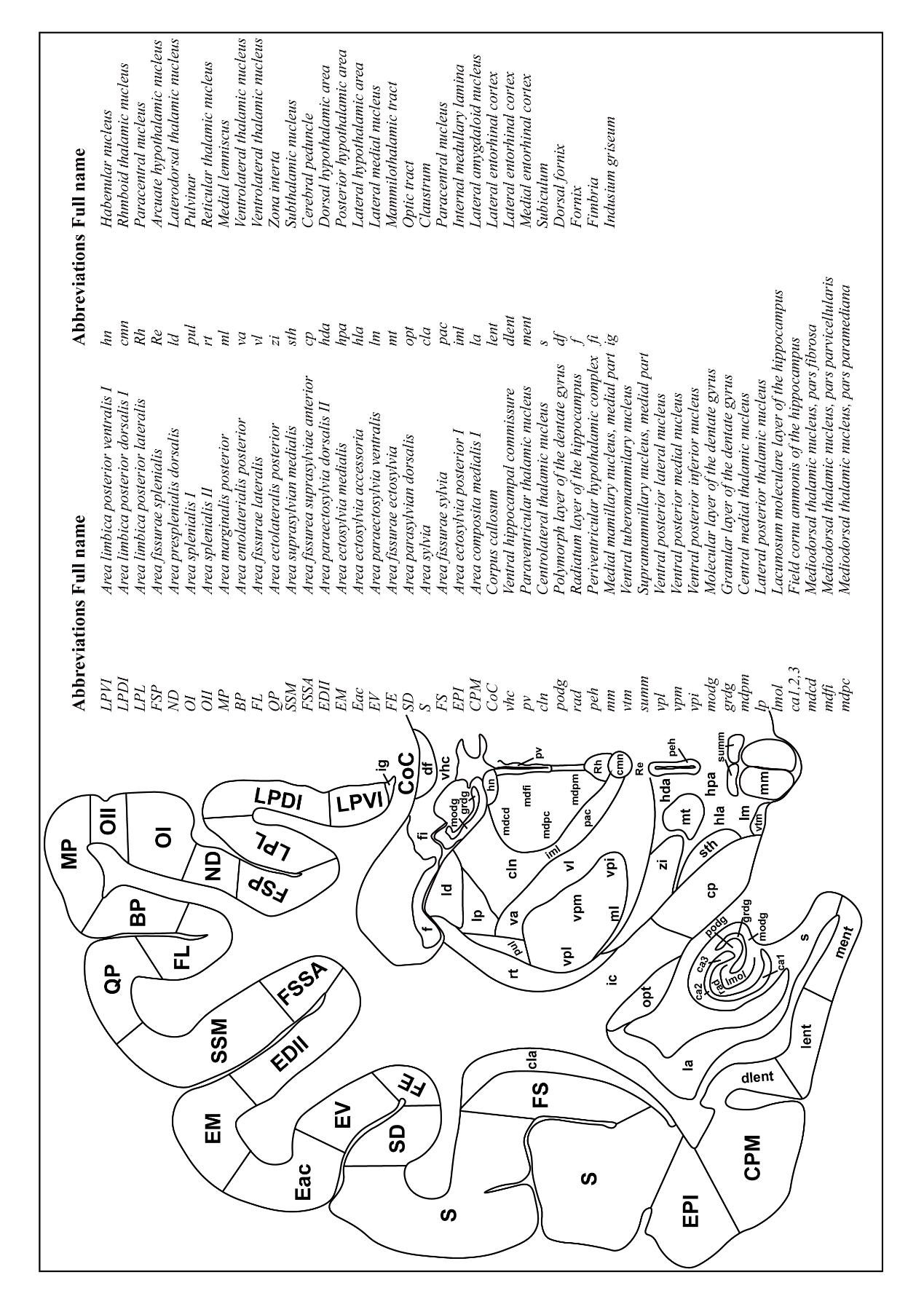

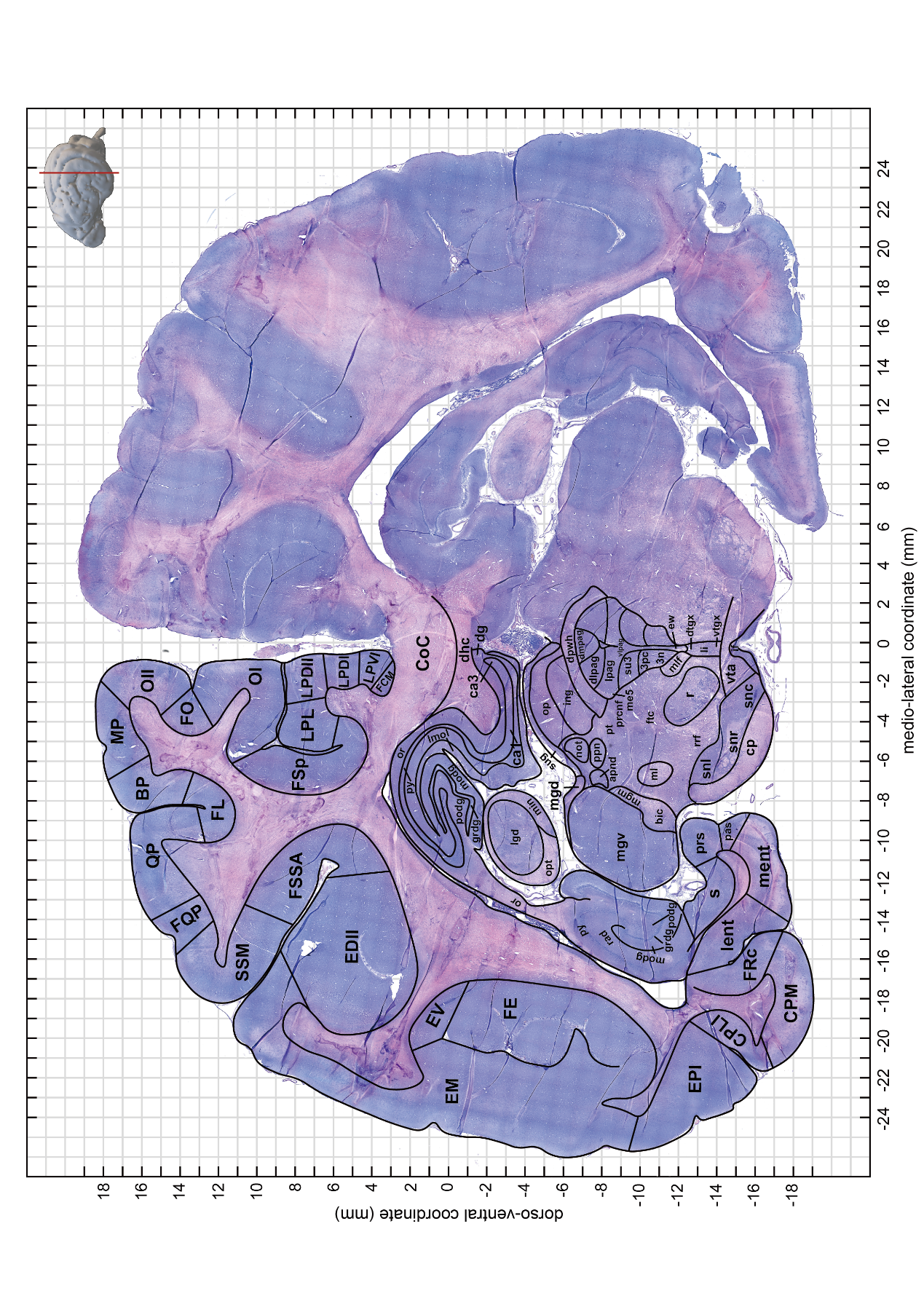

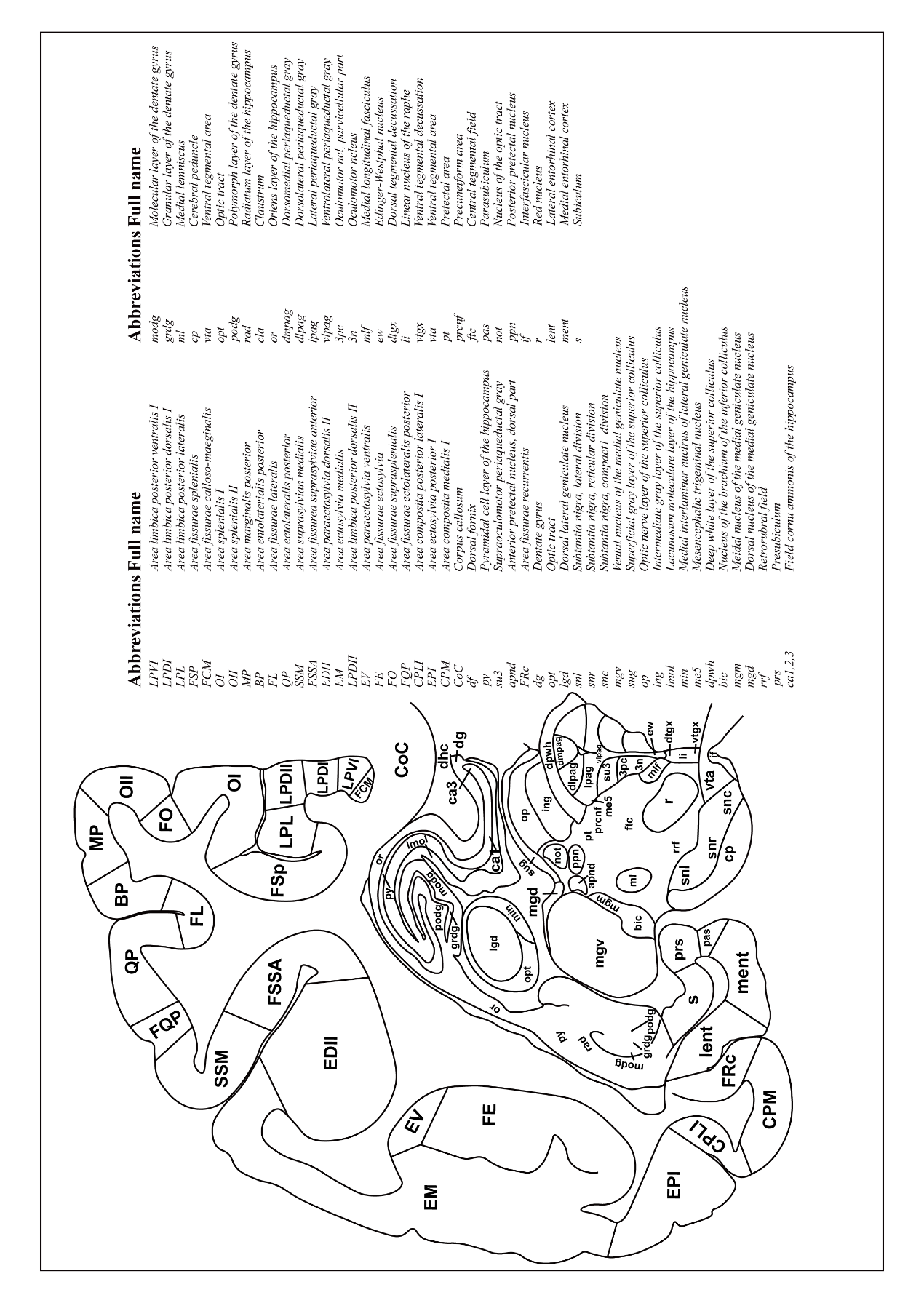

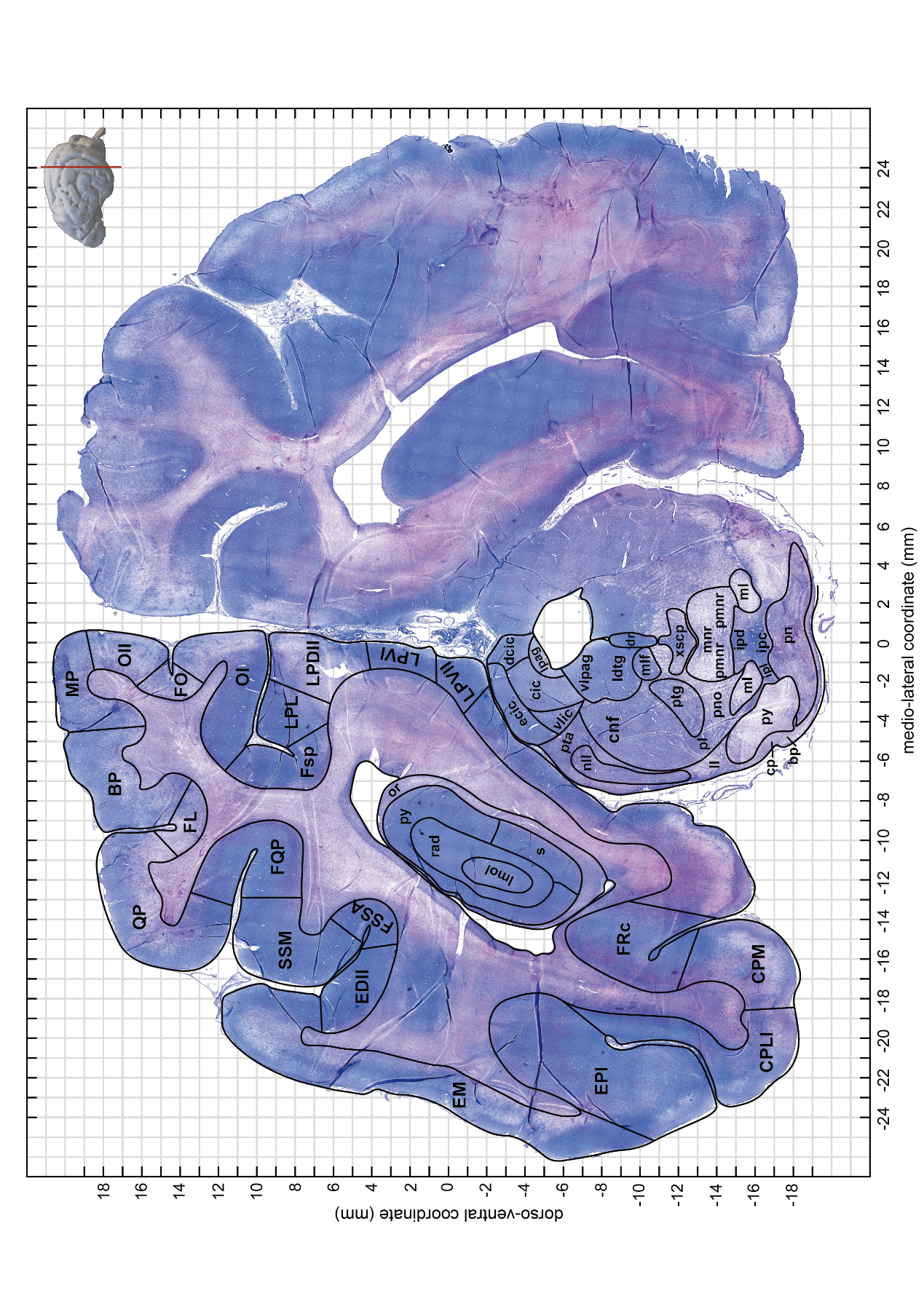

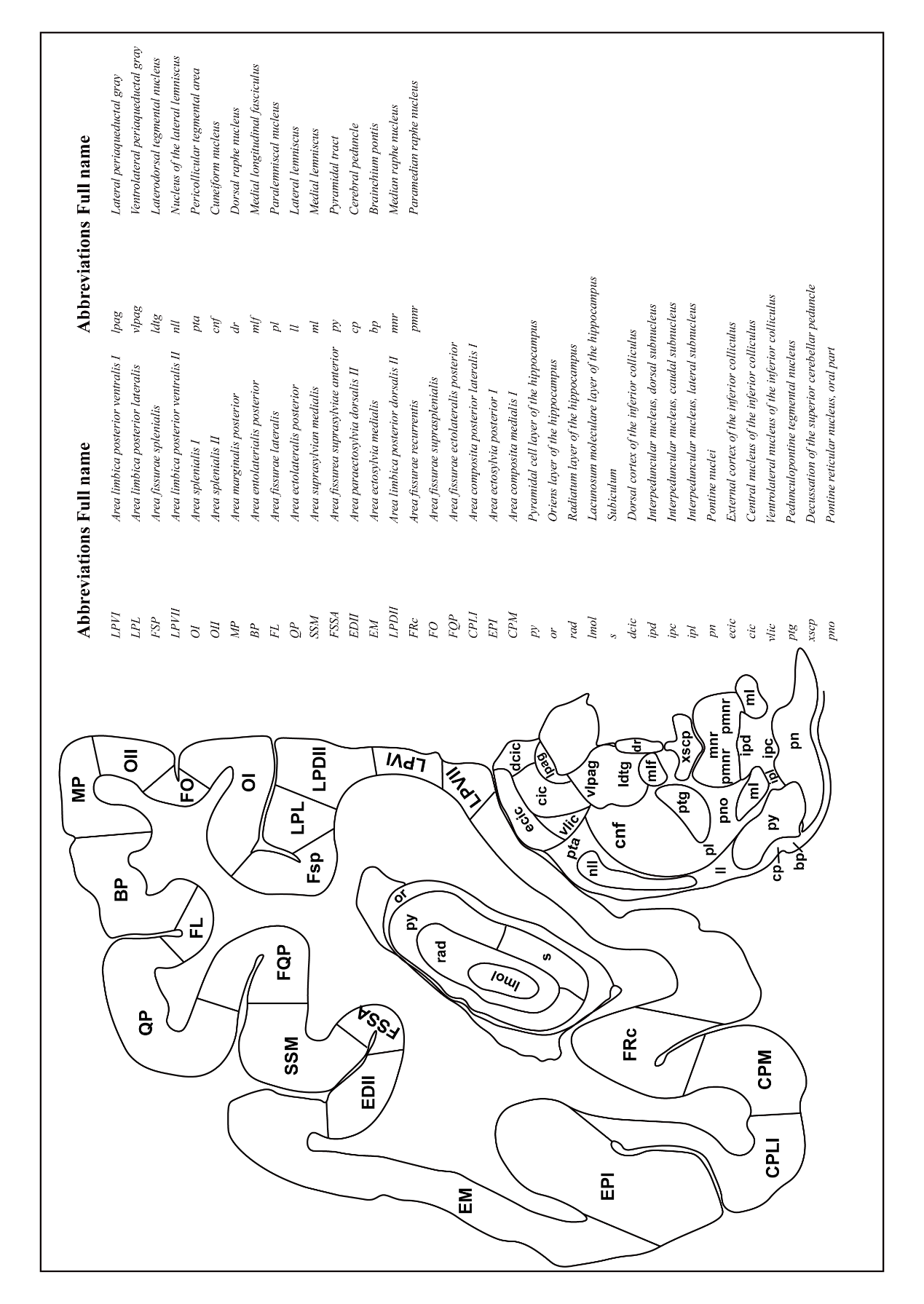

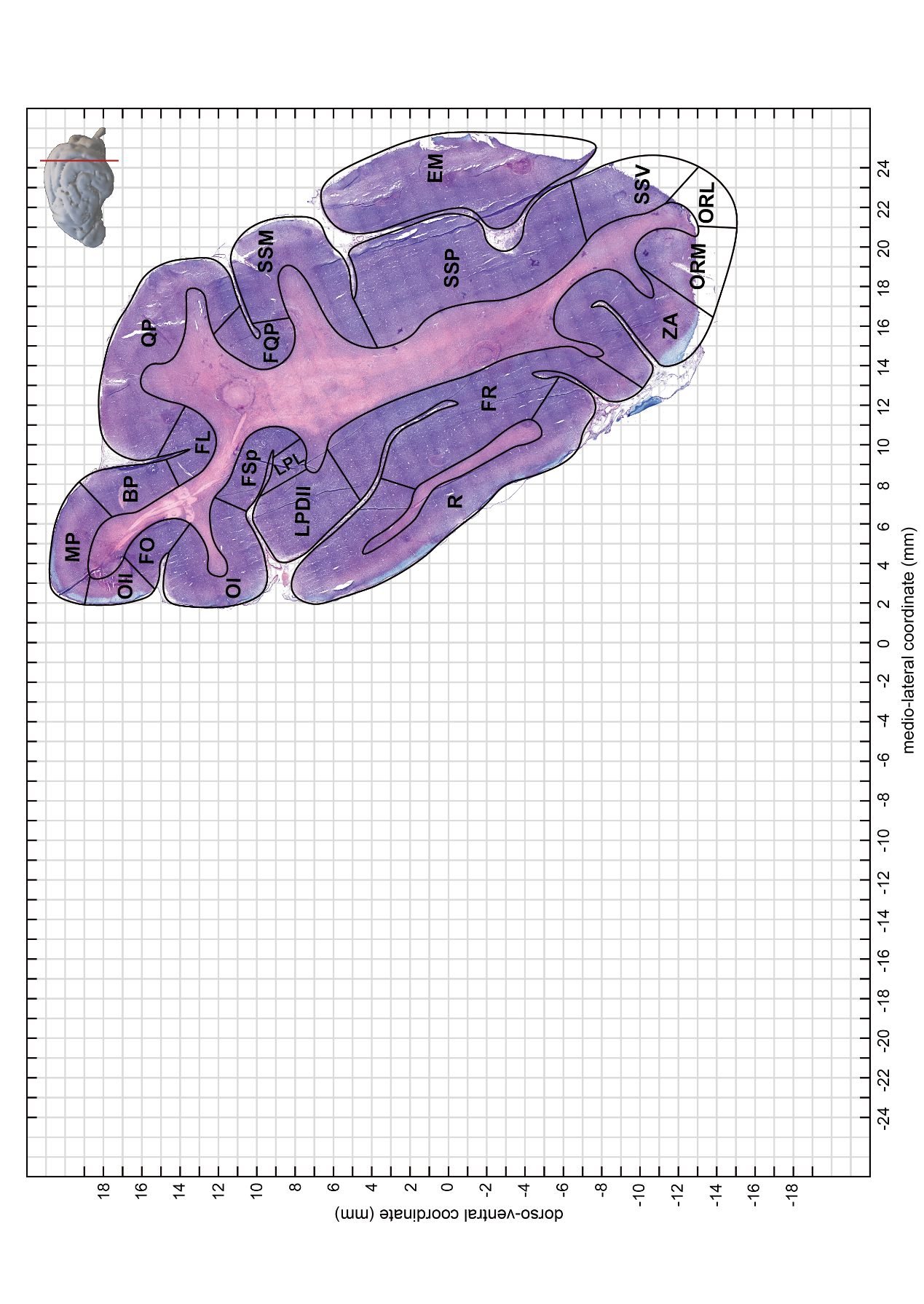

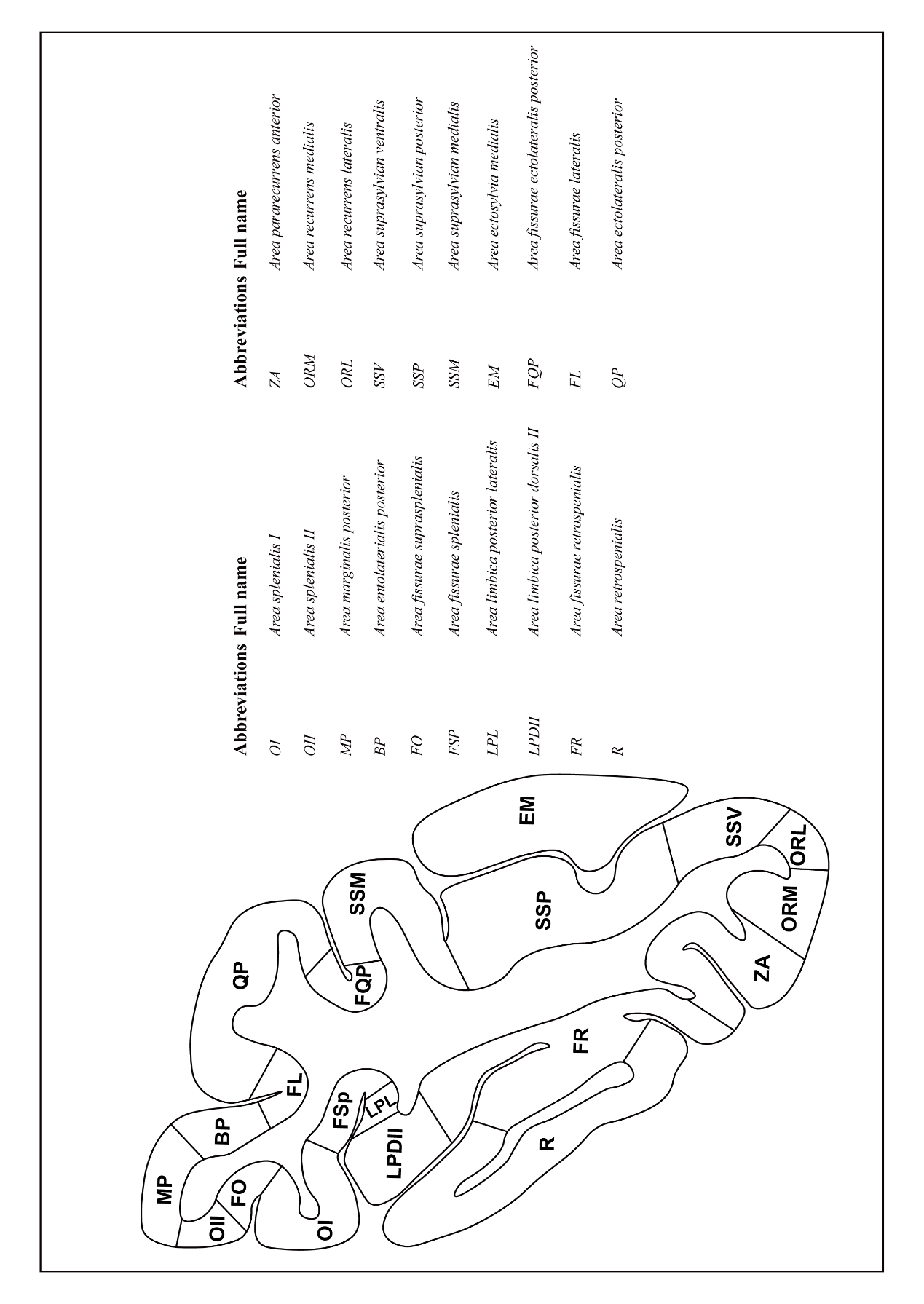

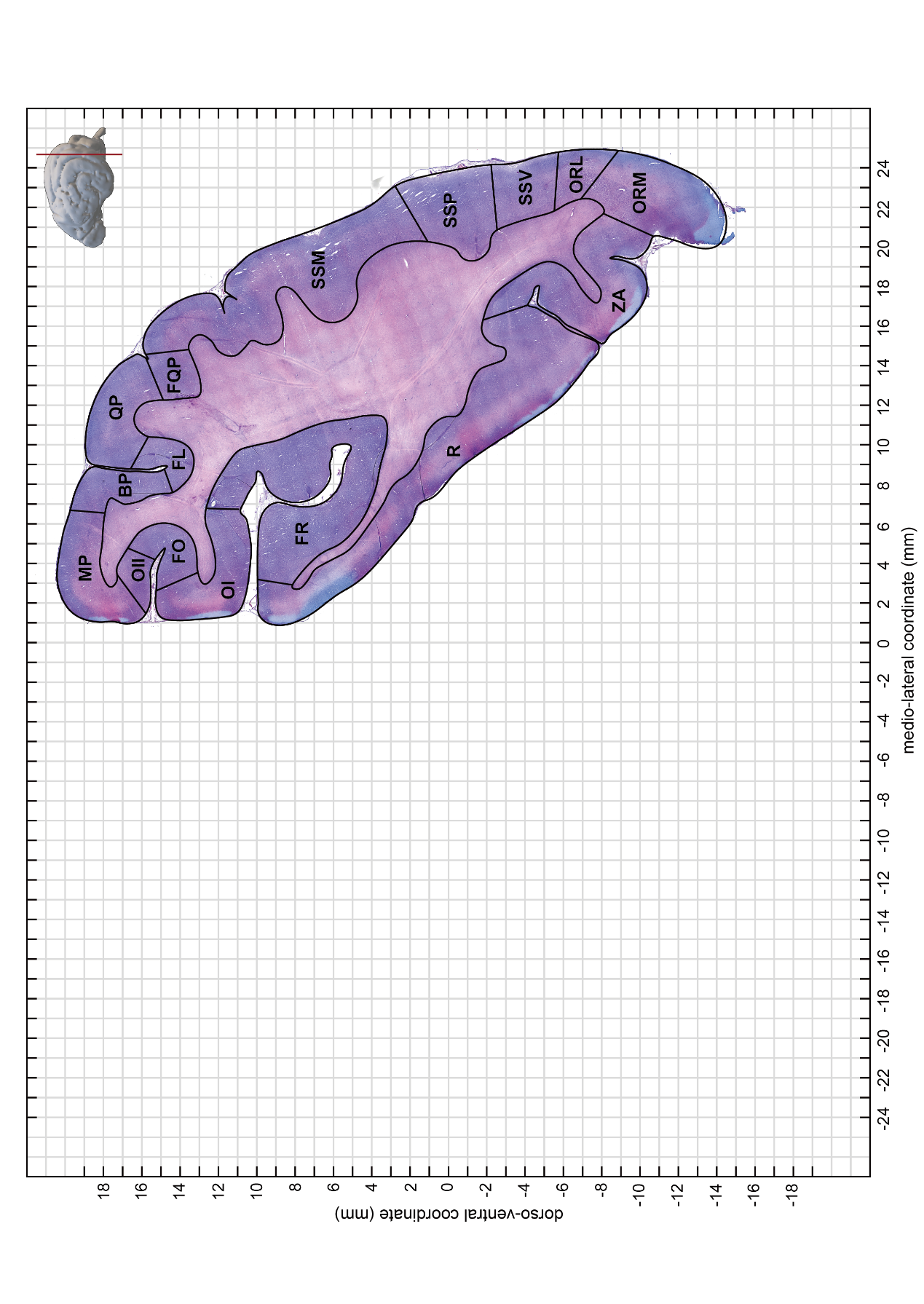

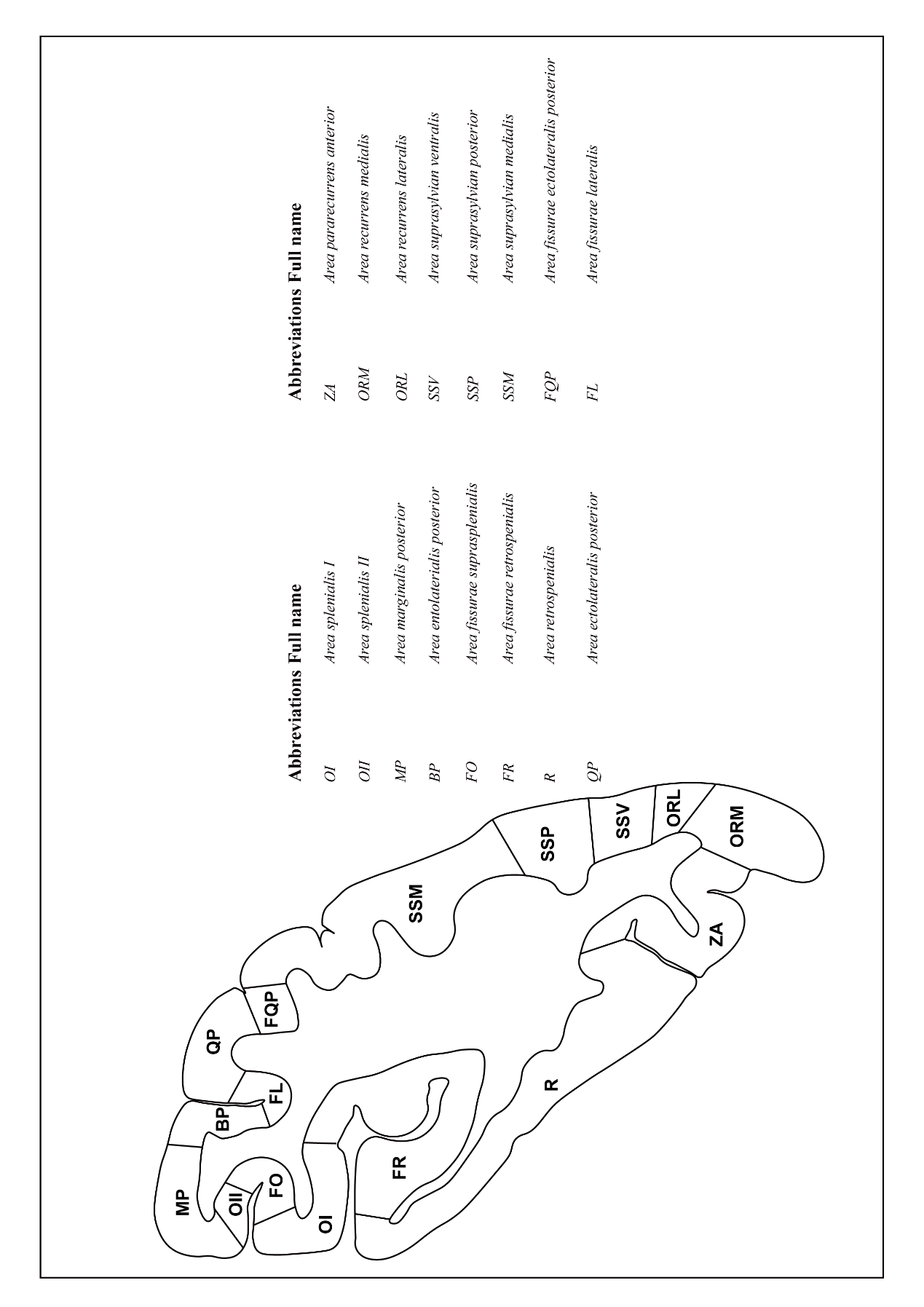

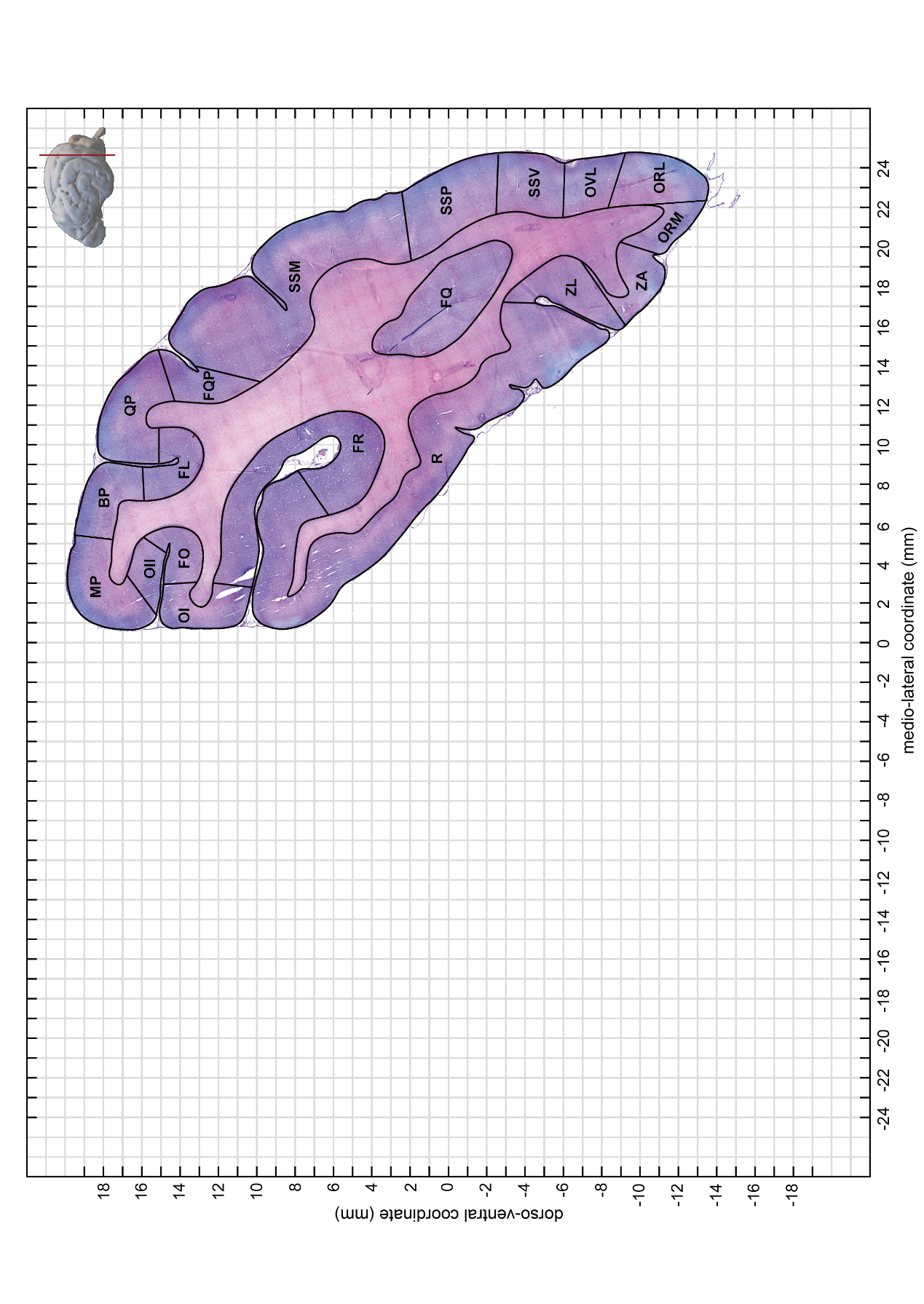

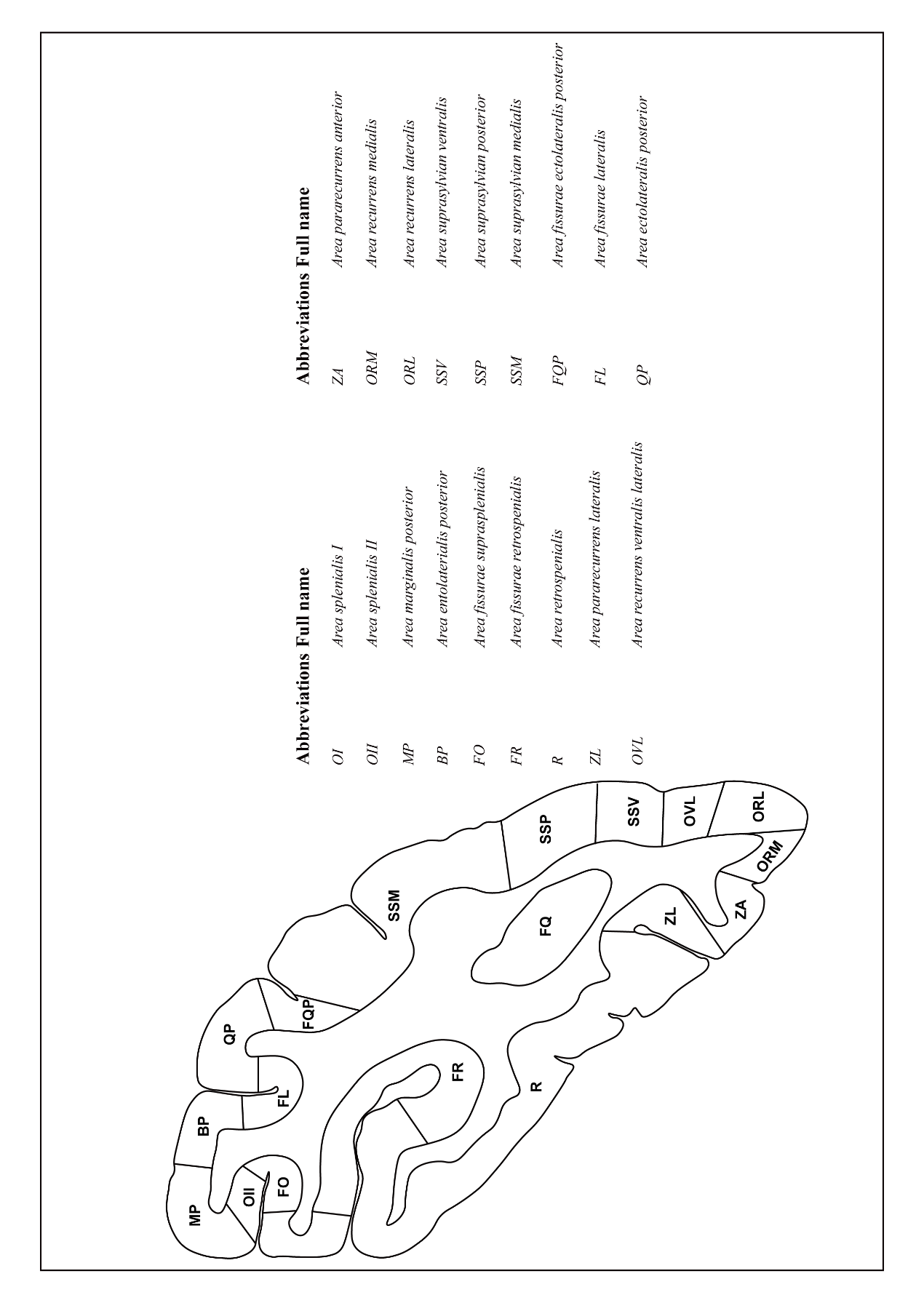

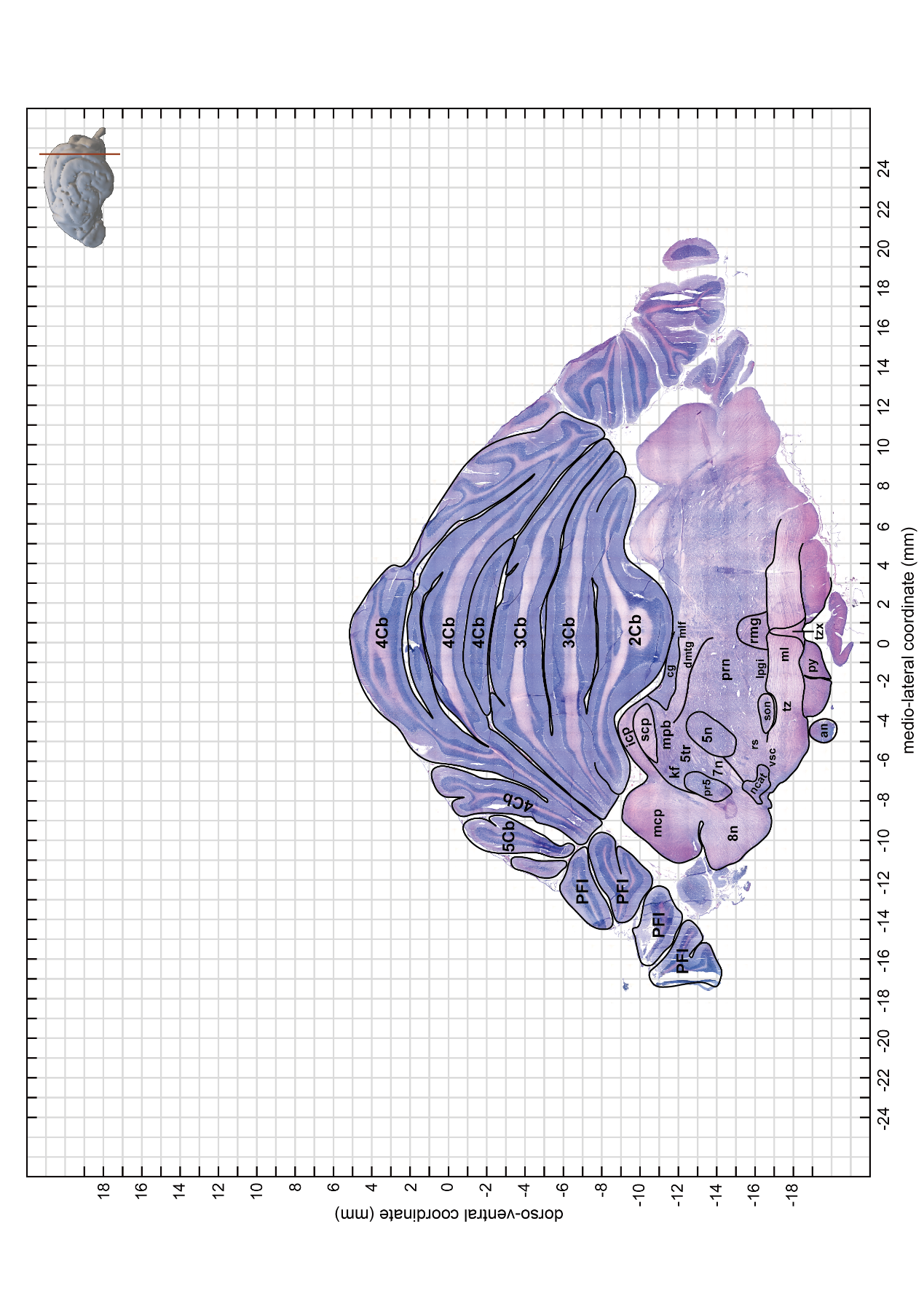

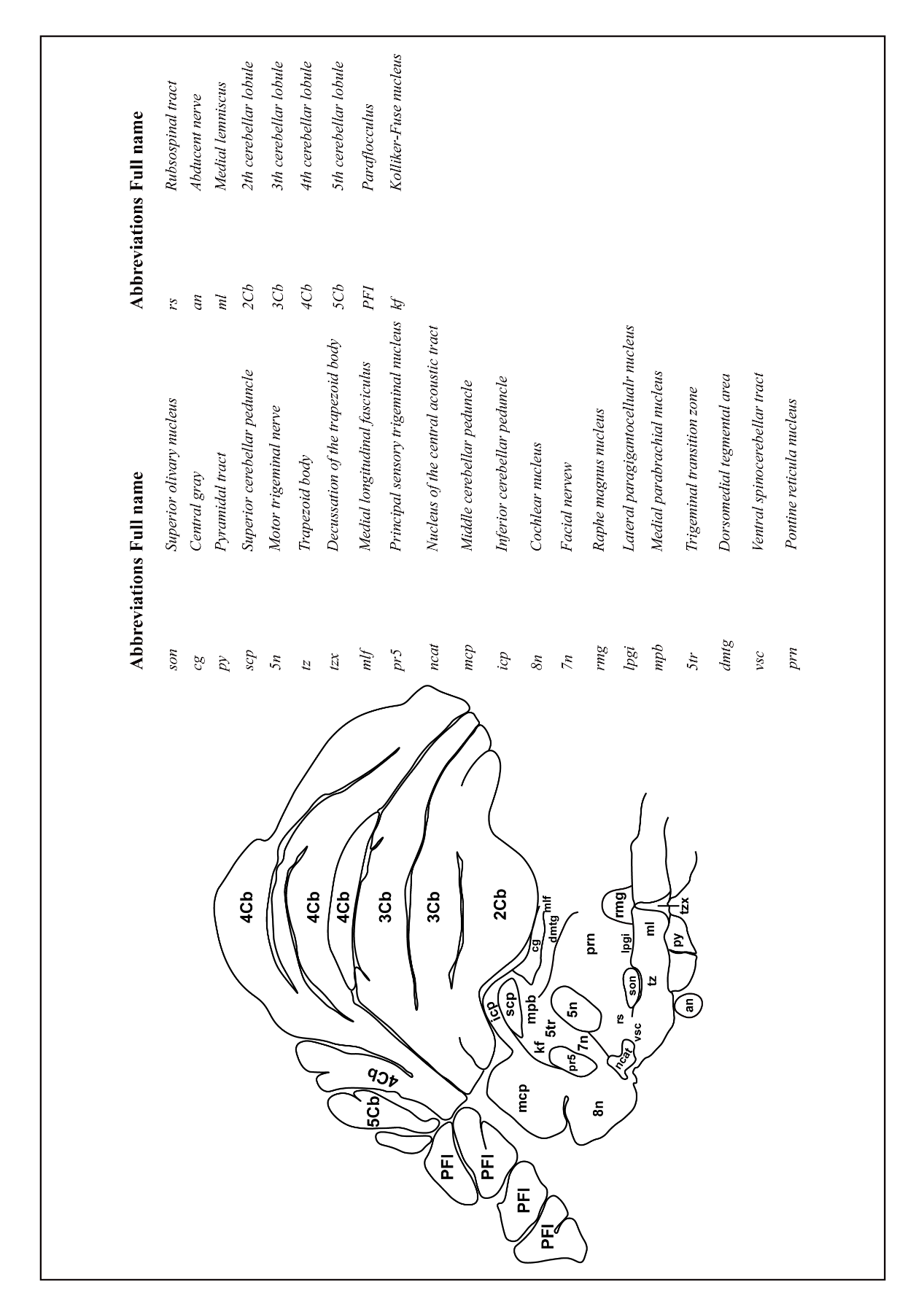

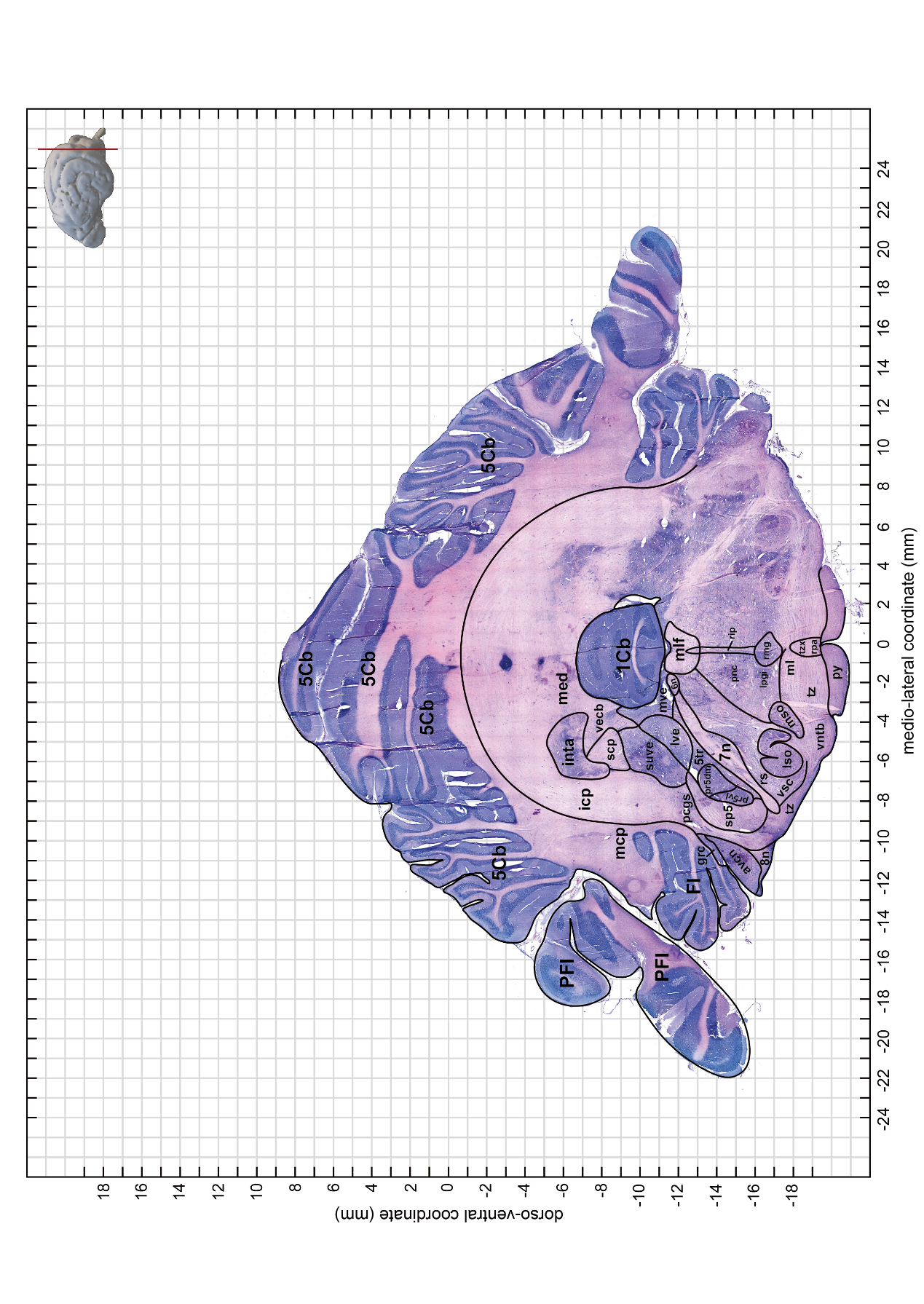

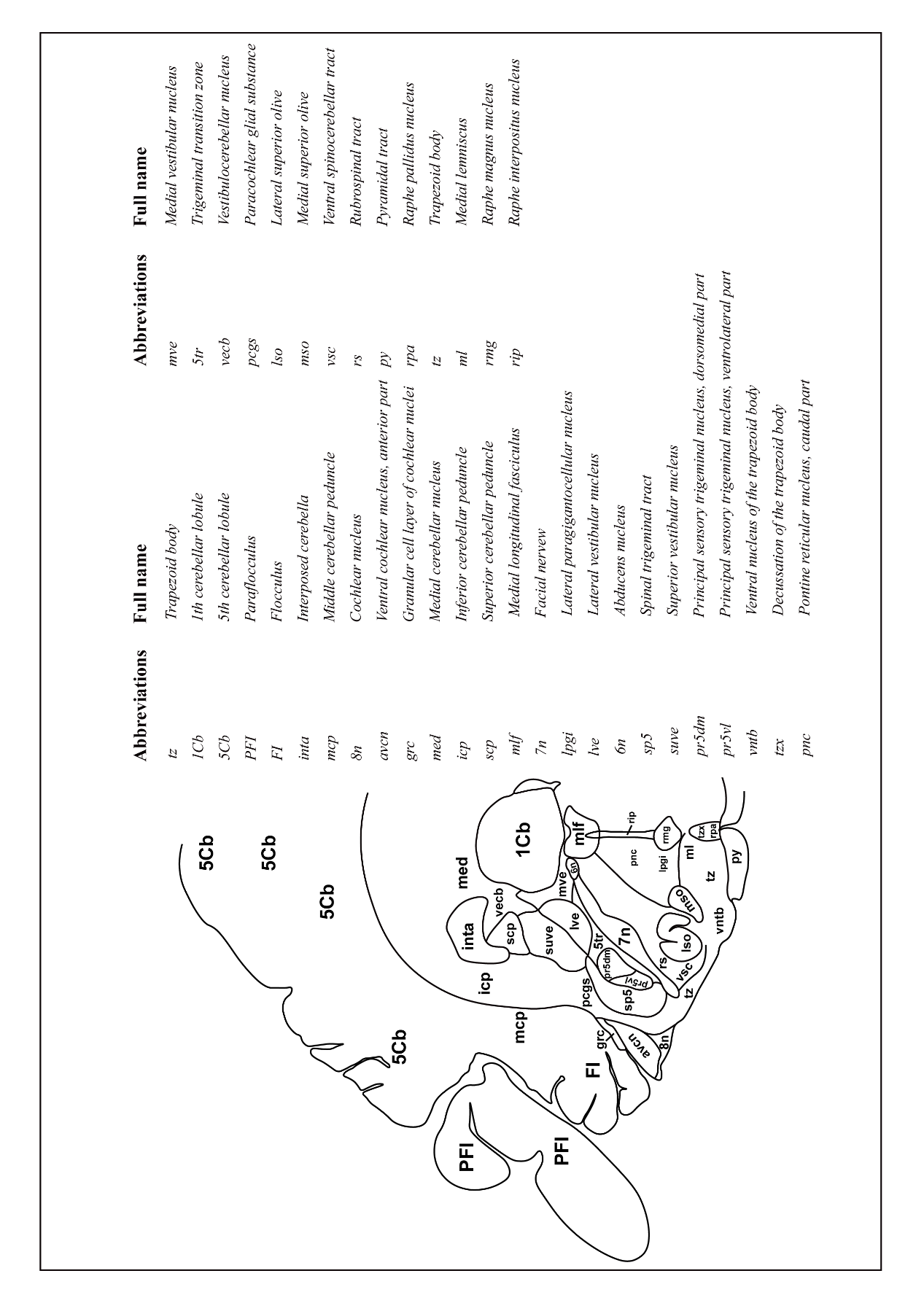

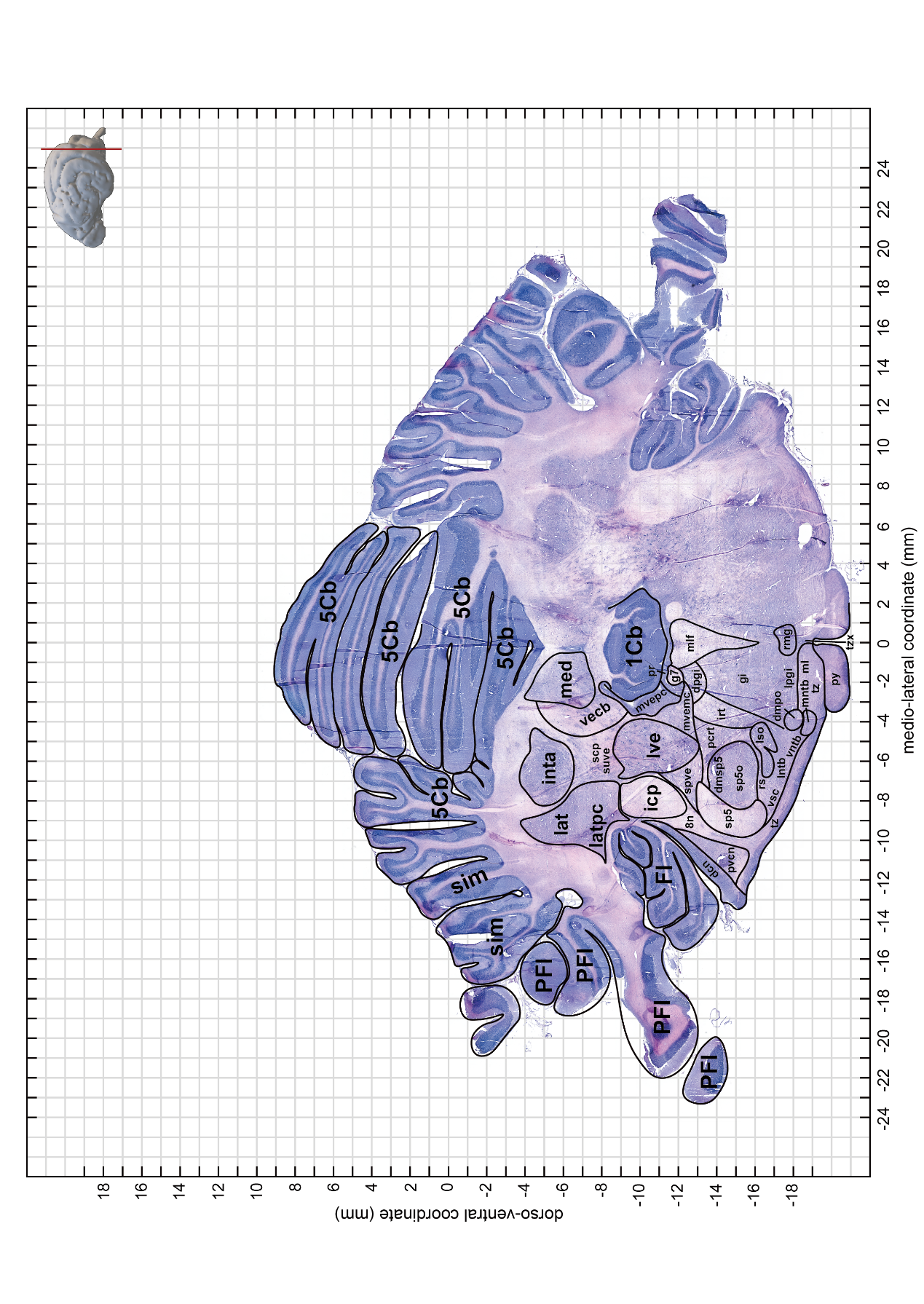

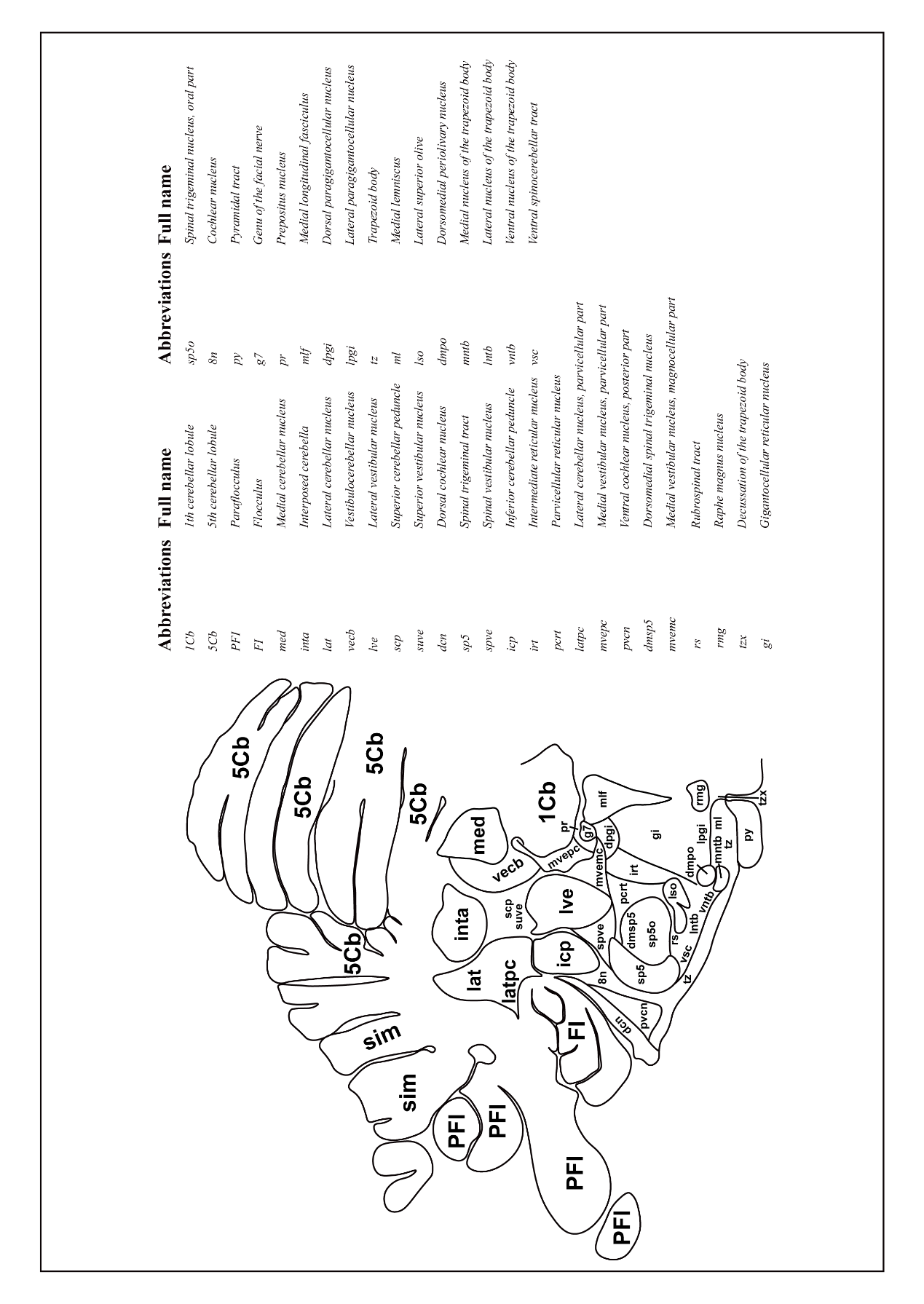

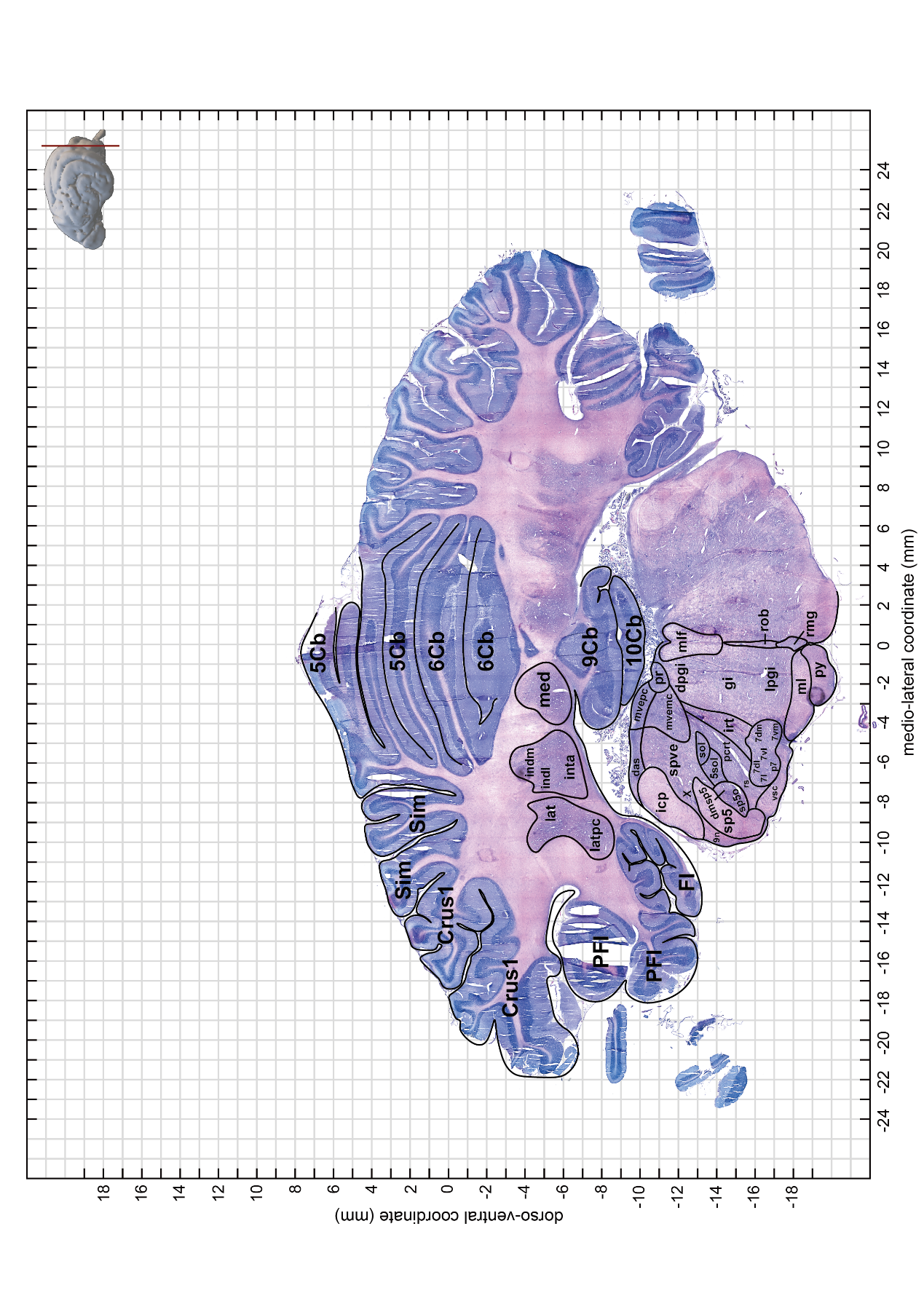

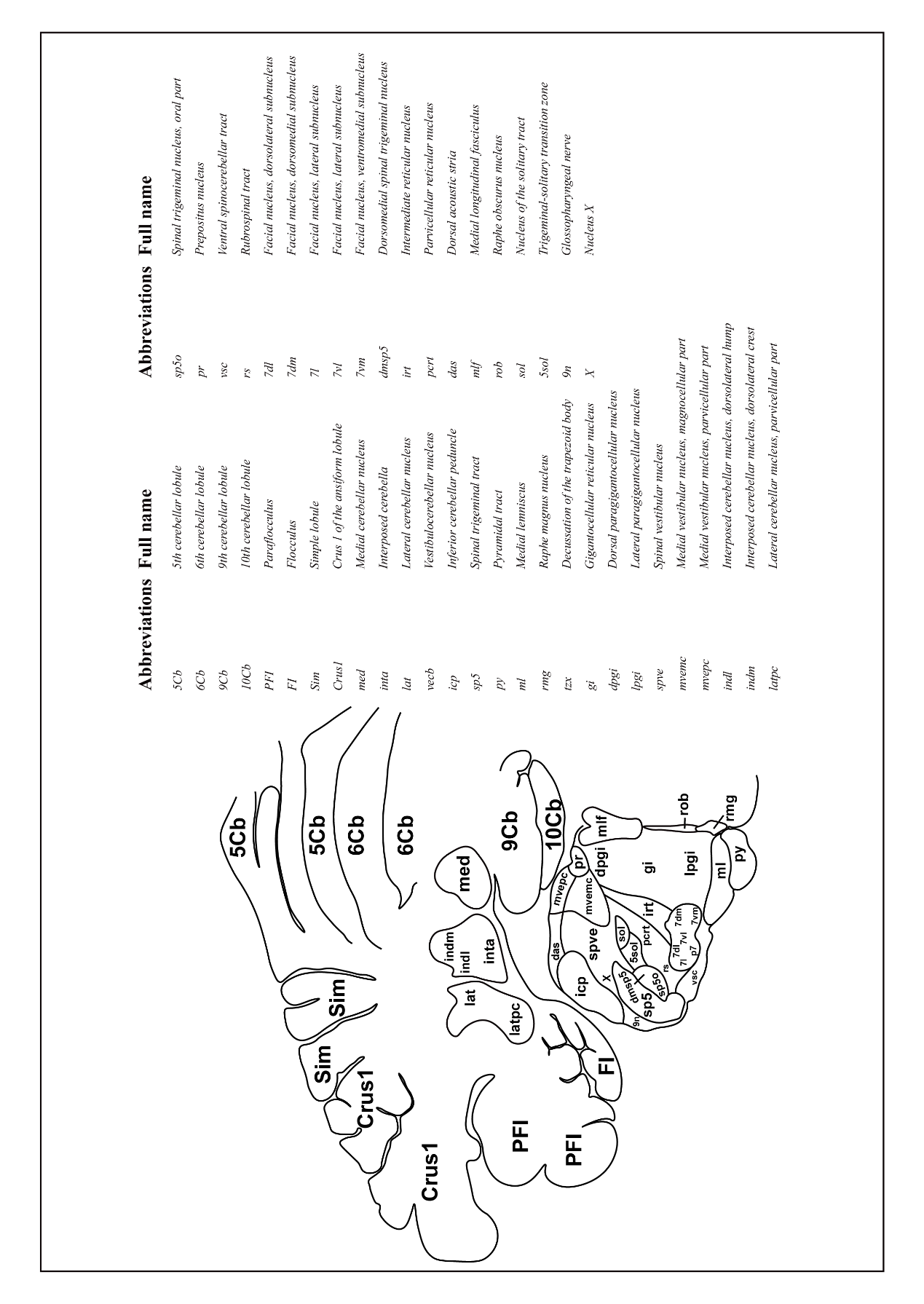

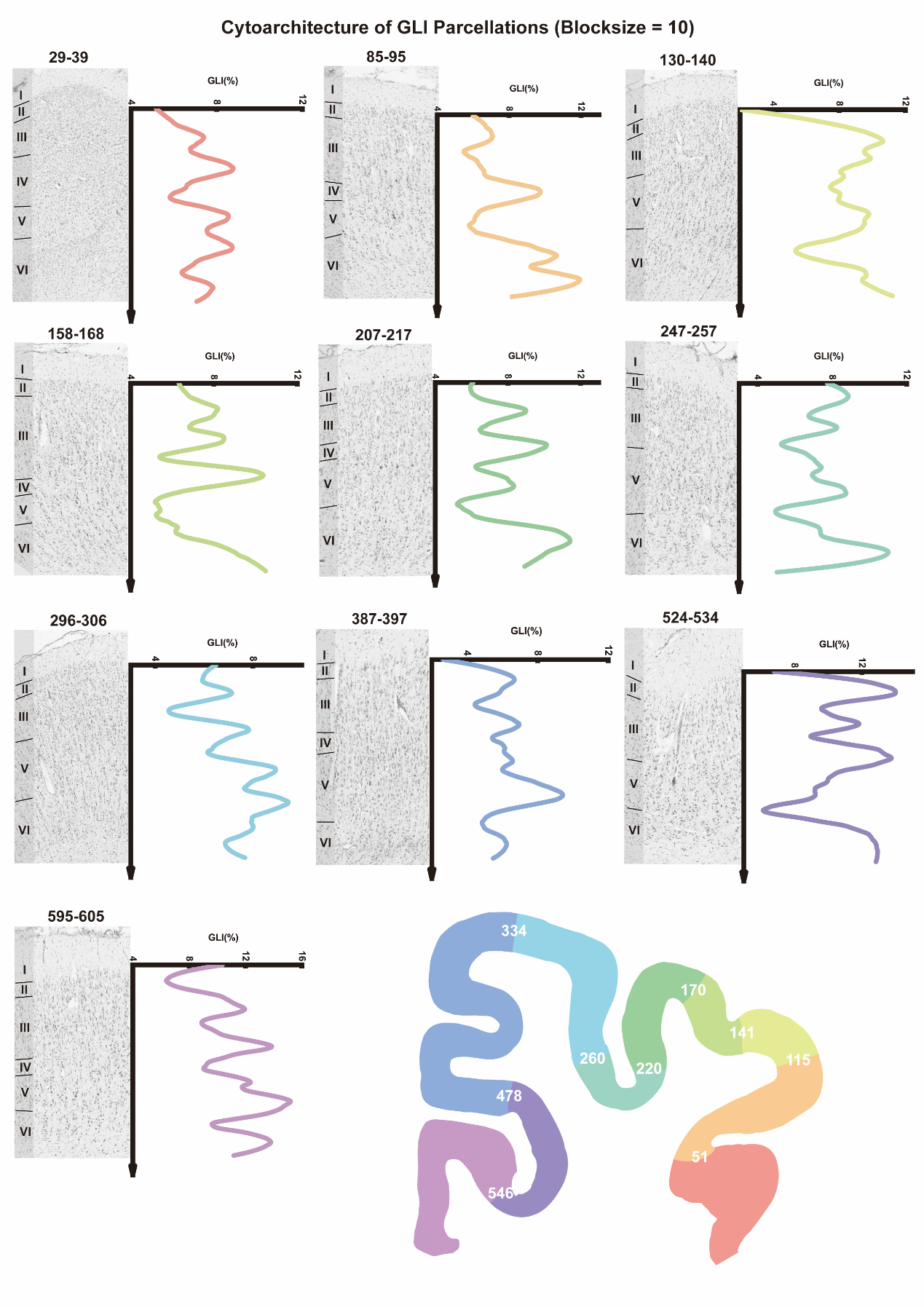
**Figure S17. The cytoarchitectonic and GLI curves of GLI parcellations in canine visual cortex (blocksize = 10).**

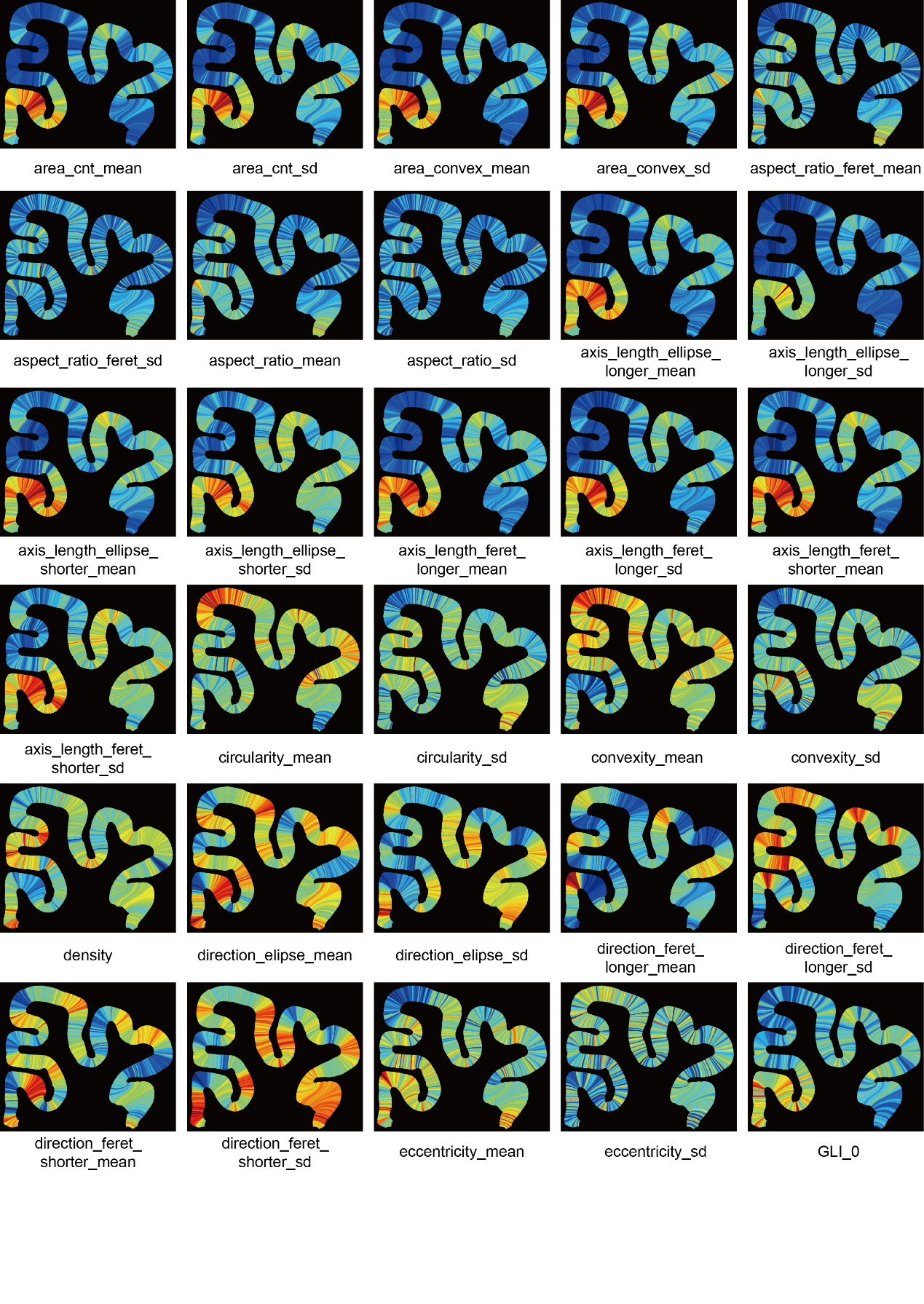

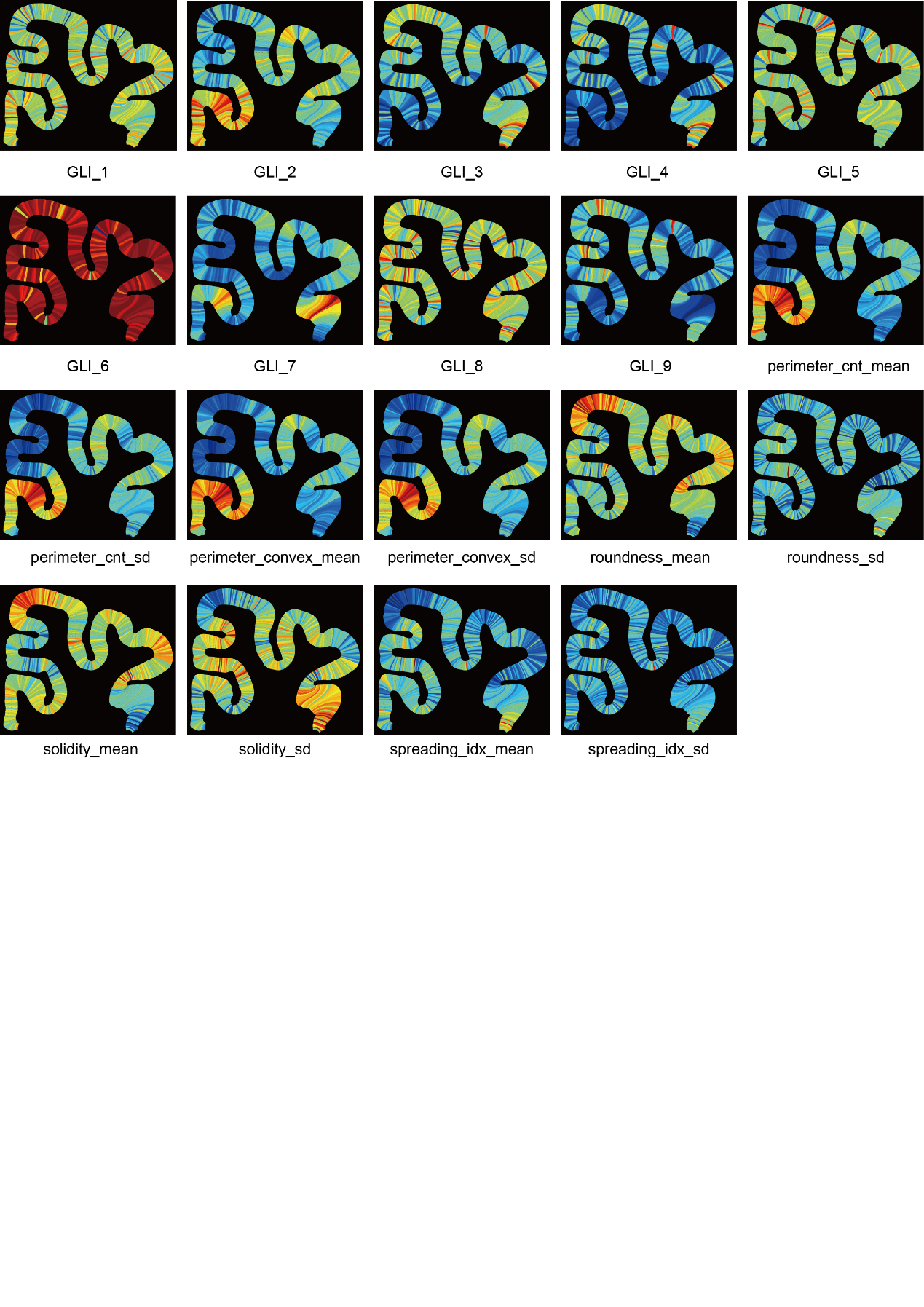

**Figure S18. The heatmap of the HCPM 49 cytoarchitectonic features in visual cortex of one slice example.**

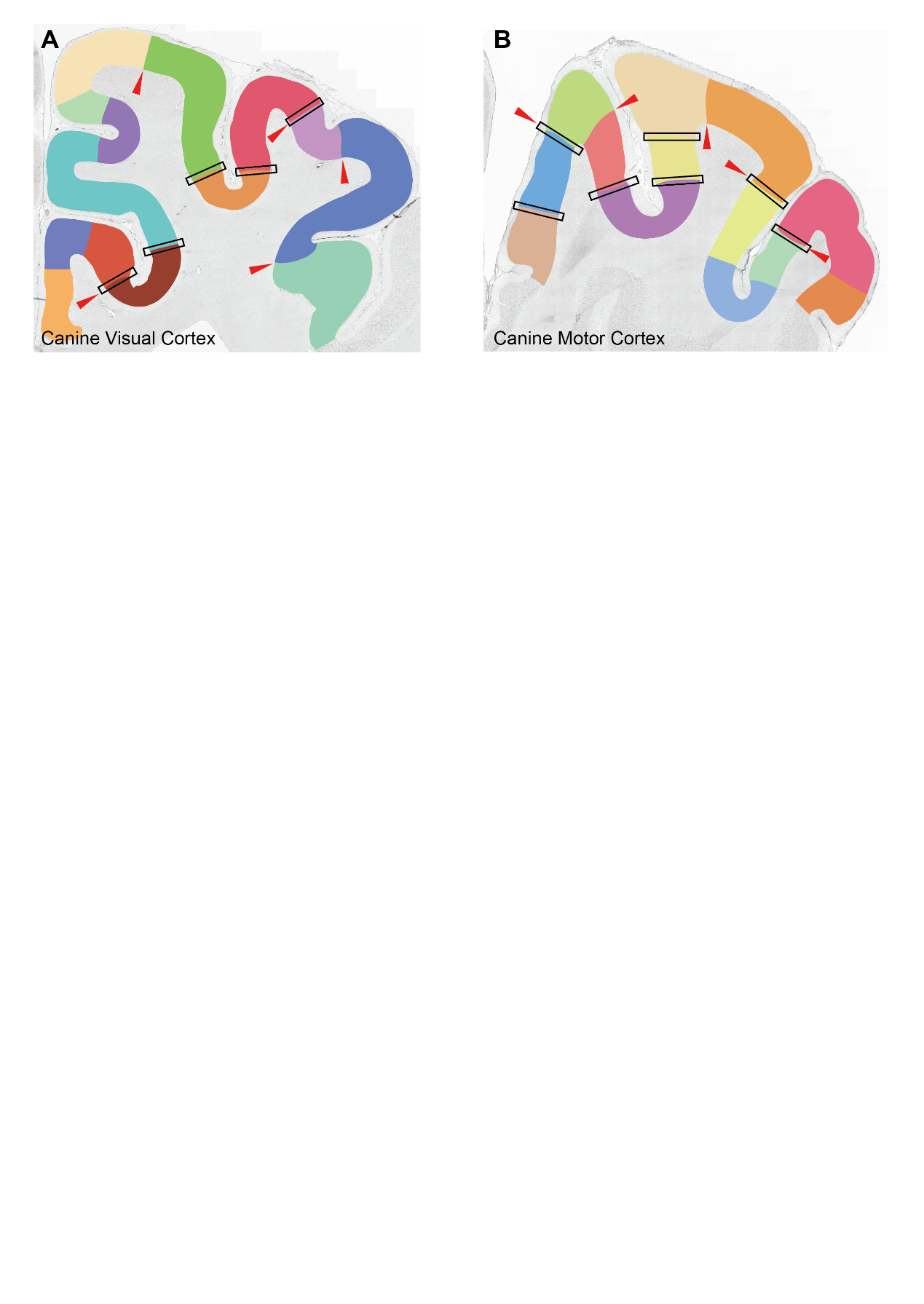

**Figure S19. Myeloarchitectonic parcellation of canine visual (A) and motor (B) cortex.** Cortical subregions are labeled with distinctive colors. Red arrowheads indicate consensus boundaries between myeloarchitecture and GLI, and black rectangles indicate consensus boundaries with HCPM.

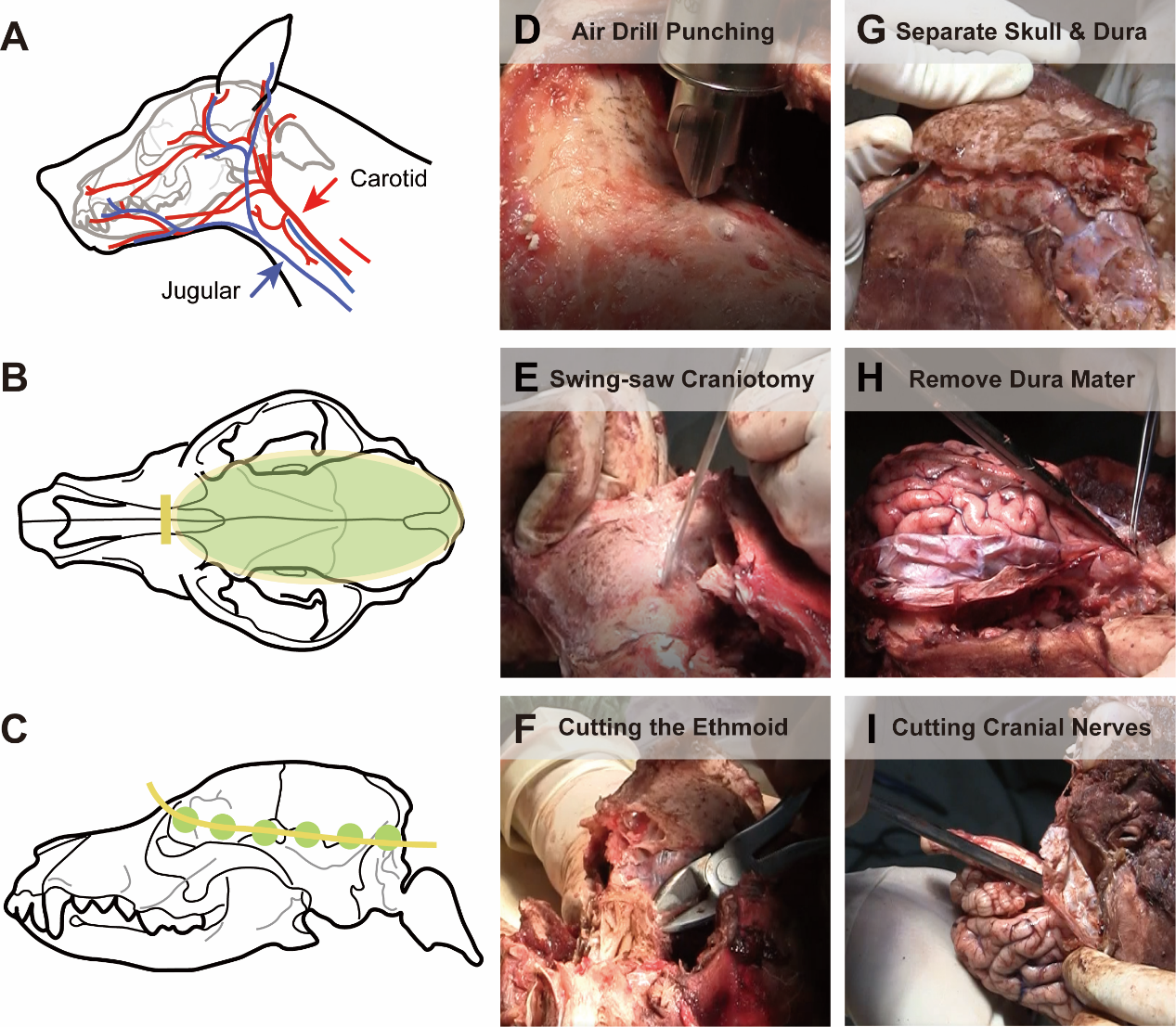

**Figure S20. Workflow of canine brain dissection.** (A) We choose to provide limited fixation of canine brains after euthanasia. We manually inject neutral formalin into head through the carotid artery and clip the jugular vein for bloodletting. (B) We use an air drill and a pendulum saw to circumcise the skull. Yellow line shows the cutting position and the green area represent the removed skull part. (C) The green dots are the positions for punching of the air drill and we connect these holes along the yellow line with the pendulum saw. (D)-(I) are photos of the anatomical steps. (D) is the air drill punching and (E) is the cutting of saw. (F) It is necessary to adequately open the ethmoid to create enough space for the dissection of the olfactory bulbs, because they are in a deep and narrow space and are easily broken. (G) Then we separate the skull and dura mater, and (H) we can see the brain tissue after opening the dura mater. (I) We can completely remove the brain tissue after the ventral nerves are cut off.

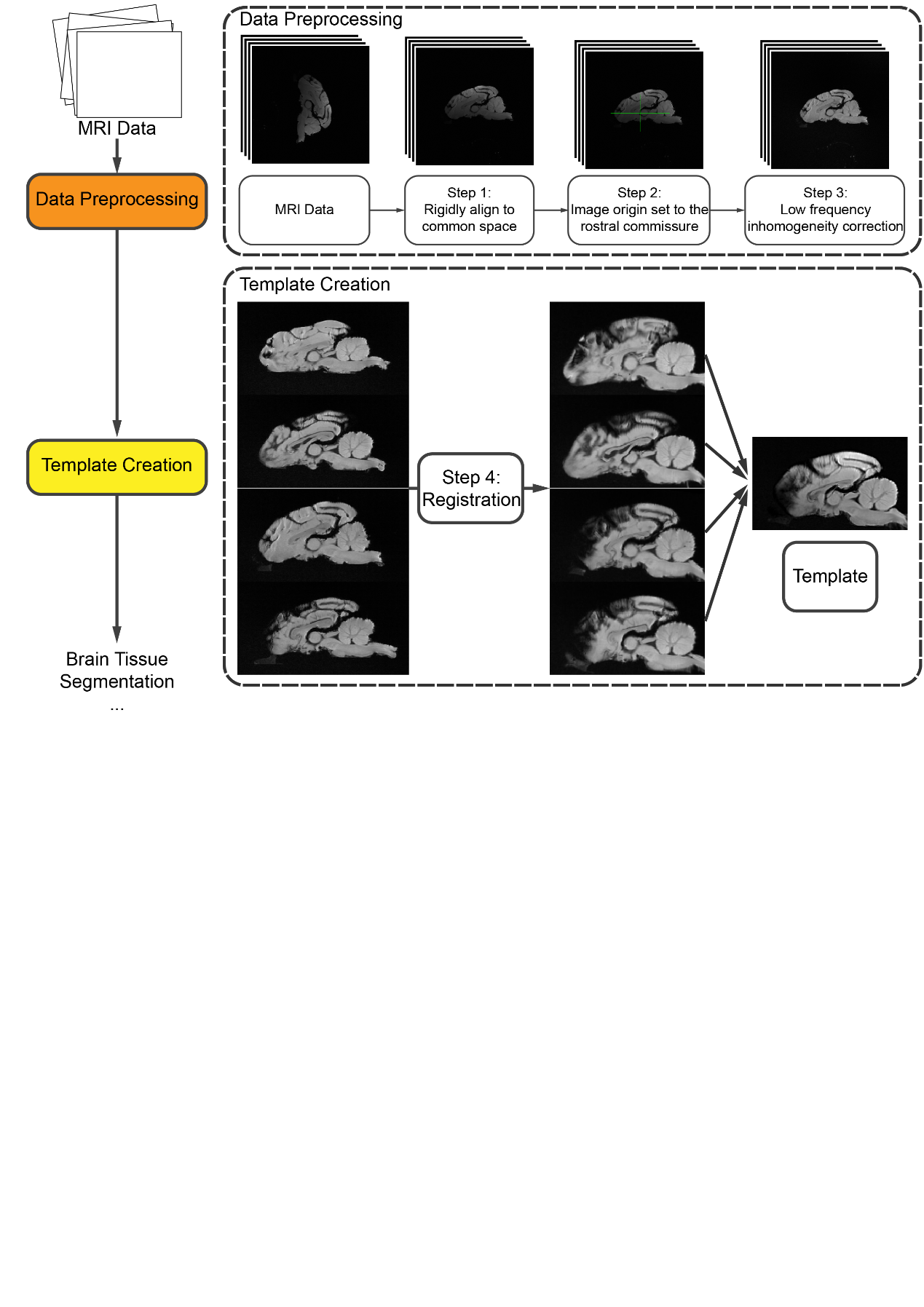

**Figure S21. Workflow of the brain MRI processing and template creation.** The images of the four subjects are first aligned into the MNI space (Step 1) and the origins are manually set to rostral commissure using SMP12 (Step 2). Then, N4 bias field correction is applied each image to reduce intensity non-uniformity using SimpleITK (Step 3). The MRI volumes are registered to a canine brain template before template construction (Step 4) exploiting Symmetric Normalization in ANTs. We use MultivariateTemplateConstruction command in ANTs to create the average template of Kunming dog.

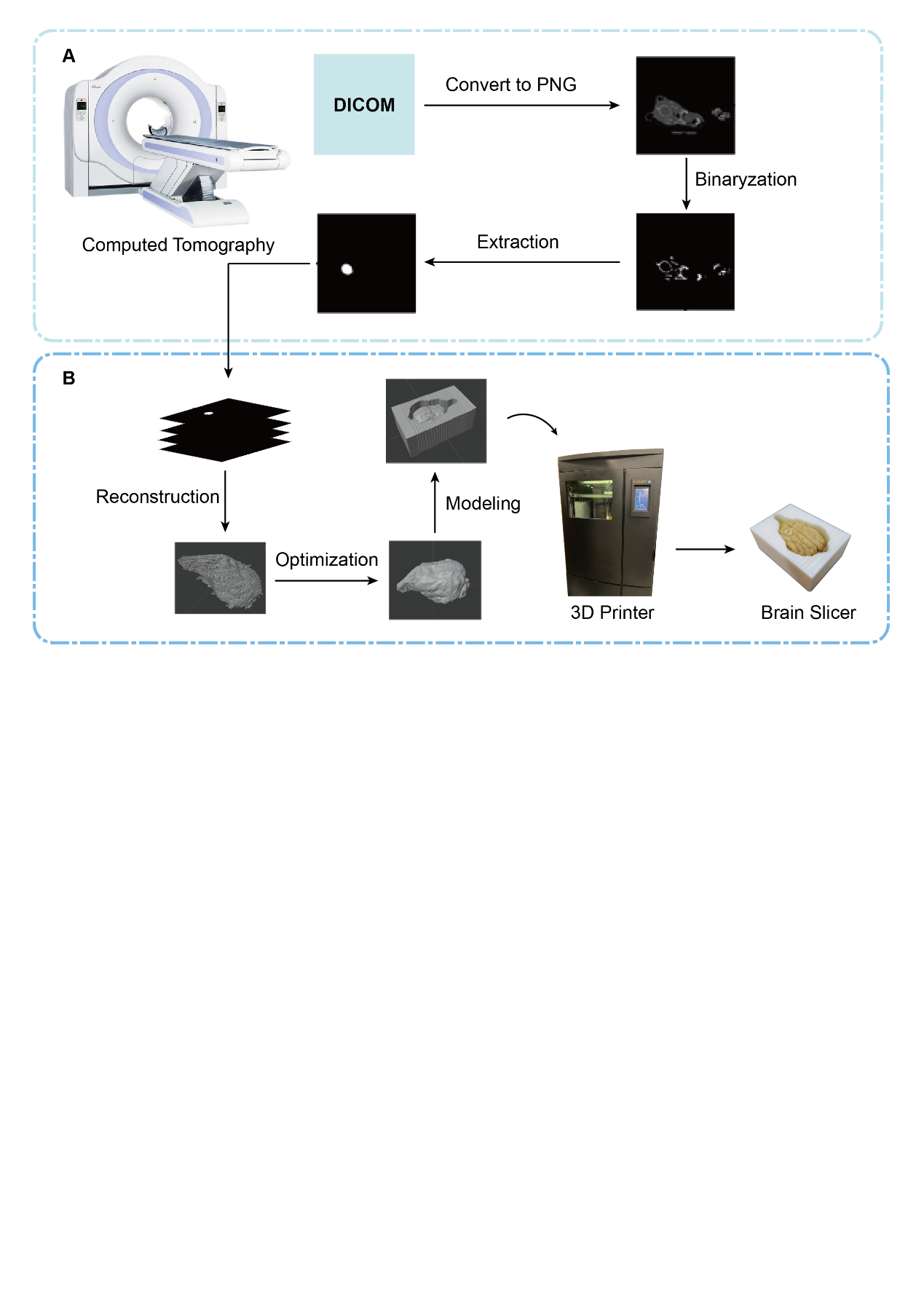

**Figure S22. Illustration of the construction and 3D printing a brain chunking mold for Kunming dog brain.** (A) shows the preprocessing of serial Computed Tomography (CT) images of the Kunming dog brain. We select the CT images that cover the whole brain and convert the DICOM format to PNG. We binary them and manually segment the brain from skull. (B) We reconstruct a 3D brain model and smoothing sharp points and construct a chunking model that can accommodate the entire dog brain with 22 knife grooves (1.5mm width and 5mm interval) in the Blender. The model is printed in polylactic acid by HORI Z600.

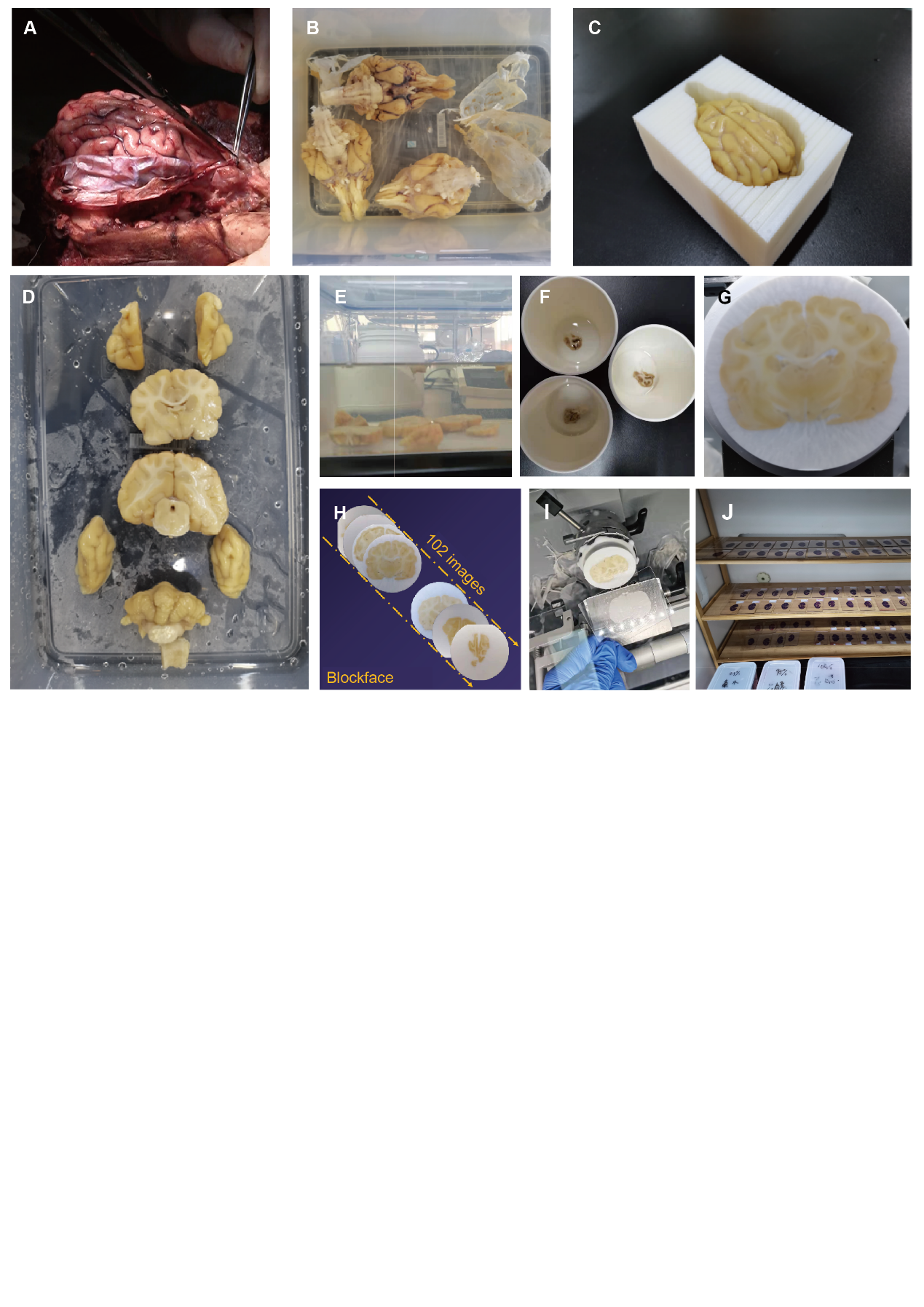

**Figure S23. Tissue processing, cryosection, and staining for brain samples.** (A) The dissection of brain from the skull. (B) Fix and store brains in formalin. (C) The isolated brain in the 3D printed brain chunking mold. (D) Coronal brain blocks. (E) Blocks are dehydrated in the sucrose solution with gradient concentrations. The samples are embedded with the OCT (F) in quick freezing (G). (H) A serial of blockface images. (I) Cryosections of brain blocks. (J) The stained sections are covered with neutral gum and coverslips and aired to dry.

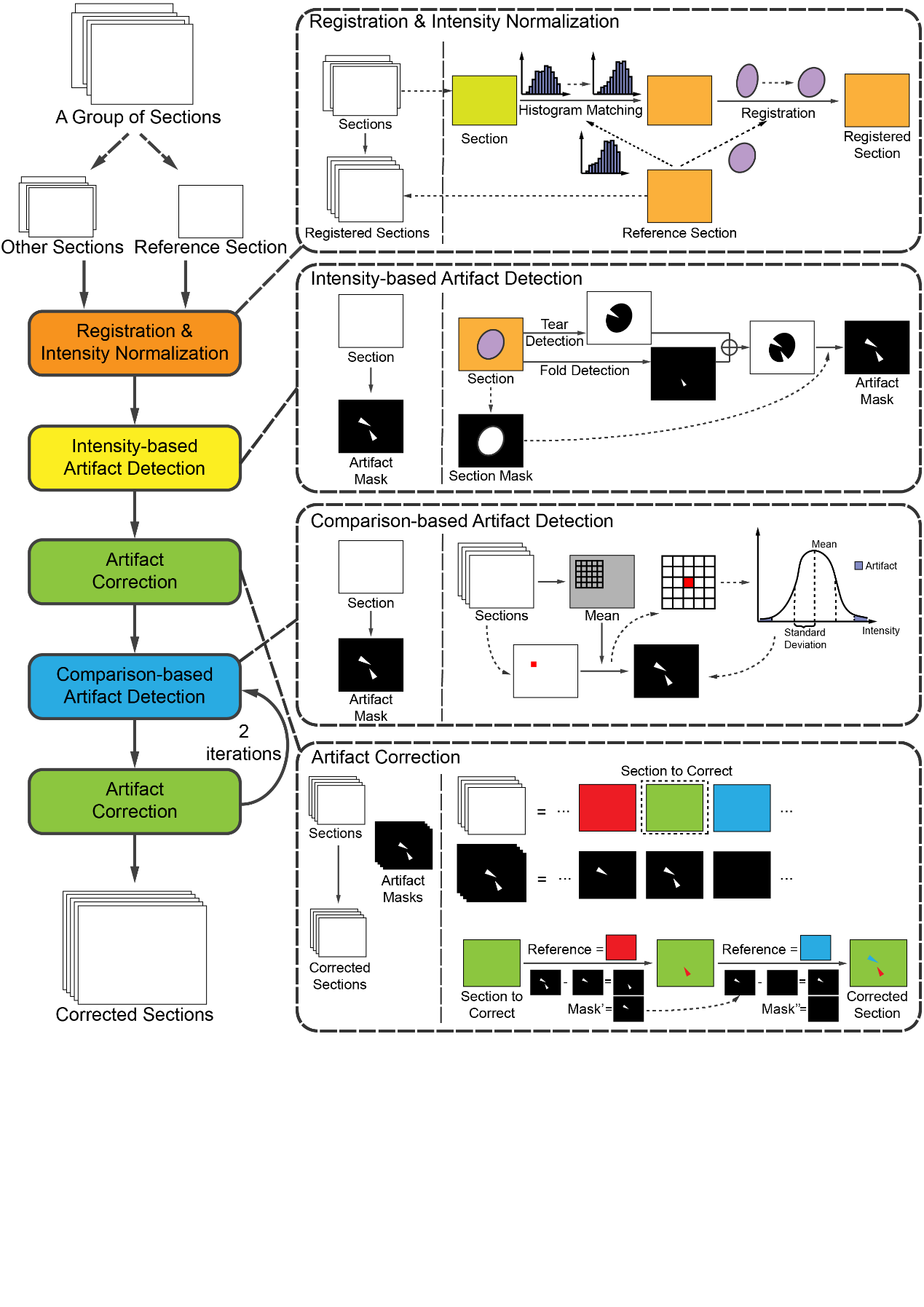

**Figure S24. The pipeline of the artifact detection and repairment of digital sections.**

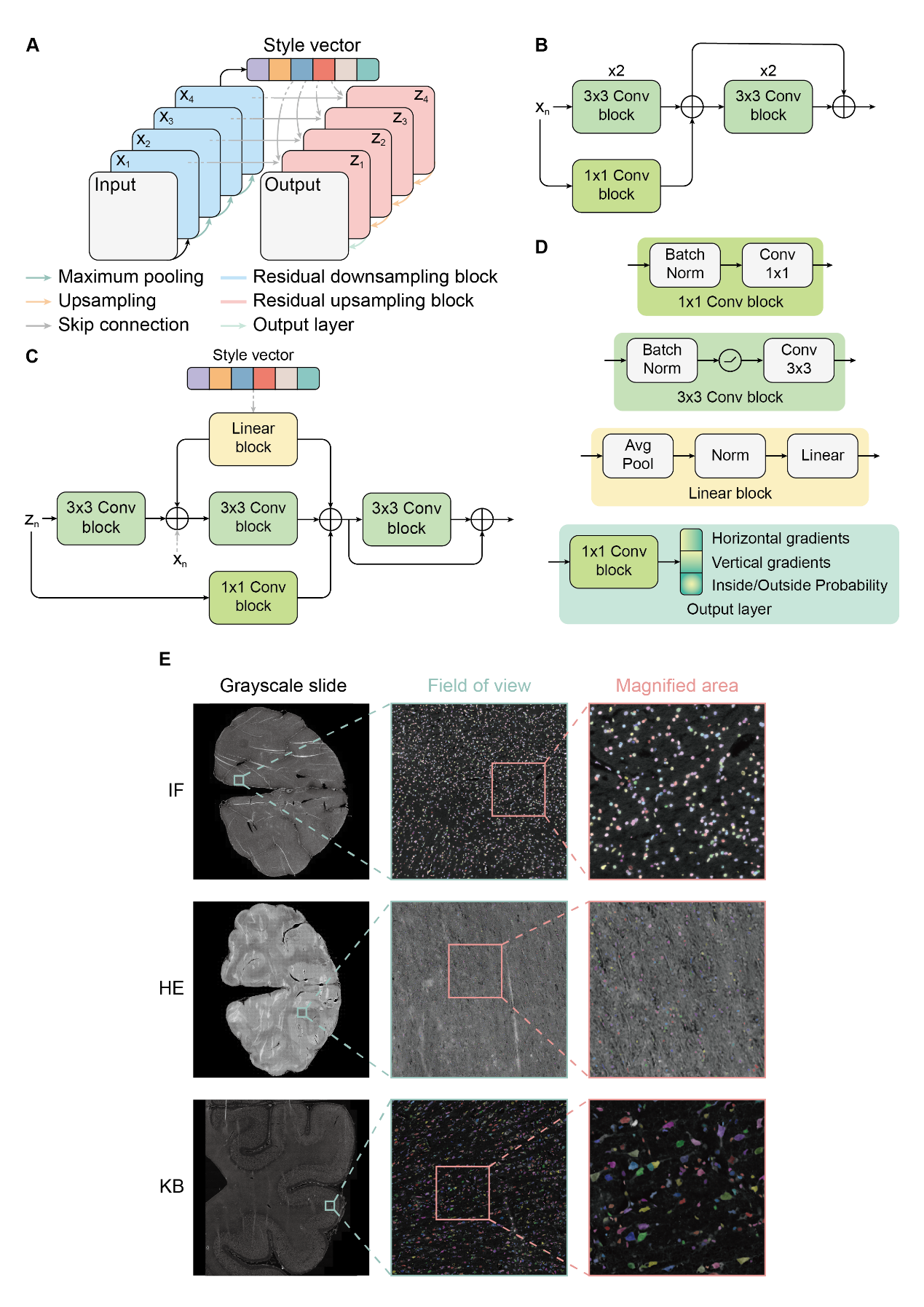

**Figure S25. The structure of the neural network of Cellpose.**
