## Supplementary Tables for "Hypergraph Cortical Cytoarchitectonic Parcellation with Multimodal Canine Brain Atlas"

*Corresponding Author*

**Table S1. Description of 49 cytoarchitectonic features of columnar unit in the HCPM**

| **Feature name** | **Description** | **Feature name** | **Description** |
| --- | --- | --- | --- |
| GLI_0 | Mean of column GLI | axis_length_feret_longer_sd | Standard deviation of longer axis of all instance Feret diameters |
| GLI_1 | Gravity center of column GLI | axis_length_feret_shorter_mean | Mean of shorter axis of all instance Feret diameters |
| GLI_2 | Standard deviation of column GLI | axis_length_feret_shorter_sd | Standard deviation of shorter axis of all instance Feret diameters |
| GLI_3 | Skewness of column GLI | circularity_mean | Mean of circularities of all instances (Value of 0 to 1 represents the degree from ellipse to perfect circle) |
| GLI_4 | Kurtosis of column GLI | circularity_sd | Standard deviation of circularities of all instances (Value of 0 to 1 represents the degree from ellipse to perfect circle) |
| GLI_5 | Mean of the first derivative of column GLI | convexity_mean | Mean of convexity of all instances (Lower values mean an enlargement of cell surface area) |
| GLI_6 | Gravity center of the profiles' first derivatives | convexity_sd | Standard deviation of convexity of all instances (Lower values mean an enlargement of cell surface area) |
| GLI_7 | Standard deviation of the first derivative of column GLI | direction_ellipse_mean | Mean of longer axis direction of all instance ellipse fitting |
| GLI_8 | Skewness of the first derivative of column GLI | direction_ellipse_sd | Standard deviation of longer axis direction of all instance ellipse fitting |
| GLI_9 | Kurtosis of the first derivative of column GLI | direction_feret_longer_mean | Mean of longer axis direction of all instance Feret diameters |
| area_cnt_mean | Mean of area of all instance masks | direction_feret_longer_sd | Standard deviation of longer axis direction of all instance Feret diameters |
| area_cnt_sd | Standard deviation of area of all instance masks | direction_feret_shorter_mean | Mean of shorter axis direction of all instance Feret diameters |
| area_convex_mean | Mean of area of all instance convex hulls | direction_feret_shorter_sd | Standard deviation of shorter axis direction of all instance Feret diameters |
| area_convex_sd | Standard deviation of area of all instance convex hulls | eccentricity_mean | Mean of eccentricity of all instance ellipse fitting |
| aspect_ratio_feret_mean | Mean of aspect ratio of all instance Feret diameters (The ratio of the maximum and minimum values of axis length) | eccentricity_sd | Standard deviation of eccentricity of all instance ellipse fitting |
| aspect_ratio_feret_sd | Standard deviation of aspect ratio of all instance Feret diameters (The ratio of the maximum and minimum values of axis length) | perimeter_cnt_mean | Mean of perimeter of all instance masks |
| aspect_ratio_mean | Mean of aspect ratio of all instance ellipse fitting | perimeter_cnt_sd | Standard deviation of perimeter of all instance masks |
| aspect_ratio_sd | Standard deviation of aspect ratio of all instance ellipse fitting | perimeter_convex_mean | Mean of perimeter of all instance convex hulls |
| spreading_idx_mean | Mean of spreading indexes of all instances (Larger values indicate more elongated structures) | perimeter_convex_sd | Standard deviation of perimeter of all instance convex hulls |
| spreading_idx_sd | Standard deviation of spreading indexes of all instances (Larger values indicate more elongated structures) | roundness_mean | Mean of roundness of all instances (Another circularity with Feret axis) |
| axis_length_ellipse_longer_mean | Mean of longer axis of all instance ellipse fitting | roundness_sd | Standard deviation of roundness of all instances (Another circularity with Feret axis) |
| axis_length_ellipse_longer_sd | Standard deviation of longer axis of all instance ellipse fitting | solidity_mean | Mean of solidity of all instances (The regular degree of boundaries) |
| axis_length_ellipse_shorter_mean | Mean of shorter axis of all instance ellipse fitting | solidity_sd | Standard deviation of solidity of all instances (The regular degree of boundaries) |
| axis_length_ellipse_shorter_sd | Standard deviation of shorter axis of all instance ellipse fitting | density | Instance density |
| axis_length_feret_longer_mean | Mean of longer axis of all instance Feret diameters |  |  |

**Table S2. The full names of abbreviations on the Figure 2A.**

| **Abbreviation** | **Full name** | **Abbreviation** | **Full name** |
| --- | --- | --- | --- |
| IIn | optic nerve | CbH | cerebellar hemisphere |
| Vn | trigeminal nerve | Cog | coronal gyrus |
| VIIIn | vestibulocochlear nerve | Cos | coronal sulcus |
| aESg | anterior ectosylvian gyrus | Cs | cruciate sulcus |
| aESs | anterior ectosylvian sulcus | Els | ectolateral sulcus |
| aRs | anterior rhinal sulcus | ESg | ectosylvian gyrus |
| aSg | anterior sigmoid gyrus | Emg | ectomarginal gyrus |
| aSSg | anterior suprasylvian gyrus | Fl | flocculus |
| aSSs | anterior suprasylvian sulcus | Is | intraproreus sulcus |
| aSyg | anterior sylvian gyrus | Lg | lateral gyrus |
| mESg | medial ectrosylvian gyrus | Lot | lateral olfactory tract |
| mESs | medial ectosylvian sulcus | Ls | lateral sulcus |
| Mo | medulla oblongata | OB | main olfactory bulb |
| Mot | medial olfactory tract | Ol | olive |
| mSSg | medial suprasylvian gyrus | OG | orbital gyrus |
| mSSs | medial suprasylvian sulcus | PC | cerebral peduncle |
| mSyg | medial sylvian gyrus | Pcg | posterior cruciate gyrus |
| pSSs | posterior suprasylvian sulcus | Pcs | postcruciate sulcus |
| pSyg | posterior sylvian gyrus | pESg | posterior ectosylvian gyrus |
| Py | pyramidal tract | pESs | posterior ectosylvian sulcus |
| Sys | sylvian sulcus | PG | pituitary gland |
| Rg | rectus gyrus | Pir | piriform lobe |
| SSs | suprasylvian sulcus | pLg | posterior lateral gyrus |
| Sys | sylvian sulcus | pRs | posterior rhinal sulcus |
| To | olfactory tubercle | pSg | posterior sigmoid gyrus |
| Tz | trapezoid body | PSs | presylvian sulcus |
| V | vermis | pSSg | posterior syprasylvian gyrus |
| vMf | ventral median fissure | Pg | proreus gyrus |

**Table S3. MRI parcellations and statistics based on the myeloarchitecture.**

| Label Id | Label Name | Number of Voxels | Volume (mm^3) | Image mean (n4_template) | Image stdev (n4_template) |
| --- | --- | --- | --- | --- | --- |
| 0 | Clear Label | 2447390 | 268905 | 0.395181 | 0.804687 |
| 1 | Cerebellum | 71300 | 7834.03 | 3.20198 | 0.706108 |
| 2 | Spinal Cord | 5369 | 589.914 | 3.03844 | 0.775523 |
| 3 | Medulla Oblongata | 21240 | 2333.73 | 3.22211 | 0.505477 |
| 4 | Pons | 16044 | 1762.82 | 3.15284 | 0.487177 |
| 5 | Midbrain | 18493 | 2031.9 | 3.30444 | 0.332993 |
| 6 | Diencephalon | 33957 | 3731 | 3.26125 | 0.344369 |
| 7 | Optic Chiasm | 648 | 71.1984 | 2.24561 | 0.540502 |
| 8 | Olfactory Bulb | 15979 | 1755.68 | 1.81231 | 0.615163 |
| 10 | Corpus Callosum | 4334 | 476.195 | 3.12874 | 0.492724 |
| 11 | Hippocampus | 13249 | 1455.72 | 3.35549 | 0.47086 |
| 12 | Caudate Nucleus | 10771 | 1183.45 | 3.61732 | 0.268526 |
| 13 | Area Polaris R | 723 | 79.439 | 2.54691 | 0.621477 |
| 14 | Area Polaris L | 705 | 77.4613 | 2.56018 | 0.717434 |
| 15 | Area Subprorealis I R | 1709 | 187.775 | 2.98689 | 0.566547 |
| 16 | Area Subprorealis I L | 1628 | 178.875 | 3.01949 | 0.538371 |
| 17 | Area Prorealis Lateralis I R | 2294 | 252.051 | 3.12617 | 0.786831 |
| 18 | Area Prorealis Lateralis I L | 2168 | 238.207 | 3.1731 | 0.751362 |
| 19 | Area Orbitalis I L | 2066 | 227 | 3.07326 | 0.679115 |
| 20 | Area Orbitalis I R | 2038 | 223.924 | 3.09169 | 0.627158 |
| 21 | Area Fissurae Orbitalis L | 797 | 87.5697 | 3.43433 | 0.305744 |
| 22 | Area Fissurae Orbitalis R | 946 | 103.941 | 3.37128 | 0.332878 |
| 23 | Area Pregenualis II L | 1024 | 112.511 | 3.09607 | 0.71498 |
| 24 | Area Pregenualis II R | 932 | 102.403 | 3.02854 | 0.76033 |
| 25 | Area Marginalis Posterior R | 6991 | 768.13 | 3.82055 | 0.970458 |
| 26 | Area Marginalis Posterior L | 6469 | 710.776 | 3.87081 | 0.991999 |
| 27 | Area Retrospenialis R | 4280 | 470.261 | 3.48477 | 0.528709 |
| 28 | Area Retrospenialis L | 4037 | 443.562 | 3.50411 | 0.487764 |
| 29 | Area Entolateralis Posterior R | 4849 | 532.78 | 3.81833 | 0.720899 |
| 30 | Area Entolateralis Posterior L | 4789 | 526.187 | 3.8421 | 0.710623 |
| 31 | Area Fissurae Lateralis R | 2151 | 236.339 | 3.41513 | 0.266485 |
| 32 | Area Fissurae Lateralis L | 2291 | 251.722 | 3.44203 | 0.344285 |
| 33 | Area Ectolateralis Posterior R | 6240 | 685.615 | 4.05414 | 1.04267 |
| 34 | Area Ectolateralis Posterior L | 5685 | 624.635 | 4.1474 | 1.01698 |
| 35 | Area Suprasylvian Medialis R | 6259 | 687.702 | 3.7147 | 0.75077 |
| 36 | Area Suprasylvian Medialis L | 6821 | 749.452 | 3.66971 | 0.756192 |
| 37 | Area Fissurea Ectolateralis R | 773 | 84.9327 | 3.38618 | 0.243989 |
| 38 | Area Fissurea Ectolateralis L | 910 | 99.9855 | 3.38735 | 0.264151 |
| 39 | Area Suprasylvian Posterior R | 3633 | 399.173 | 3.51705 | 0.544268 |
| 40 | Area Suprasylvian Posterior L | 3000 | 329.622 | 3.4848 | 0.583548 |
| 41 | Area Pararecurrens Medialis R | 297 | 32.6326 | 3.4702 | 0.26601 |
| 42 | Area Pararecurrens Medialis L | 256 | 28.1278 | 3.54587 | 0.340538 |
| 43 | Area Pararecurrens Lateralis R | 573 | 62.9579 | 3.45135 | 0.728253 |
| 44 | Area Pararecurrens Lateralis L | 599 | 65.8146 | 3.80606 | 0.727657 |
| 45 | Area Recurrens Ventralis Lateralis R | 405 | 44.499 | 3.80094 | 0.804761 |
| 46 | Area Recurrens Ventralis Medialis L | 436 | 47.9051 | 3.72979 | 0.58445 |
| 47 | Area Splenialis I R | 3925 | 431.256 | 3.23827 | 0.334162 |
| 48 | Area Splenialis I L | 3497 | 384.23 | 3.22157 | 0.323856 |
| 49 | Area Fissurae Suprasplenialis R | 746 | 81.9661 | 3.20427 | 0.315241 |
| 50 | Area Fissurae Suprasplenialis L | 912 | 100.205 | 3.26227 | 0.273553 |
| 51 | Area Splenialis II R | 746 | 81.9661 | 3.46181 | 0.43467 |
| 52 | Area Splenialis II L | 722 | 79.3291 | 3.49123 | 0.433338 |
| 53 | Area Fissurea Splenialis R | 2348 | 257.985 | 3.4766 | 0.220404 |
| 54 | Area Fissurea Splenialis L | 2595 | 285.123 | 3.45328 | 0.209934 |
| 55 | Area Limbica Posterior Lateralis R | 588 | 64.606 | 3.38606 | 0.179328 |
| 56 | Area Limbica Posterior Lateralis L | 530 | 58.2333 | 3.38645 | 0.145979 |
| 57 | Area Limbica Posterior Dorsalis I R | 985 | 108.226 | 3.35172 | 0.209364 |
| 58 | Area Limbica Posterior Dorsalis I L | 1116 | 122.62 | 3.35654 | 0.244303 |
| 59 | Area Limbica Posterior Ventralis I R | 530 | 58.2333 | 3.48621 | 0.253528 |
| 60 | Area Limbica Posterior Ventralis I L | 570 | 62.6283 | 3.46549 | 0.248705 |
| 61 | Area Fissurae Calloso-Marginalis R | 270 | 29.666 | 3.24068 | 0.164396 |
| 62 | Area Fissurae Calloso-Marginalis L | 265 | 29.1166 | 3.29017 | 0.142356 |
| 63 | Area Fissurea Suprasylviae Anterior R | 1676 | 184.149 | 3.54753 | 0.293444 |
| 64 | Area Fissurea Suprasylviae Anterior L | 1657 | 182.061 | 3.53547 | 0.262879 |
| 65 | Area Paraectosylvia Dorsalis II R | 3888 | 427.191 | 3.37614 | 0.268018 |
| 66 | Area Paraectosylvia Dorsalis II L | 4331 | 475.865 | 3.38554 | 0.272477 |
| 67 | Area Ectosylvia Medialis R | 6189 | 680.011 | 3.40799 | 0.69458 |
| 68 | Area Ectosylvia Medialis L | 5341 | 586.838 | 3.50108 | 0.744966 |
| 69 | Area Paraectosylvia Ventralis R | 2202 | 241.943 | 3.44809 | 0.262225 |
| 70 | Area Paraectosylvia Ventralis L | 2148 | 236.01 | 3.50455 | 0.239847 |
| 71 | Area Fissurae Ectosylvia R | 3341 | 367.09 | 3.48337 | 0.26095 |
| 72 | Area Fissurae Ectosylvia L | 3445 | 378.516 | 3.53754 | 0.250439 |
| 73 | Area Sylvia R | 7689 | 844.822 | 3.18558 | 0.563367 |
| 74 | Area Sylvia L | 6624 | 727.806 | 3.2313 | 0.495816 |
| 75 | Area Parasylvian Dorsalis R | 769 | 84.4932 | 3.46688 | 0.217865 |
| 76 | Area Parasylvian Dorsalis L | 834 | 91.635 | 3.4972 | 0.196488 |
| 77 | Area Ectosylvia Posterior I R | 3103 | 340.939 | 3.17027 | 0.541044 |
| 78 | Area Ectosylvia Posterior I L | 3617 | 397.415 | 3.20757 | 0.492372 |
| 79 | Area Composita Posterior Lateralis I R | 2157 | 236.999 | 3.23205 | 0.470091 |
| 80 | Area Composita Posterior Lateralis I L | 2412 | 265.016 | 3.24383 | 0.443774 |
| 81 | Area Composita Medialis I R | 4167 | 457.846 | 3.00052 | 0.606691 |
| 82 | Area Composita Medialis I L | 4682 | 514.431 | 3.1234 | 0.613328 |
| 83 | Area Fissurea Recurrentis R | 1306 | 143.496 | 3.37771 | 0.214232 |
| 84 | Area Fissurea Recurrentis L | 1452 | 159.537 | 3.35828 | 0.204021 |
| 85 | Area Limbica Posterior Dorsalis II R | 1023 | 112.401 | 3.23771 | 0.342908 |
| 86 | Area Limbica Posterior Dorsalis II L | 915 | 100.535 | 3.20929 | 0.319921 |
| 87 | Area Fissurae Sylvia R | 2654 | 291.606 | 3.39931 | 0.340193 |
| 88 | Area Fissurae Sylvia L | 2703 | 296.99 | 3.49245 | 0.301463 |
| 89 | Claustrum R | 842 | 92.514 | 3.39982 | 0.157942 |
| 90 | Claustrum L | 801 | 88.0092 | 3.49673 | 0.155675 |
| 91 | Area Ectosylvia Accessoria R | 2460 | 270.29 | 3.27829 | 0.671156 |
| 92 | Area Ectosylvia Accessoria L | 2304 | 253.15 | 3.33919 | 0.60186 |
| 93 | Area Suprasylvian Accessoria R | 115 | 12.6355 | 3.56962 | 0.172507 |
| 94 | Area Suprasylvian Accessoria L | 102 | 11.2072 | 3.59792 | 0.179141 |
| 95 | Area Coronalis Posterior R | 1928 | 211.837 | 3.29251 | 0.5939 |
| 96 | Area Coronalis Posterior L | 1370 | 150.528 | 3.21417 | 0.63047 |
| 97 | Area Limbica Media R | 638 | 70.0997 | 3.48055 | 0.184574 |
| 98 | Area Limbica Media L | 729 | 80.0983 | 3.43356 | 0.204193 |
| 99 | Area Presplenialis Dorsalis R | 1628 | 178.875 | 3.32409 | 0.288864 |
| 100 | Area Presplenialis Dorsalis L | 1701 | 186.896 | 3.19722 | 0.444494 |
| 101 | Area Marginalis Anterior R | 1489 | 163.603 | 3.39253 | 0.697612 |
| 102 | Area Marginalis Anterior L | 1650 | 181.292 | 3.33972 | 0.835383 |
| 103 | Area Limbica Anterior Ventralis R | 656 | 72.0774 | 3.36196 | 0.21275 |
| 104 | Area Limbica Anterior Ventralis L | 653 | 71.7478 | 3.37518 | 0.218798 |
| 105 | Area Limbica Anterior Dorsalis I R | 434 | 47.6854 | 3.42195 | 0.343524 |
| 106 | Area Limbica Anterior Dorsalis I L | 380 | 41.7522 | 3.36147 | 0.36126 |
| 107 | Area Limbica Anterior Lateralis R | 413 | 45.378 | 3.58617 | 0.231036 |
| 108 | Area Limbica Anterior Lateralis L | 471 | 51.7507 | 3.5153 | 0.228069 |
| 109 | Area Fissurae Splenialis R | 933 | 102.513 | 3.60747 | 0.275446 |
| 110 | Area Fissurae Splenialis L | 1130 | 124.158 | 3.57142 | 0.270633 |
| 111 | Area Precentralis Interna R | 1345 | 147.781 | 3.50759 | 0.263673 |
| 112 | Area Precentralis Interna L | 1080 | 118.664 | 3.51542 | 0.316439 |
| 113 | Area Precentralis III R | 1231 | 135.255 | 3.27118 | 0.760218 |
| 114 | Area Precentralis III L | 1197 | 131.519 | 3.34168 | 0.959078 |
| 115 | Area Centralis R | 464 | 50.9816 | 3.35042 | 0.453301 |
| 116 | Area Centralis L | 399 | 43.8398 | 3.41804 | 0.501314 |
| 117 | Area Postcentralis I R | 1926 | 211.618 | 3.34993 | 0.580808 |
| 118 | Area Postcentralis I L | 1731 | 190.192 | 3.36641 | 0.637355 |
| 119 | Area Precentralis I/II R | 1617 | 177.666 | 3.35959 | 0.57334 |
| 120 | Area Precentralis I/II L | 1568 | 172.283 | 3.37921 | 0.617584 |
| 121 | Area Precentral Lateralis R | 317 | 34.8301 | 3.54574 | 0.20149 |
| 122 | Area Precentral Lateralis L | 262 | 28.787 | 3.57028 | 0.279766 |
| 123 | Area Fissurae Coronalis R | 1705 | 187.335 | 3.50077 | 0.315896 |
| 124 | Area Fissurae Coronalis L | 1469 | 161.405 | 3.52182 | 0.330044 |
| 125 | Area Coronalis Medialis R | 632 | 69.4405 | 3.45825 | 0.254963 |
| 126 | Area Coronalis Medialis L | 718 | 78.8896 | 3.40557 | 0.319338 |
| 127 | Area Coronalis Anterior R | 4281 | 470.371 | 3.22863 | 0.654483 |
| 128 | Area Coronalis Anterior L | 4133 | 454.11 | 3.24789 | 0.680629 |
| 129 | Area Composita Ectosylvia R | 2272 | 249.634 | 3.26948 | 0.44341 |
| 130 | Area Composita Ectosylvia L | 2239 | 246.008 | 3.20897 | 0.593119 |
| 131 | Area Sylvia Insularis R | 1422 | 156.241 | 3.15256 | 0.432113 |
| 132 | Area Sylvia Insularis L | 1545 | 169.756 | 3.11688 | 0.431398 |
| 133 | Area Fissurae Presylviae R | 1590 | 174.7 | 3.53046 | 0.283725 |
| 134 | Area Fissurae Presylviae L | 1648 | 181.073 | 3.56354 | 0.238255 |
| 135 | Area Composita Interna R | 2139 | 235.021 | 3.34269 | 0.337319 |
| 136 | Area Composita Interna L | 2187 | 240.295 | 3.36476 | 0.325206 |
| 137 | Area Orbitalis II R | 3262 | 358.409 | 3.37349 | 0.279289 |
| 138 | Area Orbitalis II L | 3545 | 389.504 | 3.36274 | 0.282837 |
| 139 | Area Paraorbitalis Ventralis R | 463 | 50.8717 | 3.3195 | 0.254988 |
| 140 | Area Paraorbitalis Ventralis L | 433 | 47.5755 | 3.39735 | 0.275837 |
| 141 | Area Subprorealis II R | 258 | 28.3475 | 3.60162 | 0.14661 |
| 142 | Area Subprorealis II L | 264 | 29.0068 | 3.59754 | 0.19914 |
| 143 | Area Prorealis R | 547 | 60.1012 | 3.61995 | 0.455958 |
| 144 | Area Prorealis L | 546 | 59.9913 | 3.6374 | 0.424693 |
| 145 | Area Subgenualis R | 845 | 92.8437 | 3.63633 | 0.190681 |
| 146 | Area Subgenualis L | 857 | 94.1621 | 3.61978 | 0.208878 |
| 147 | Area Precruciata Medialis I R | 1126 | 123.718 | 3.39947 | 0.23242 |
| 148 | Area Precruciata Medialis I L | 1156 | 127.015 | 3.36465 | 0.271695 |
| 149 | Area Composita Sigmoid R | 1086 | 119.323 | 3.36965 | 0.372915 |
| 150 | Area Composita Sigmoid L | 828 | 90.9758 | 3.52805 | 0.476195 |
| 151 | Area Composita Precruciata R | 1362 | 149.649 | 3.34474 | 0.505658 |
| 152 | Area Composita Precruciata L | 1764 | 193.818 | 3.29601 | 0.524715 |
| 153 | Area Composita Anterior R | 522 | 57.3543 | 2.93798 | 0.551001 |
| 154 | Area Composita Anterior L | 467 | 51.3112 | 2.97926 | 0.603298 |
| 155 | Area Fissurae Pregenualis R | 726 | 79.7686 | 3.4485 | 0.373064 |
| 156 | Area Fissurae Pregenualis L | 687 | 75.4835 | 3.39799 | 0.41872 |
| 157 | Area Precruciata Medialis II R | 991 | 108.885 | 3.4732 | 0.31833 |
| 158 | Area Precruciata Medialis II L | 942 | 103.501 | 3.41015 | 0.296959 |
| 159 | Area Pregenualis III R | 157 | 17.2502 | 3.4588 | 0.466711 |
| 160 | Area Pregenualis III L | 117 | 12.8553 | 3.48254 | 0.490514 |
| 161 | Area Precruciata Centralis R | 683 | 75.044 | 3.3621 | 0.447839 |
| 162 | Area Precruciata Centralis L | 739 | 81.197 | 3.24817 | 0.593571 |
| 163 | Area Precruciata Posterior R | 743 | 81.6365 | 3.28095 | 0.399103 |
| 164 | Area Precruciata Posterior L | 725 | 79.6588 | 3.34003 | 0.267872 |
| 165 | Area Genualis I R | 915 | 100.535 | 3.4784 | 0.194317 |
| 166 | Area Genualis I L | 802 | 88.1191 | 3.44713 | 0.215558 |
| 167 | Area Subprorealis II R | 826 | 90.756 | 3.08183 | 0.383018 |
| 168 | Area Subprorealis II L | 868 | 95.3708 | 3.0784 | 0.37085 |
| 171 | Area Limbica Anterior Dorsalis II R | 317 | 34.8301 | 3.58622 | 0.246814 |
| 172 | Area Limbica Anterior Dorsalis II L | 341 | 37.4671 | 3.52956 | 0.330956 |
| 173 | Area Fissurae Ansata R | 265 | 29.1166 | 3.30295 | 0.207709 |
| 174 | Area Fissurae Ansata L | 339 | 37.2473 | 3.31355 | 0.244914 |
| 175 | Area Presplenialis Ventralis R | 440 | 48.3446 | 3.56971 | 0.169495 |
| 176 | Area Presplenialis Ventralis L | 518 | 56.9148 | 3.5171 | 0.169688 |
| 177 | Area Fissurae Presylviae Lateralis R | 555 | 60.9802 | 3.52624 | 0.178644 |
| 178 | Area Fissurae Presylviae Lateralis L | 614 | 67.4627 | 3.47035 | 0.167707 |
| 179 | Area Ectolateralis Anterior R | 731 | 80.318 | 3.35972 | 0.528025 |
| 180 | Area Ectolateralis Anterior L | 650 | 71.4182 | 3.60456 | 0.517952 |
| 181 | Area Marginalis Lateralis R | 149 | 16.3712 | 3.20736 | 0.19876 |
| 182 | Area Marginalis Lateralis L | 178 | 19.5576 | 3.22078 | 0.367576 |
| 183 | Area Fissurae Suprasplenialis R | 182 | 19.9971 | 3.04142 | 0.421192 |
| 184 | Area Fissurae Suprasplenialis L | 203 | 22.3045 | 3.10241 | 0.322042 |
| 185 | Area Entolateralis Anterior Lateralis R | 252 | 27.6883 | 3.20543 | 0.261986 |
| 186 | Area Entolateralis Anterior Lateralis L | 306 | 33.6215 | 3.37695 | 0.17547 |
| 187 | Area Fissurae Lateralis R | 291 | 31.9734 | 3.46702 | 0.207984 |
| 188 | Area Fissurae Lateralis L | 313 | 34.3906 | 3.50777 | 0.202685 |
| 189 | Area Coronalis Posterior Medialis R | 721 | 79.2193 | 3.41071 | 0.214315 |
| 190 | Area Coronalis Posterior Medialis L | 547 | 60.1012 | 3.33177 | 0.233596 |
| 191 | Area Entolateralis Anterior R | 558 | 61.3098 | 3.52955 | 0.510408 |
| 192 | Area Entolateralis Anterior L | 647 | 71.0886 | 3.36227 | 0.5853 |
| 193 | Area Coronalis Posterior Lateralis R | 271 | 29.7759 | 3.39034 | 0.188925 |
| 194 | Area Coronalis Posterior Lateralis L | 195 | 21.4255 | 3.38403 | 0.214025 |
| 195 | Area Fissurae Ectolateralis Posterior L | 761 | 83.6142 | 3.32462 | 0.253919 |
| 196 | Area Fissurae Ectolateralis Posterior R | 731 | 80.318 | 3.41685 | 0.258023 |
| 197 | Area Fissurea Retrospenialis R | 2719 | 298.748 | 3.46326 | 0.270459 |
| 198 | Area Fissurea Retrospenialis L | 2731 | 300.066 | 3.40012 | 0.263005 |
| 199 | Area Limbica Posterior Ventralis II R | 685 | 75.2638 | 3.50079 | 0.619913 |
| 200 | Area Limbica Posterior Ventralis II L | 618 | 67.9022 | 3.57288 | 0.396628 |
| 201 | Area Suprasylvian Ventralis R | 1391 | 152.835 | 3.4706 | 0.436353 |
| 202 | Area Suprasylvian Ventralis L | 1169 | 128.443 | 3.59801 | 0.463106 |
| 203 | Area Fissurae Entolateralis Pars Anterior R | 76 | 8.35044 | 3.74624 | 0.531476 |
| 204 | Area Fissurae Entolateralis Pars Anterior L | 54 | 5.9332 | 4.03591 | 0.560814 |
| 205 | Area Pregenualis I R | 888 | 97.5682 | 3.1558 | 0.367322 |
| 206 | Area Pregenualis I L | 800 | 87.8993 | 3.19368 | 0.306035 |
| 207 | Area Subprorealis Lateralis II R | 331 | 36.3683 | 3.262 | 0.30456 |
| 208 | Area Subprorealis Lateralis II L | 307 | 33.7314 | 3.47042 | 0.261577 |
| 209 | Area Prorealis Lateralis II R | 467 | 51.3112 | 3.46531 | 0.408026 |
| 210 | Area Prorealis Lateralis II L | 388 | 42.6312 | 3.37832 | 0.413469 |
| 211 | Area Composita Sigmoidea Lateralis R | 263 | 28.8969 | 3.41208 | 0.229843 |
| 212 | Area Composita Sigmoidea Lateralis L | 306 | 33.6215 | 3.51947 | 0.252802 |
| 213 | Area Recurrens Lateralis R | 414 | 45.4879 | 3.38264 | 0.508834 |
| 214 | Area Recurrens Lateralis L | 530 | 58.2333 | 3.58759 | 0.416733 |
| 215 | Area Pararecurrens Anterior R | 1574 | 172.942 | 3.3722 | 0.286326 |
| 216 | Area Pararecurrens Anterior L | 1630 | 179.095 | 3.35344 | 0.296632 |
| 217 | Area Recurrens Medialis R | 766 | 84.1636 | 3.40257 | 0.491639 |
| 218 | Area Recurrens Medialis L | 708 | 77.7909 | 3.49965 | 0.539122 |
| 219 | Area Recurrens Ventralis Medialis R | 284 | 31.2043 | 3.50646 | 0.448957 |
| 220 | Area Recurrens Ventralis Lateralis L | 454 | 49.8829 | 3.83117 | 0.592043 |
| 221 | Area Genualis II R | 286 | 31.424 | 3.439 | 0.203976 |
| 222 | Area Genualis II L | 263 | 28.8969 | 3.41846 | 0.242862 |
| 223 | Area Paraorbitalis Dorsalis R | 1055 | 115.917 | 3.59436 | 0.259201 |
| 224 | Area Paraorbitalis Dorsalis L | 945 | 103.831 | 3.6088 | 0.235574 |
| 225 | Area Precruciata Lateralis R | 691 | 75.923 | 3.3965 | 0.242316 |
| 226 | Area Precruciata Lateralis L | 718 | 78.8896 | 3.46217 | 0.206877 |

**Table S4. Basic information of the experimental dogs.**

| **Subject** | **Breed** | **Sex** | **Age(years)** | **Weight(kg)** |
| --- | --- | --- | --- | --- |
| 1 | Kunming dog | M | 3 | 30 |
| 2 | Kunming dog | M | 7 | 30 |
| 3 | Kunming dog | M | 5 | 34 |
| 4 | Kunming dog | M | 8 | 32 |
